# A Bayesian active learning platform for scalable combination drug screens

**DOI:** 10.1101/2023.12.18.572245

**Authors:** Christopher Tosh, Mauricio Tec, Jessica White, Jeffrey F. Quinn, Glorymar Ibanez Sanchez, Paul Calder, Andrew L. Kung, Filemon S. Dela Cruz, Wesley Tansey

## Abstract

Large-scale combination drug screens are generally considered intractable due to the immense number of possible combinations. Existing approaches use ad hoc fixed experimental designs then train machine learning models to impute novel combinations. Here we propose BATCHIE, an orthogonal approach that conducts experiments dynamically in batches. BATCHIE uses information theory and probabilistic modeling to design each batch to be maximally informative based on the results of previous experiments. On retrospective experiments from previous large-scale screens, BATCHIE designs rapidly discover highly effective and synergistic combinations. To validate BATCHIE prospectively, we conducted a combination screen on a collection of pediatric cancer cell lines using a 206 drug library. After exploring only 4% of the 1.4M possible experiments, the BATCHIE model was highly accurate at predicting novel combinations and detecting synergies. Further, the model identified a panel of top combinations for Ewing sarcomas, all of which were experimentally confirmed to be effective, including the rational and translatable top hit of PARP plus topoisomerase I inhibition. These results demonstrate that adaptive experiments can enable large-scale unbiased combination drug screens with a relatively small number of experiments, thereby powering a new wave of combination drug discoveries. BATCHIE is open source and publicly available (https://github.com/tansey-lab/batchie).

## Main

Single-agent treatment interventions in cancers, viruses, and bacterial infections impose evolutionary selective pressures that can lead to therapeutic resistance and poor outcomes for patients. Combination therapies have the ability to constrain multiple potential avenues of evolutionary escape and thus reduce the likelihood of treatment resistance. Consequently, rational combination therapies have long formed the basis for rapidly evolving pathogens like HIV ^1^ and are increasingly seen as the future of antibiotics ^2^ and cancer therapies ^3,4^.

Screening for novel effective drug combinations presents the singular challenge of scale. The number of possible experiments in a combination screen grows at a rate of *n* × *m*^*d*^ ×*t*^*d*^ for *n* conditions, *m* drugs, *t* doses, and *d*-way combinations. For instance, a single-agent screen of 100 drugs and 50 cell lines at 5 doses would only be 25K experiments whereas a pairwise drug screen on the same libraries would require 12.5M experiments. Given the rapid growth of the experimental design space, even the most efficient high-throughput screening team would struggle to conduct an exhaustive pairwise combination screen of a modest-sized drug library over a modest number of doses and cell lines. Conducting such a screen is currently a substantial undertaking requiring large funding, detailed planning, advanced equipment, and several years of experiments. Thus, even the largest published combination screens have been restricted to libraries of less than 120 drugs ^5,6,7^.

The intractability of combination drug screens has led to the development of machine learning methods for predicting novel drug combinations^8,9,10,11,12^. The goal of such modeling is to use the predictive model to simulate experiments *in silico* and filter the list of combinations down to a set of top hits to be validated *in vitro*. However, predictive models are fundamentally limited by the data on which they are trained. Small libraries, which allow for exhaustive enumeration of the combination landscape, are limited to discoveries involving those drugs in the library. If the library is too small, it simply may not contain any useful candidate combinations and thus machine learning models will extract no meaningful signals (Fig. 1a). Larger libraries may contain useful combinations, but if a screen is performed on a random or fixed-design subset, then it is unlikely to generate observations of the most surprising and informative combinations. Models fit to pre-designed observations are likely to then have poor accuracy and may be unable to confidently pinpoint the maximally useful combinations (Fig. 1b). Thus while a number of sophisticated models have been proposed, their success as discovery tools for novel rational combinations has been limited.

**Figure 1:**
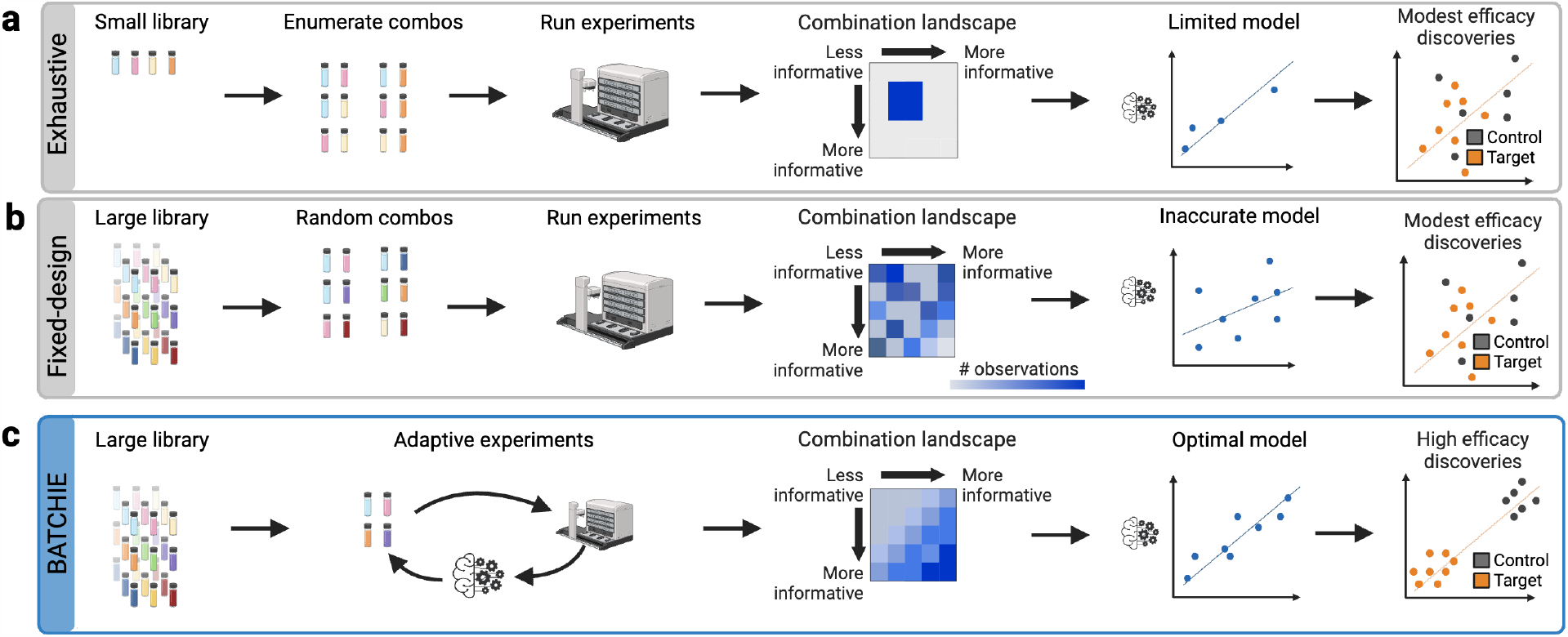
Benefits of adaptive experimental design. **(a**,**b)** Traditional approaches to combination studies entail exhaustive enumeration of a small library or random subsampling of a large library. In either case, the resulting models generally are not powerful enough to find clinically-relevant discoveries. **(c)** Adaptive experimental design allows exploration of a large library. By focusing on the most informative experiments, one can create a powerful model that is capable of finding clinically-relevant discoveries.

We sought to overcome both the wet lab scalability and predictive modeling challenges of combination screens by rethinking the experimental design approach. Rather than performing a fixed-design experiment and fitting a model post-hoc, we instead drew on the long history of Bayesian optimal sequential experimental design ^13^. In the sequential setup, experiments are conducted in small batches where each batch is designed adaptively based on the results of the previous batches. When designs are driven by a machine learning model that aims to acquire the most informative training data in each batch, the sequential experimental design task is known as active learning ^14^. Active learning has experienced a recent burst in interest in the drug discovery literature ^15,16,17,18^ where the objective is to design *de novo* molecules aimed at better docking on a target. We are aware of only one recent method, RECOVER ^19^, that has attempted to integrate adaptive experimentation and combination screening. RECOVER uses a multi-armed bandit algorithm to search for top synergistic hits for a single cell line at each iteration. Unfortunately, the RECOVER bandit approach provides no clear way to scale to large sample libraries, provides no theoretical guarantees that designs are optimal, and can only target single cell line synergy which is generally rare ^6^ and does not typically characterize successful combination therapies in the clinic ^20,21,22,23^. Thus, while using active learning to identify novel molecules has seen robust investigation, its application to discover rational combinations from large libraries of existing drugs has not been thoroughly explored.

In this paper, we introduce Bayesian Active Treatment Combination Hunting via Iterative Experimentation (BATCHIE), a framework for orchestrating large-scale combination drug screens through sequential experimental design. BATCHIE uses a novel optimal Bayesian active learning ^24,25^ strategy to design sequential experiments. These sequential designs are theoretically near-optimal (see Supplement for theory and proofs) and guarantee that BATCHIE designs are highly efficient. Pragmatically, BATCHIE enables sequential experimental designs that will best improve any user-provided probabilistic (Bayesian) model. Thus, while we implemented an initial model for our experiments, BATCHIE can alternatively be paired with any existing and future Bayesian machine learning method for combination drug response. The end result of a BATCHIE screen is a maximally informative dataset and an optimal predictive model that enables the discovery of more effective combinations than in a fixed design (Fig. 1c).

We validate the empirical performance of BATCHIE using retrospective simulations and through a prospective study. We first use data from large-scale combination screens ^7,5,6^ to retrospectively simulate adaptive screens. BATCHIE consistently outperforms fixed designs in these simulations and better prioritizes effective combinations as top hits. We then implement BATCHIE in a drug screening facility and use it to conduct a combination drug screen across a 206 drug library over 16 cancer cell lines, focusing on pediatric sarcomas. The BATCHIE screen generates a model with near-optimal predictive accuracy on random novel combinations. Further, we use the model to prioritize ten combinations to validate experimentally for a panel of Ewing sarcoma lines; all ten combinations achieve a high therapeutic index score. The top identified hit corresponds to a PARP inhibitor (talazoparib) plus a topoisomerase I inhibitor (topotecan), a biologically rational combination and the subject of two of the five NCI-supported Phase II combination therapy clinical trials currently underway. BATCHIE is readily available as open source, along with accompanying tutorials.

## Results

### Adaptive experimental design for combination drug screens

BATCHIE uses an active learning algorithm to choose the most informative data to collect in each sequential batch of experiments (Fig. 2). In the initial batch, BATCHIE uses a design of experiments approach ^26^ to cover the drug and cell line space efficiently (Fig. 2a). The initial batch is then run (Fig. 2b) and used to train a Bayesian predictive model that estimates a distribution over drug combination responses for each cell line (Fig. 2c). For subsequent batches, BATCHIE uses the model’s posterior distribution to simulate plausible outcomes of candidate combination experiments along with how they would change the model (Fig. 2d). BATCHIE then measures how much each experiment is expected to reduce the posterior uncertainty over the drug responses (Fig. 2e) and uses a submodular approach to design a maximally informative batch (Fig. 2f). After designing an optimal batch, the batch is run, the model is updated with the new results, and the next optimal batch is constructed. When the exploratory budget runs out or the model converges to concentrated posterior, the active learning loop ends. The optimally trained model is then used to predict effective novel combinations that are prioritized for experimental validation (Fig. 2g).

**Figure 2:**
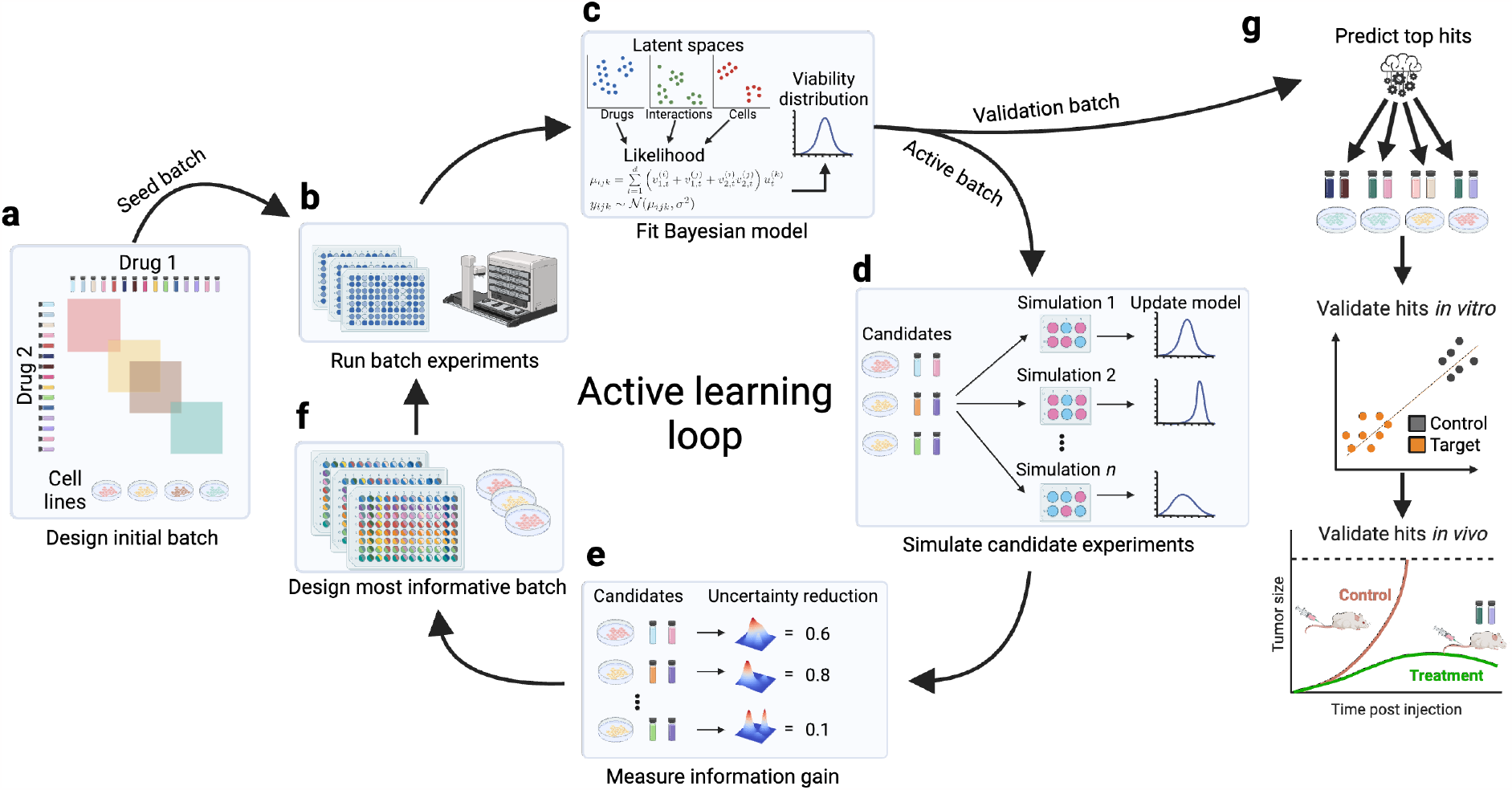
BATCHIE overview. **(a)** The BATCHIE workflow begins by specifying a cell line library, a drug library, and an initial ‘seed batch’ of plates to cover every cell line and drug with at least one experiment. **(b)** Selected plates are assembled, run, measured, and post-processed to obtain viability scores. Quality control (QC) checks filter out problematic wells. **(c)** A Bayesian tensor factorization model is fit to the current data. Posterior samples are drawn via MCMC. **(d)** The joint distributions of candidate experiments are estimated using the current set of posterior samples. **(e)** The active learning criterion is applied to the joint distribution estimates to score the utility of individual experiments. **(f)** The scores of individual experiments are aggregated to define the most informative batch of experiments to run next, possibly subject to design constraints. **(g)** After terminating the active learning loop, the most recently fitted Bayesian model is used to predict top hits for individual combinations. These top hits can then be validated *in vitro* and, potentially, *in vivo*.

BATCHIE is compatible with any Bayesian model capable of modeling combination drug screen data. There have been many models developed for modeling this type of data ^27,28,11,29^. Integrating any of these models into BATCHIE would be possible by reformulating them as fully Bayesian models capable of quantifying posterior uncertainty. In our implementation, we use a hierarchical Bayesian tensor factorization model (Fig. 2c). The model contains embeddings for each cell line and each drug-dose, as well as embeddings that capture the effects of drug interactions. The BATCHIE model assumes that the response of a combination of drugs on a cell line can be decomposed into the individual effects of the drugs and an interaction term. Specifically, the model posits an embedding *u*^(*k*)^ ∈ ℝ^*d*^ for each cell line *k* and embeddings 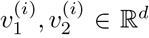 for each drug-dose *i*, which capture the individual and interaction effects of the drug-dose, respectively.

When drug-doses *i* and *j* are applied to cell line *k*, the logit-transformed viability is assumed to be normally distributed with mean *μ*_*ijk*_ satisfying

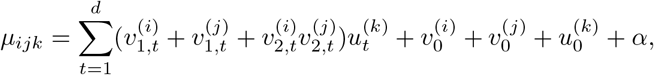

where 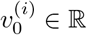 is a drug-dose specific offset, 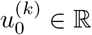 is a cell line specific offset, and *α* ∈ ℝ. The variance of the normal distribution is assumed to be a global paremeter. Hierarchical priors are placed on top of the embeddings, offsets, and variance to automatically adapt to the complexity of the data ^30^. The priors are chosen to be conditionally conjugate with the likelihoods, allowing the entire model to sampled efficiently using blocked Gibbs sampling (see Methods for modeling details).

To make BATCHIE practical to implement in a high-throughput screening facility, we consider pools of experiments in the form of microwell plates (Supplementary Fig. 3a). Each plate is formed by combining a single cell line with a ‘row’ plate and a ‘column’ plate. Here, a row plate is an *n* ×*m* plate in which every well of a particular row contains the same drug at the same dose. Similarly, a column plate is of the same size, but constructed so that each column contains the same drug at the same dose. When a row plate and a column plate are overlaid, the resulting combination plate contains all possible combinations of their constituent drug-doses. Row and column plates are constructed with high and low control wells so that all associated drug-doses are measured singly and viabilities can be computed. In this way, the number of plates that need to be stamped scales linearly with the drug library size, as opposed to quadratically like the number of possible combinations (see Methods).

### Validation of BATCHIE on existing combination datasets

The core goal of BATCHIE is to reduce the number of experiments needed to make useful discoveries. To benchmark the effectiveness of BATCHIE in pursuit of this goal, we conducted retrospective simulations that compared the performance of a BATCHIE-trained model after a small number (≤ 15) of rounds against (a) a model that was trained on the entire dataset and (b) a model that was trained on a random subset of the dataset matching the size and constraints of BATCHIE’s.

We benchmarked BATCHIE using three large, publicly-available combination drug screen datasets: the NCI ALMANAC study ^5^ (ALMANAC), the Genomics of Drug Sensitivity in Cancer combination screen ^6^ (GDSC^2^), and Merck’s unbiased combination drug screen ^7^ (MERCK). The three datasets differed in cell line library size (60 for ALMANAC, 126 for GDSC^2^, 39 for MERCK) and drug library size (104 for ALMANAC, 66 for GDSC^2^, 38 for MERCK). The ALMANAC screen covered the NCI-60 panel of cell lines ^31^, spanning leukemia, lung, colon, central nervous system, melanoma, ovarian, renal, prostate, and breast cancers. The GDSC^2^ screen covered breast, colon and pancreatic cancer cell lines. The MERCK screen covered a panel of lung, ovarian, melanoma, colon, breast, and prostate cancer cell lines. All three screens used fixed experimental designs but differed in the subset of the combination space to explore. Consequently, the overall sparsity pattern varies substantially between datasets (Fig. 3(b,c)).

**Figure 3:**
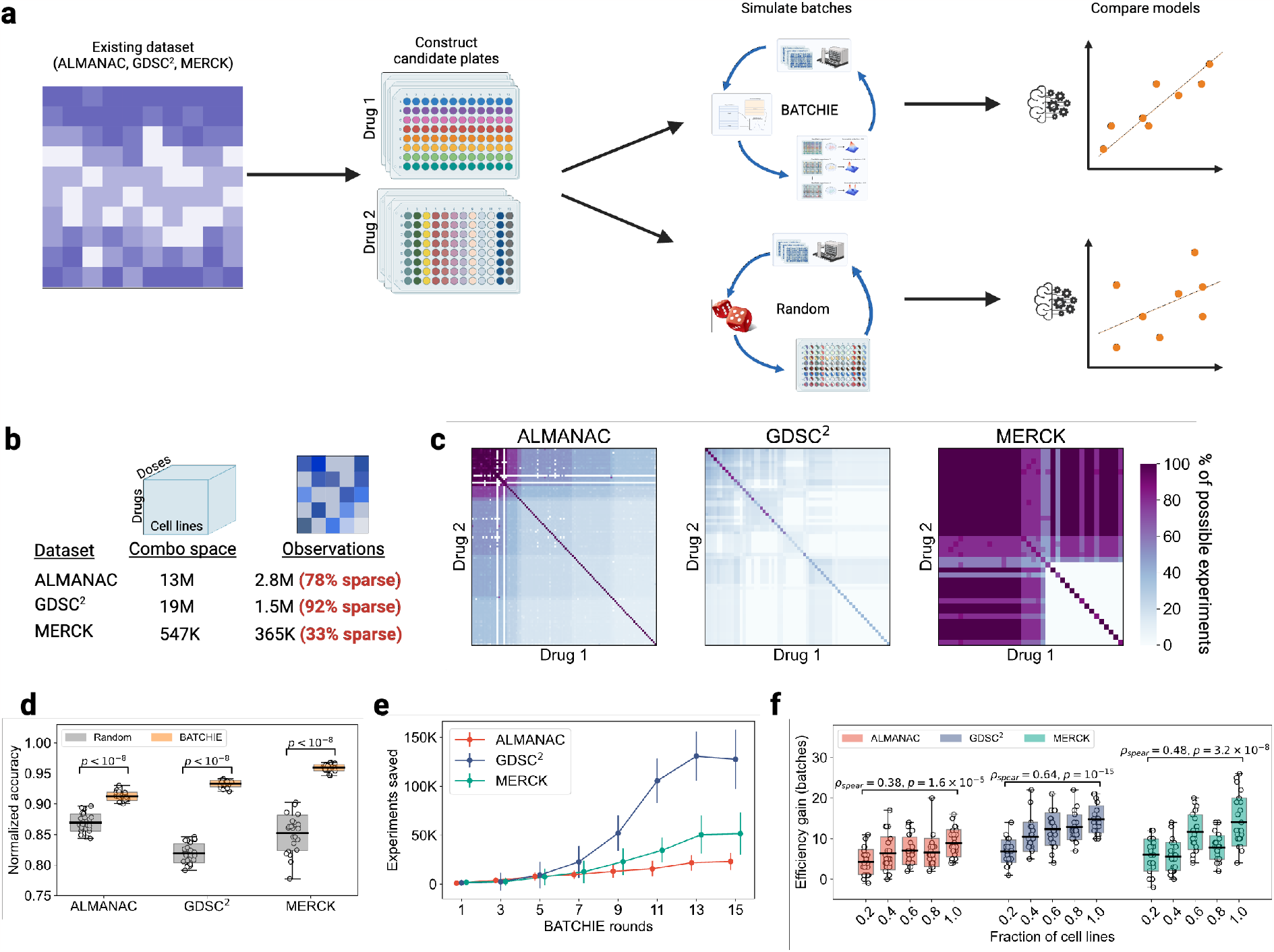
Retrospective study design and results. **(a)** Retrospective studies are conducted by processing an existing dataset into candidate row/column plates of the kind considered in this work and simulating the choices made by both the Random and BATCHIE data collection procedures. After data collection, the models are trained on the data collected by each of the procedures and evaluated for accuracy. **(b)** Statistics for the datasets used in our retrospective studies. **(c)** Heatmaps showing the fraction of possible experiments observed for each drug combination. **(d)** BATCHIE outperforms Random when both are evaluated after 15 rounds of data collection; p-values derived from a Mann-Whitney U-Test. **(e)** The number of additional experiments needed for Random to achieve comparable performance to BATCHIE grows with the number of BATCHIE rounds across all datasets. Error bars denote standard deviations. **(f)** The number of additional batches needed for Random to achieve comparable performance to BATCHIE grows with the size of the cell line library; *ρ*_spear_ is Spearman’s rho for nonparametric rank correlation. For boxplots in **(d**,**f)**, center lines denote means, box limits denote standard deviations, and whiskers denote extremal values.

To simulate the behaviour of BATCHIE in a realistic adaptive data collection screen, the existing datasets were divided into simulated plates (Fig. 3a). Plates were designed to replicate the statistics of the plates used in our prospective study (Methods). The data collection processes (BATCHIE and Random) were given batch constraints that approximately 10% of cell lines be selected at every round and three combination plates be selected per chosen cell line. We stopped the Random and BATCHIE screens after 15 batches, the same as in our prospective study. We randomly held out 10% of all experiments as a test set to evaluate predictive performance of the resulting models. Model performance was compared relative to a model trained on the full set of experiments conducted, not including the test set. We repeated the simulations 25 times per dataset, with different randomly-organized row and column plates for stamping.

At the end of 15 batches, the Random and BATCHIE models observed 1.7%-20.4% of the full training dataset (1.66%-1.7% for ALMANAC, 10.3%-11.1% for GDSC^2^, and 19.3% -20.4% for MERCK). Total observed percentages varied due to the difference in the original screen designs. Across the three datasets, BATCHIE produced models with holdout *R*^2^ accuracy within 5-7% of the models that were fit using all available training data and significantly outperformed the models trained on data collected by the Random strategy with the same number of rounds (Fig. 3d).

We also tracked the number of experiments that the Random strategy would need to perform in order to achieve comparable performance to BATCHIE. We observed that the number of experiments saved grew as a function of the number of BATCHIE rounds, with the number of experiments saved by round 15 numbering in the 10s of thousands for the ALMANAC and MERCK datasets to over 100 thousand for the GDSC^2^ dataset (Fig. 3e). When we looked at the excess number of experiments needed to achieve a certain normalized accuracy, BATCHIE shows an exponential speedup as the model accuracy threshold increases (Supplementary Fig. 1c). When measured by the number of batches needed for Random to achieve BATCHIE-level performance, we again saw a similar exponential trend, but the scale is more consistent across datasets (Supplementary Fig. 1b). Similar trends were seen when measuring the number of excess batches required for the Random model to reach the equivalent performance of the BATCHIE model at each round (Supplementary Fig. 1d).

To test how well BATCHIE scales with the size of the experimental landscape, we simulated smaller experimental spaces by restricting each dataset to a random subsample of 20%, 40%, 60%, or 80% of cell lines. For all datasets, we observed a strong positive correlation (*ρ*_spear_ = 0.38, 0.64, 0.48, max(*p*) = 1.6 ×10^−5^) between the size of the experimental landscape and the improvement offered by BATCHIE over Random (Fig. 3f). These results indicate that BATCHIE screens become increasingly more efficient as the overall landscape becomes larger.

While these results confirm that BATCHIE produces accurate models with very few experiments, average predictive accuracy alone does not ensure that the model will be able to identify highly effective combinations. This is because desirable properties for treatments, such as high therapeutic index (TI)^32^, often correspond to extremal points, which by definition are not average. To evaluate the ability of BATCHIE to discover effective combinations, we used the BATCHIE-trained model to estimate the TI of all drug combinations using all pairs of cell lines as target and control. We chose the top 20 predictions and calculated their average observed TI. For all of the datasets, the BATCHIE-trained model selections had significantly higher TI (max(*p*) = 0.001) than those selected by the Random-trained model (Supplementary Fig. 1e). We also observed that BATCHIE was better at identifying high TI combos in terms of area under the curve (AUC) of the receiver operating characteristic (ROC) curve (Supplementary Fig. 1f).

We also implemented BATCHIE in the setting where the goal is solely to model synergy or antagonism (Supplementary Fig. 2a). We adapted our retrospective setup so that all single drug data was made available to the models before data collection and models only needed to predict Bliss scores. We also implemented a synergy-only Bayesian hierarchical model, making this benchmark an example of the flexibility of BATCHIE to adapt to new designs and alternative models (Methods). Synergy detection is a challenging task as synergies are rare ^6^. In the three benchmark datasets, only 0.12%-1.19% of combinations are synergistic and 0.09%-0.82% are antagonistic (Supplementary Fig. 2b). We again observed significant gains (max(*p*) = 10^−7^) in predictive accuracy on held out data when comparing BATCHIE to the Random design baseline (Supplementary Fig. 2c). We also confirmed that the BATCHIE-trained model is better at detecting top synergy hits and top antagonism hits (Supplementary Fig. 2d,e). Across all three datasets, the BATCHIE model saves approximately 25K experiments compared to the Random strategy by batch 15 with similar upward trends in each dataset (Supplementary Fig. 2f).

### A prospective pediatric sarcoma study with BATCHIE

Current treatments for many pediatric sarcomas have unacceptably high failure rates, particularly for metastatic and recurrent presentations^33^. Ewing sarcoma ^34^ (EWS), Rhabdomyosarcoma (RMS), and osteosarcoma ^35^ (OST) are amongst the most common pediatric sarcomas in need of improved treaments. We conducted a large-scale combination drug screen on pediatric cell lines, with a focus on pediatric sarcomas. The study covered 16 cell lines: 5 Ewing sarcoma lines (A673, MSKEWS-38338, MSKEWS-66647, SKNEP, TC-71), 5 osteosarcoma lines (MG-63, MSKOST-11890, SAOS-2, SJSA-1 U2OS), 1 rhabdomyosarcoma line (MSKRMS-12808), 3 other cancer cell lines (Kelly, MDA-MB-231, Wit49), and 2 non-cancer lines (RPE, BJ)(Fig. 4a). The non-cancer lines were included in the study to allow us to evaluate meaningful notions of TI, as high activity in a target cell line alone does not necessarily translate to clinical utility ^32^. Mathematically, we defined therapeutic index to be the minimum viability of the control cell lines (i.e. RPE and BJ) minus the median viability of the target cell lines (e.g. all EWS lines). Thus, a high TI indicates a drug has high activity in the target lines and not the control lines.

**Figure 4:**
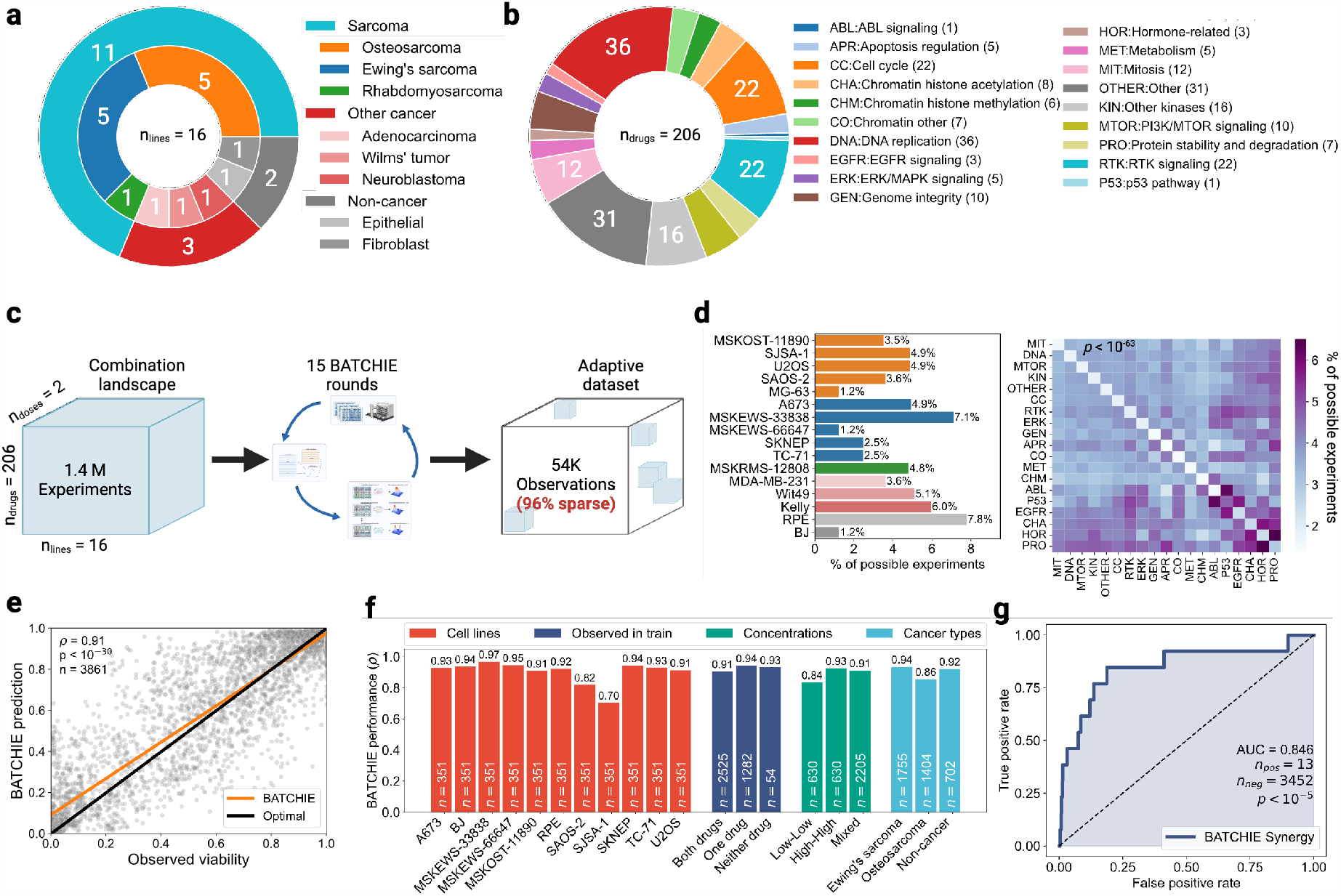
Prospective study design and random validation results. **(a)** Prospective study cell line library broken down by type. **(b)** Prospective study drug library broken down by mechanism of action. **(c)** Overview of prospective study: after 15 rounds of BATCHIE data collection, approximately 4% of possible combinations were observed. **(d)** Observation breakdown by cell line and drug mechanism of action; Pearson’s chi-square test *p <* 10^−63^ where the null hypothesis is that experiments were sampled uniformly at random. Colors in the left panel match the corresponding type colors from panel **(a). (e)** Scatter plot of mean BATCHIE predictions v.s. observed viabilities on random validation data. Orange line indicates regression of predictions onto observations, and black line denotes the identity line. **(f)** Pearson correlation between predictions and observed viabilities, broken down by cell line, observation status, concentration and cancer type. All associated p-values satisfy *p <* 10^−25^. **(g)** ROC curve for synergy identification on random validation data. Synergy is defined here as having an observed Bliss score larger than 0.25. *ρ* in **(e**,**f)** is Pearson’s *ρ* correlation coefficient.

The drug library for consisted of 206 drugs, both FDA approved and investigational. In order to ensure adequate coverage of complementary mechanisms, the drug library was chosen to span a variety of targets (Fig. 4b). We ensured that the library included inhibitors of the most theoretically promising targets for Ewing sarcoma and osteosarcoma such as PARP, CDK4/6, and CD99^36,37^ as well as commonly-used chemotherapy drugs. Each drug was tested at two concentrations: 0.1*μ*M (the *low* dose) and 1*μ*M (the *high* dose). All drugs on a single row or column plate were plated at the same concentration, allowing each drug combination to occur at 4 dose combinations (low-low, low-high, high-low, and high-high). Each active learning batch consisted of choosing 3 cell lines and 3 combination plates for each chosen cell line, resulting in 9 plates in total per batch.

The screen was divided into two phases (Supplementary Fig. 3b). In phase I, we started with a focus on OST, using 4 OST lines and a single line each for EWS and RMS. We also included the three non-sarcoma cancer cell lines and used a single control line (RPE). After 10 rounds of BATCHIE, we observed the model was converging, as measured by a diminishing improvement of cross-validation accuracy (Supplementary Fig. 3d). We therefore paused the screen and assessed the model’s predictions on the three sarcoma types. We observed over three-times as many combinations predicted to have high therapeutic index in the EWS line compared to either the OST lines or the RMS line (Supplementary Fig. 3f). As a preliminary test, we selected three drugs (Clofarabine, Eltanexor, and Talazoparib) each of which was predicted to have high TI on the EWS lines at the low concentration when paired with another of the three at the low concentration. We experimentally validated that the three hits had TI in the 90th percentile of all observed combinations through round 10 (Supplementary Fig. 3g).

Since the phase I results suggested that our drug library may contain effective combinations particularly for EWS, in phase II we focused the screen on EWS. We added four EWS lines and removed the other non-sarcoma cancer cell lines. We also added an additional control line (BJ) to increase the robustness of our control set in TI estimates; a fifth OST line (MG-63) was also added. After the single initial seed batch for the newly added cell lines, the BATCHIE model quickly learned that the EWS lines were highly related. Two of the four EWS lines introduced in phase II were significantly positively correlated with the phase I EWS line and none of the other phase II lines (Supplementary Fig. 3h). Phase II proceeded for five rounds of BATCHIE until we again observed the model’s cross-validation accuracy converging (Supplementary Fig. 3e). We then ended the adaptive portion of the screen and moved to the validation phase.

In total, we ran BATCHIE for 15 rounds of data collection, generating approximately 54K unique cell line, drug-dose pair combinations. As the full design space was approximately 1.4M possible experiments, BATCHIE explored approximately 4% of the total landscape (Fig. 4c). This dataset exhibited significant variability (Pearson’s chi-square test, *p <* 10^−63^) in the sampling frequencies with respect to both cell lines and drug classes (Fig. 4d), indicating that certain cell lines and drugs were more informative than others to the model. The correlations among predictions made by the final model reveal that it clearly learned to separate the EWS cell lines from the OST and control lines (Supplementary Fig. 4f). The OST lines were less homogeneous in the model predictions, as expected due to OST being a disease marked by chromothripsis which leads to potentially hundreds of random chromosomal translocations and thus high genomic heterogeneity compared to relatively stable EWS cancers ^38^.

At the drug level, the predictions of the model correlated strongly amongst drugs of similar mechanisms. Hierarchical clustering of predictions (Supplementary Fig. 4d) grouped targeted therapies (top left) distinctly from more broad-spectrum cytotoxic agents (bottom right). Among the targeted therapies, the predictions clearly distinguished androgen receptor inhibitors and MEK inhibitors. Within the broad-spectrum cytotic agents, the predictions identify two clades. The first of these contained taxanes and vinca alkaloids, drugs whose mechanisms of action (MoAs) all target mitosis. The majority of the drugs in the other clade, containing anthracycline topoisomerase inhibitors, nucleoside metabolic inhibitors, selective inhibitors of nuclear transport, and antihelmintics, were predominantly focused on targeting DNA replication. Across both groups, we observed that the finer-grained drug class structure was largely respected by the hierarchical clustering. In an orthogonal analysis, we projected high dose predictions to 2d using t-SNE and again observed drugs clustering by MoA with the dominant spatial axis corresponded to drug potency (Supplementary Fig. 4e).

### Validation of BATCHIE predictions on random unseen combinations

To evaluate the accuracy of the BATCHIE-trained model, we constructed a test set of randomly selected combination plates from the remaining unexplored experimental space. We randomly selected from among the experimental plates that had no overlap in any cell line, drug-dose pair combinations with the BATCHIE-collected data. One plate was selected for every cell line that was under investigation in phase II. One of the test plates (cell line MG-63) did not pass the quality control checks (Methods) and was excluded from performance measurement.

The BATCHIE model predictions were highly accurate on the unseen validation data (Fig. 4e, Pearson’s *ρ*=0.91, *p <* 10^−30^). The accuracy of the model was robust to stratification by cell line, previous observation status, drug concentration, and cancer type (Fig. 4f). Indeed, for 10 of the 11 cell lines, Pearson’s *ρ* was above 0.82, and was above 0.91 for 9 of the 11. The previous observation status of the drug on the cell line also appeared not to make a large difference, as the correlation remained above 0.91 regardless of whether one, both, or neither of the drugs had previously been observed on the chosen cell lines. We observed that there was dip in performance to *ρ* = 0.84 when restricting to the low-low doses. This decrease in performance however is confounded since by chance half of the low-low validation set was on the SJSA-1 cell line, the cell line on which BATCHIE performed worst. SJSA-1 is an OST line that shares little predictive similarity to the other OST lines (Supplementary Fig. 4f), which may explain its relative difficulty in the test set.

Similar to previous combination cell line screens^6,7^, we found very few synergistic combinations in the random validation set. Of the 3465 observed combinations, only 13 (0.004%) resulted in a Bliss score larger than 0.25. Nevertheless, the BATCHIE model accurately identified these combinations (Fig. 4g), achieving an AUC of the ROC curve of 0.846 (*p <* 10^−5^ by permutation test over 100K random permutations).

### BATCHIE discovers rational drug combinations for Ewing sarcoma

To validate BATCHIE’s ability to discover effective combinations, we used the BATCHIE model to identify combinations with an expected high therapeutic index. We ranked the drug combinations by looking at their predicted viabilities at low concentrations and taking the difference between the minimum predicted viability on the two non-cancer lines and the median viability on the OST and EWS lines. We found that no combinations were predicted to be robust across all OST lines such that the TI would be high. However, we did identify several drug combinations that were predicted to have high TI across a range of EWS lines. We selected 10 of the top-ranked candidates and collected a fine-grained dose-response matrix spanning 0.006nM - 400nM via four-fold dilution (Methods), which included the low concentration combination (Fig. 5a). We also selected 13 negative control combinations that exhibited a large predicted differential effect for at least two cell lines but were not predicted to have a high TI over the five EWS lines.

**Figure 5:**
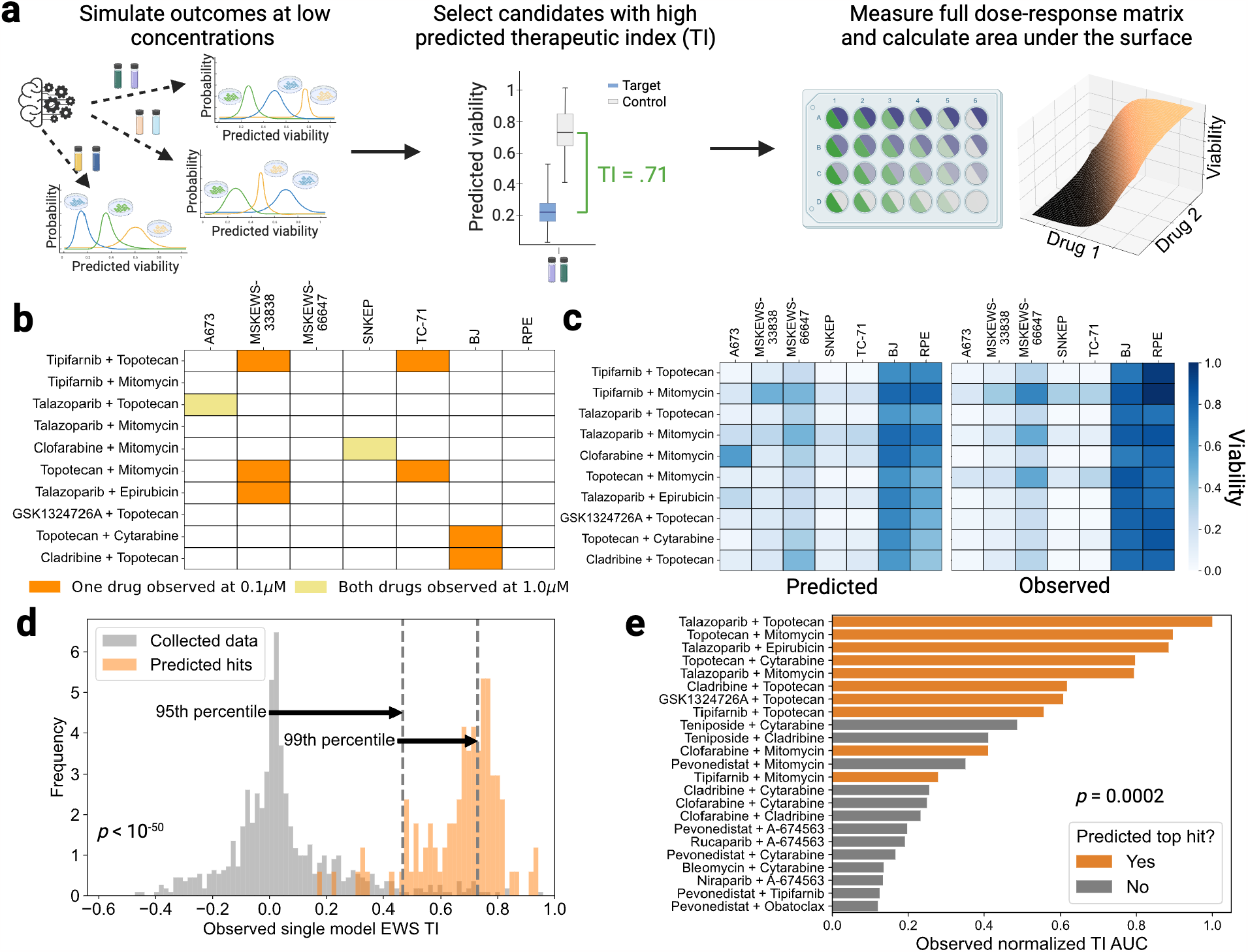
Prospective study validation of top hits. **(a)** Pipeline for top hit selection and validation in the prospective study. The model fit on BATCHIE-collected data is used to simulate outcomes for drug combinations at the low (0.1*μ*M) concentration. Those simulated values are collected into confidence-rated predictions of TI values. The top hits are then selected for further *in vitro* experimentation, collecting observations over the full dose-response matrix. **(b)** Observation status of selected top hits in BATCHIE-collected data. **(c)** Mean predictions and observed viabilities for selected top hits at 0.1*μ*M concentration. Pearson’s correlation *ρ >* 0.92 with p-value *p <* 10^−28^. **(d)** Histogram of observed single model EWS TI values for combinations at 0.1*μ*M concentration in BATCHIE-collected data (gray, *n* = 659) and top predicted hits (orange, *n* = 100). Percentiles are drawn with respect to BATCHIE-collected data. **(e)** Observed AUCs with respect to the TI dose-response surface for selected top hits and not top hits. p-values for **(d**,**e)** computed using a Mann-Whitney U Test.

The top hit selections for EWS had not been observed in the training data at the low concentration for any cell line, and the general observation pattern was sparse (Fig. 5b). Nevertheless, at the low-concentration there was a strong correlation (Pearson’s *ρ* = 0.92, *p <* 10^−28^) between the predicted viabilities and the observed viabilities (Fig. 5c). This accuracy at the viability level translated to the observed TIs being large, with the median TI score in the top hit predictions being higher than the 98th percentile of observed TI scores in the 54K training observations (Fig. 5d).

Although the top hits were chosen solely on the basis of their predicted TI at a specific concentration, we found that they generally exhibited high TI across a wide range of dose pairs. After computing the TI for each entry of the dose-response matrix, we calculated the area under the TI surface (3d curve) and observed that the top hits exhibited significantly higher AUC values than the reference combinations (*p* = 0.0018, Fig. 5e).

The selected combinations exhibit biologically plausible rationales in Ewing sarcomas. Ewing sarcomas frequently exhibit EWS-FLI1 genomic fusions, which tend to interact with the DNA repair protein PARP-1^39^. Talazoparib is a PARP inhibitor, and combining PARP inhibitors with treatments that induce DNA damage has previously been shown to lead to cytotoxicity in preclinical Ewing sarcoma studies^40^. These observations have led to clinical trials combining PARP inhibitors with irinotecan (a topoisomerase 1 inhibitor) and temozolomide (an alkylating agent) for Ewing sarcoma and related cancers ^41,42,43^. Topotecan is a topoisomerase 1 inhibitor, mitomycin is an alkylating agent that cross-links complementary DNA strands, and epirubicin is an anthracycline that blocks the action of topoisomerase 2. Thus, these selected combinations with talazoparib may facilitate the utility of PARP inhibition by accelerating DNA damage.

Of the remaining drugs, cytarabine and GSK1324726A, may act by reducing the overall abundance of the EWS-FLI1. This has been directly shown for cytarabine *in vitro* ^44^. On the other hand, GSK1324726A is a BET bromadine inhibitor, and it has been shown that BET bromadine proteins are required for EWS-FLI1 transcription ^45,46^. Clofarabine and cladribine are deamination resistant analogues of deoxyadenosine, and as such interfere with DNA synthesis through incorporation into DNA ^47,48^. Both drugs have been shown to inhibit Ewing sarcoma growth *in vitro* by binding to C99, which is overexpressed in Ewing sarcoma ^49^. EWS-FLI1 also suppresses SPRY1, a downstream feedback inhibitor of certain Ras-activating receptors ^50^. Tipifarnib is a farnesyltransferase inhibitor that interferes with the Ras signaling pathway ^51^. Combining tipifarnib with drugs that induce DNA damage can be seen as targeting two separate downstream effects of EWS-FLI1.

To evaluate whether the results from our screen on established cell lines would potentially translate to the clinic, we tested six of the combination hits in an ex vivo study on two Ewing patient-derived samples. Each hit was evaluated across a fine-grained grid spanning 0.02nM-1*μ*M (Supplementary Fig. 5a). After linearly interpolating (in log-space) to align with the concentration grid applied to the previous Ewing lines, we computed the TI scores for the individual ex vivo models. We found a robust TI response on both models, particularly for the top cell line hit topotecan and talazoparib (Supplementary Fig. 5b). The TI AUC scores were broadly comparable with the median Ewing cell line TI AUC scores (Supplementary Fig. 5d). Similar to the results on EWS lines, we found that the single model TI scores at the low dose were significantly higher than those found in the training set (Supplementary Fig. 5c, *p <* 10^−13^). Moreover, the median ex vivo TI score fell in the 96th percentile of TI scores in the training set.

Finally, we hypothesized that higher order combinations could be discovered through analysis of the pairwise screens. To test this, we ran a triplet screen on the drugs talazoparib, topotecan, and mitomycin over a fine-grained grid spanning 0.1nM - 400nM (see Methods, Supplementary Fig. 6a). Each pairwise combination of the three drugs appeared in our EWS top hits screen, suggesting that the triplet combination of the three would enable additional efficacy. At the single agent level, talazoparib was observed to have low activity with an IC50 nearly 40x higher than topotecan and 4x higher than mitomycin (Supplementary Fig. 6b). Pairwise combinations with talazoparib yielded higher TI for both mitomycin and talazoparib. However, isotonic interpolation analysis revealed that for many choices of cumulative concentration, the addition of mitomycin did not lead to substantial improvements over the combination of talazoparib and topotecan (Supplementary Fig. 6c). Instead, the optimal concentration strategy allocates more towards talazoparib as the total concentration increases but actually reduces the other two drugs (Supplementary Fig. 6d-g). This further supports preclinical evidence that PARP inhibitors sensitize EWS cells to DNA damage ^52,53^, with less of the two DNA damaging drugs needed as talazoparib dosing increases. Overall, the inability of the triplet combination to meaningfully improve on the pairwise score indicates that simple “pairwise additivity” is insufficient to detect effective higher order combinations.

### BATCHIE discovers useful interactions between osteosarcomas and Aurora A kinase inhibitors

We next investigated the ability of BATCHIE to identify high TI combinations for OST lines. The BATCHIE model did not predict any drug pairs would have high TI values over a broad section of OST lines. This is in concordance with the observation that OSTs are more genomically diverse than EWS as OSTs often undergo chromothripsis leading to each tumor having a unique set of rearrangements ^54,55^. Instead of broadly active pairs, we used the BATCHIE model to identify six drugs that were predicted to have high TI values in some pairwise combination for at least one OST line: eltanexor, talazoparib, cladribine, cytarabine, alisertib, and trametinib.

We evaluated all 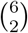 combinations at all four possible high/low dose combinations across all five OST lines and our two control lines. The SJSA-1 cell line failed our quality control checks and was removed from our results. We observed a high concordance (Pearson’s *ρ* = 0.81, *p <* 10^−80^) between the predicted viabilities and the observed viabilities (Fig. 6a) on the remaining four lines and six drugs. We grouped each observation by degree of sensitivity or resistance and noted a clear separation between predicted sensitivity level and therapeutic index (Fig. 6b).

**Figure 6:**
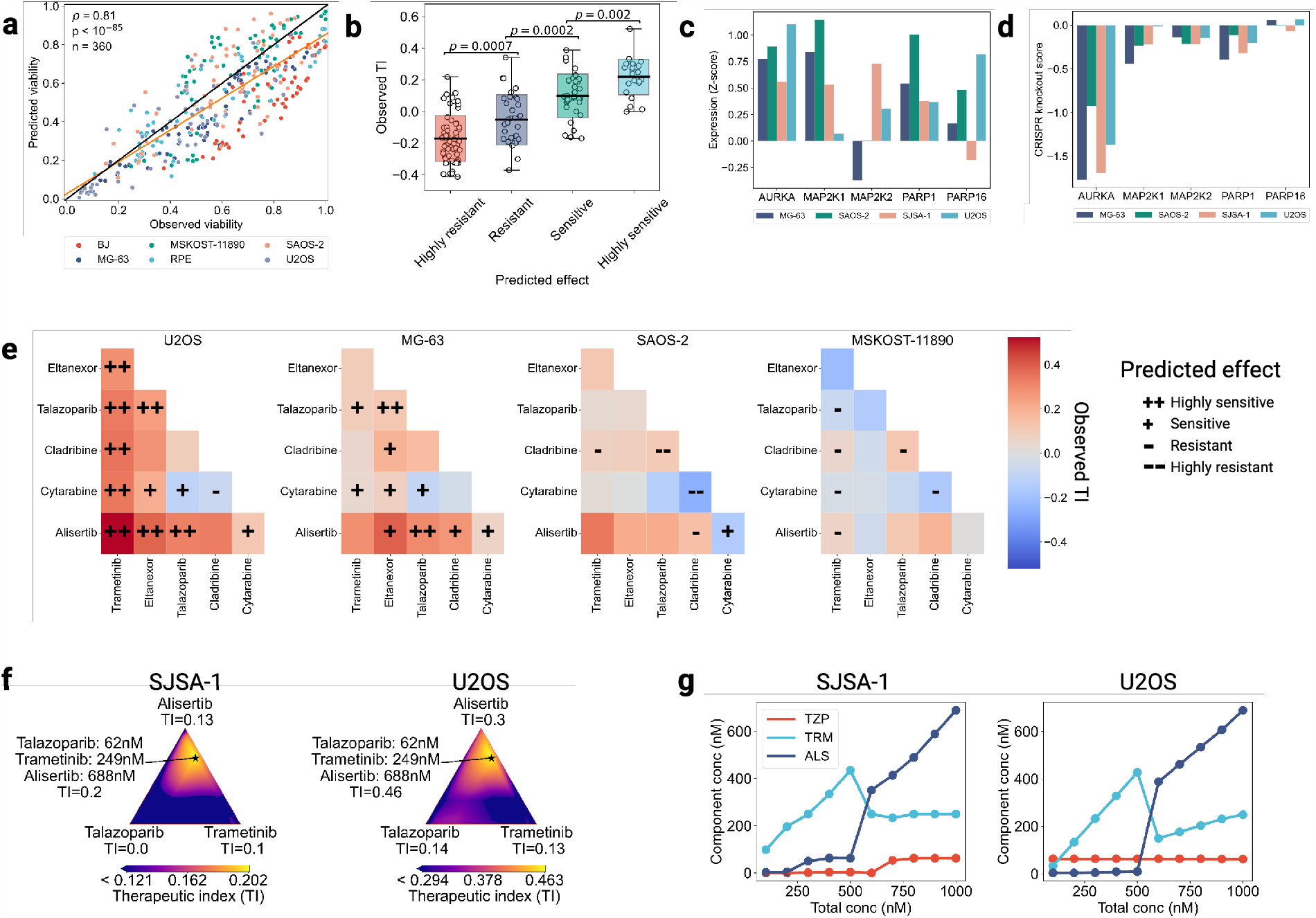
Validation of top osteosarcoma hits. **(a)** Scatter plot of predicted viability against observed viability on OST hit validation data, colored by cell line. **(b)** Observed single cell line TI scores broken down by prediction status, where highly sensitive/sensitive/resistant/highly resistant correspond to TI scores in the range (0.2,1.0]/(0.1,0.2]/[0.2,-0.1)/[-1.0,-0.2), respectively (*n* = 61, 33, 32, 24); *p*-values were computed using Mann-Whitney U-Test. Box plot center lines denote means, box limits denote standard deviations, and whiskers denote extremal values. **(c**,**d)** Z-scores of RNA expression and CRISPR knockout scores from DepMap grouped by gene and colored by cell line. **(e)** Observed single cell line TI scores; entries correspond to largest mean observed TI score over 4 pairwise concentration combinations. Predicted effect designations are identical to those in **(b). (f)** Interpolated TI values for the drug triplet alisertib, talazoparib, trametinib on SJSA-1 and U2OS when the total concentration is held fixed at 1 *μ*M; optimal concentration is denoted by a star. **(g)** Per-drug concentrations that optimize single cell line TI score as a function of total concentration. TZP=talazoparib, TRM=trametinib, ALS=alisertib.

For all pairwise combinations of alisertib, trametinib, and talazoparib, we observed high TI values on U2OS, and both alisertib/trametinib and alisertib/talazoparib achieved substantial TI on both MG-63 and SAOS-2. Alisertib is an Aurora A kinase (AURKA) inhibitor. Using data from Depmap ^56,57^, we investigated the mechanistic rationale behind alisertib as a component of an effective combination. We found that the MG-63, SAOS-2, SJSA-1, and U2OS lines all have high AURKA RNA expression levels (Fig. 6c) and high sensitivity to CRISPR knockout (Fig. 6d). Related work has found AURKA inhibition generally, and alisertib in particular, has been shown to increase the ‘BRCAness’ of cells in vitro ^58^, where BRCAness is defined by defects in the homologous repair pathway that mimick the loss of BRCA1/2^59^. BRCAness has been shown to be correlated with increased sensitivity of OST lines to PARP inhibitors in vitro ^60,61^, making the combination of alisertib and talazoparib particularly rational.

Trametinib is from the class of MEK inhibitors, which have been shown to increase sensitivity to PARP inhibitors in RAS mutant cancer lines ^62^ and ovarian and pancreatic cancer models ^63^. This increased sensitivity to PARP inhibition is possibly due to the downregulation of BRCA2 expression and disruption of the homologous repair pathway ^63^, similar to the effects of AURKA inhibition. While the MEK-PARP approach has been investigated in these other cancers, we are not aware of this combination or MEK-AURKA being investigated in OST, suggesting the BATCHIE discoveries are novel. Again using Depmap data, we observed that all lines in our panel are sensitive to MAP2K1/2 knockout (Fig. 6d) and overexpress MAP2K1 and/or MAP2K2 (Fig. 6d).

Motivated by the pairwise results, we hypothesized that a triplet combining alisertib and talazoparib with one of cladribine, topotecan, and trametinib would be rational. We included cladribine and topotecan as their mechanisms of DNA damage had been shown to lead to increased TI in EWS models in combination with talazoparib, and trametinib was included due to its efficacy in the pairwise combination screen. We evaluated each triplet over a fine-grained grid spanning 0.02nM - 1*μ*M. Isotonic interpolation analysis revealed that in all triplets, on all cell lines, and on most total concentrations, the concentration mixtures that achieved optimal TI scores were dominated by alisertib (Supplementary Fig. 7, Supplementary Fig. 8a). Indeed, only for the highest total concentrations on MSKOST-11890 did we observe diminishing returns for alisertib (Supplementary Fig. 8a). However, we did observe that non-negligible proportions of talazoparib and trametinib made up the optimal mixtures on SJSA-1 and U2OS (Fig. 6f,g). We also found that the inclusion of higher concentrations of alisertib led to a more robust TI score as evidenced by the increased TI AUC scores for all combinations considered in conjunction with alisertib (Supplementary Fig. 8b, Spearman’s *ρ* = 0.3, *p* = 0.0036). Overall, the results suggest strong rationale for high doses of alisertib combined with low-to-moderate doses of trametinib and talazoparib for a subset of patients.

Translation of alisertib-based therapy is challenging as it is currently discontinued due to failing its phase III trial. To evaluate the translatability of the the mechanistic combination, we replicated the above triplet screen with LY329566, an investigational AURKA inhibitor, in place of alisertib. The results of this screen were similar to those with alisertib, with LY329566 generally dominating the optimal concentrations (Supplementary Fig. 9, Supplementary Fig. 10a) and leading to more robust TI scores (Supplementary Fig. 10b). Also similar to the alisertib results, we found that on U2OS, non-neglibigle mixtures of LY329566, talazoparib, and trametinib led to improved TI scores (Supplementary Fig. 9, Supplementary Fig. 10a).

## Discussion

We introduced BATCHIE, an active learning platform that enables large-scale combination drug screens. We derived theory guaranteeing that the batches designed by BATCHIE will always be highly informative. Retrospective simulations on data from previous large-scale combination screens confirmed strong empirical performance of BATCHIE. A prospective study on pediatric cancer cell lines showed BATCHIE screens can enable the rapid discovery of highly efficacious and synergistic drug combinations within libraries of hundreds of drugs.

The probabilistic modeling required in BATCHIE is modular. Our algorithm is able to take any Bayesian model and design optimal batches with respect to that model. We evaluated two different hierarchical models, one focused on viability and another on synergy prediction. BATCHIE showed performance gains with both models, but each could be improved with more sophisiticated modeling. Any modeling improvements that lead to performance gains would be complementary to the gains from BATCHIE screens. Thus, as new predictive models continue to be developed, they can be readily integrated into BATCHIE for improved screening efficiency.

Both our retrospective and prospective experiments were conducted on cancer cell lines. Immortalized 2d cell lines have a number of well-understood limitations and better 3d models such as spheroids and organoids are rapidly being developed to replace them in drug screening ^64^. BATCHIE screens transfer seamlessly to the 3d setting and are arguably more useful here since 3d models tend to have longer doubling times and require more expensive equipment and media, exacerbating the need for efficient screens.

We have focused our implementation on pairwise drug viability screens as they are the most common in the combination literature. However, BATCHIE can be readily adapted to screen the drug interactome for any measurable outcome where experiments are batched. New technologies are emerging that enable a wide range of phenotypic measurement, such as proteome-wide drug effects ^65^, but are currently limited to single-agent screens. Multiplexed CRISPR perturbation screens ^66^ enable combinatorial screens but require specifying a small library of genes. BATCHIE could be used to design optimal libraries in order to efficiently discover synthetic lethal combinations.

Drug combinations represent an increasingly important therapeutic strategy in cancer and other diseases. We expect that approaches such as BATCHIE will be critical to overcoming the combinatorial explosion in the experimental design space as preclinical screens grow to larger libraries and higher-order combinations like triplets and quadruplets. In doing so, these methods will play an integral role in enabling the discovery new combination therapies.

## Methods

### Bayesian tensor factorization model for predicting combination drug response

We use a hierarchical generative model that simultaneously models both single drug and combination drug observations. We treat each drug at each dose individually as a single drug-dose. Observations are modeled as

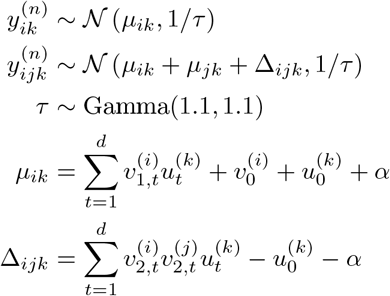

where 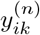 is the *n*-th logit-transformed viability measurement of applying drug-dose *i* to cell line *k*. Similarly, 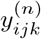 is the logit-transformed viability measurement of applying the combination of drug-doses *i* and *j* to cell line *k*. We also have that *τ* is the global precision of the observations, *μ*_*ik*_ is the mean response of applying *I* to *k*, ∆_*ijk*_ is the combination effect of *i* and *j* applied to *k*, and *α* is the global mean of all the observations. *u*^(*k*)^ ∈ ℝ^*d*^ is the embedding of cell line *k* with the following generative process:

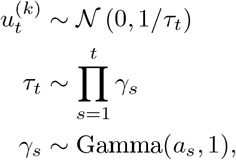

where *τ* ∈ ℝ^*d*^ is the vector of precisions for each cell line embedding coordinate, and it follows a Gamma process prior with elements *γ*_*s*_. The hyper-parameters of the Gamma process are chosen as *a*_1_ = 2 and *a*_*s*_ = 3 for *s* ≥ 2.

The vector 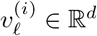 is the order *ℓ* (for *ℓ* ∈ *{*1, 2*}*) embedding of drug-dose *i* with the following Horseshoe prior, also known as a local shrinkage model

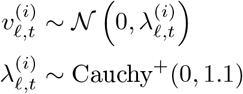

where 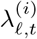 encourages sparsity and Cauchy^+^ is the Cauchy distribution truncated to the positive real numbers.

Also included in the model are offsets 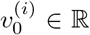 (for drug-dose *i*) and 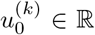 (for cell line *k*). These follow the generative process:

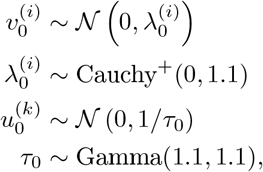

where 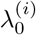 encourages sparsity and *τ*_0_ are precisions.

To fit this model, we utilize Gibbs sampling to sample from the posterior distribution, since all of the relevant priors are conditionally conjugate. For the prospective study, the Gibbs samplers were run for 20K steps.

### Bayesian model for predicting combination drug synergy

For the pure synergy modeling setting, we use the following model.

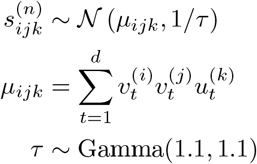

Here, 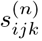 is the *n*-th synergy score between drug-doses *i* and *j* on cell line *k*. It is calculated as 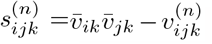, where 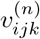is the *n*-th observed viability of applying *i* and *j* to *k*, and 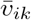 is the average observed viability of applying *i* to *k*. When 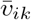 is not available because drug-dose *i* was not tested directly on *k*, then it is imputed by linear interpolation (in log-concentration space) of neighboring concentrations of the same drug.

*μ*_*ijk*_ is the mean synergy value of applying *i* and *j* to *k* and *τ* is global observational precision, and *u*^(*k*)^ ∈ ℝ^*d*^ is the embedding of cell line *k* that follows the same prior as its counterpart in the previous model:

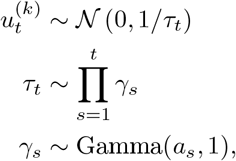

where *τ* ∈ ℝ^*d*^ is the vector of precisions for each cell line embedding coordinate, and it follows a Gamma process prior with elements *γ*_*s*_. The hyper-parameters of the Gamma process are chosen as *a*_1_ = 2 and *a*_*s*_ = 3 for *s* ≥ 2.

The drug-dose embeddings *v*^(*i*)^ ∈ ℝ^*d*^ also follow the same prior as the counterparts in the previous model:

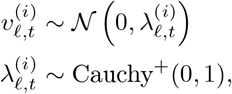

where 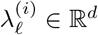 is a vector of scales for the embedding of drug-dose *i*.

This model is also fit via Gibbs sampling.

### Active learning algorithm

Our active learning procedure is a generalization of an optimal active learning procedure called Diameter-based Active Learning (DBAL) ^24,25^. Our approach applies to general probabilistic models that consist of an experimental space 𝒳, an outcome space 𝒴, and a set of parameters Θ. We assume that the likelihoods factorize, so that for a sequence (*x*_1_, *y*_1_), …, (*x*_*n*_, *y*_*n*_) ∈ 𝒳 ×𝒴 and parameter *θ* ∈ Θ, we have

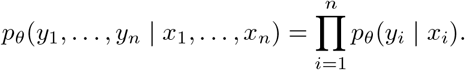

A *plate* of experiments is a sequence *P* = (*x*_1_, …, *x*_*b*_) of experiments *x*_*i*_ ∈ 𝒳 . For a corresponding set of outcomes (*y*_1_, …, *y*_*b*_), we use the shorthand *y*_*P*_ to denote the sequence and 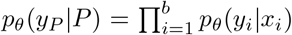 to denote the likelihood of the outcome sequence for a given plate.

In the combination drug setting, the parameters *θ* include all of the parameters from the tensor factorization model, i.e. the *μ*_*ijk*_’s, the 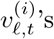, etc. The experiment space consists of triples (*i, j, k*) and pairs (*i, k*) where *i* and *j* are drug-doses and *k* is a cell line. Outcomes in this setting are logit-transformed viabilities, and so the outcome space corresponds to the reals, i.e. 𝒴= ℝ.

Given a prior distribution *π* over Θ and a set of observations (*x*_1_, *y*_1_), …, (*x*_*n*_, *y*_*n*_) ∈ 𝒳 ×𝒴, the posterior distribution over Θ is given by

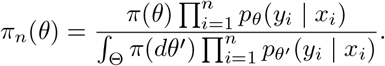

Let *d*(·, ·) be a bounded, non-negative, symmetric distance over Θ. The goal of DBAL-style active learning procedures is to run batches of experiments that will rapidly lead to a posterior *π*_*n*_ with small *average diameter* :

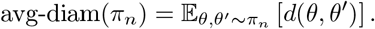

Given a plate *P* and a current posterior distribution *π*_*n*_, our active learning strategy assigns an ideal score to each plate:

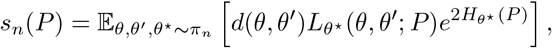

where

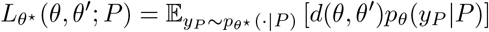

and *H*_*θ*_(*P* ) is the Shannon entropy of the plate *P* under *θ*, i.e.

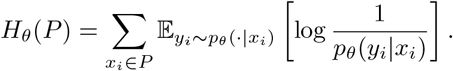

In the case of the normal likelihood (among others), one can explicitly compute the functions 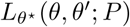 and *H*_*θ*_(*P* ).

Since computing *s*_*n*_(*P* ) requires integrating over the posterior, we cannot hope to do so directly. Instead, we compute a Monte Carlo approximation of *s*_*n*_(*P* ) by sampling *θ*_1_, … *θ*_*m*_ ∼ *π*_*n*_ and computing

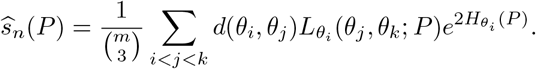

This sum can be further approximated by sub-sampling triples (*i, j, k*) and computing the Monte Carlo average only over the selected triples.

Armed with the estimator 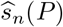, our active learning procedure is to enumerate a set of candidate plates *P*_1_, …, *P*_*T*_ and select the plate 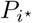 with the lowest score 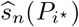. In the supplement, we show that this strategy leads to provably near-optimal guarantees. Specifically, we prove that the convergence rate of BATCHIE is upper-bounded by a function of a problem-specific parameter, called the splitting index, that determines the complexity of the learning problem in the sense that any active learning strategy, regardless of computational power, must have a convergence rate that is lower-bounded by this same splitting index.

To select a batch of plates, we select sequentially. We first select 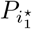 as the plate minimizing 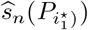. Having selected plates 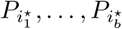, we select plate 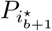 as the plate minimizing 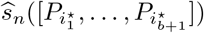, where 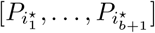 is the plate formed by concatenating the constituent plates 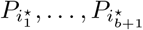 . Observe that this is equivalent to selecting 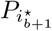 conditioned on having already selected 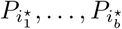 . This iterative strategy of optimization has been shown to enjoy strong theoretical guarantees ^68,69^.

### Retrospective simulations

#### Data retrieval and preparation

We downloaded the ALMANAC ^5^, GDSC^2 6^, and MERCK ^7^ datasets from their respective sources.^1,2,3^ For the MERCK dataset, viabilities were provided. For the ALMANAC dataset, PercentGrowth values were provided. We converted these to viability scores using the formula

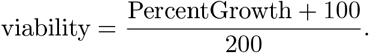

For the GDSC^2^ dataset, well intensity values were provided, along with high control and low control intensities. These were converted to viability scores using the same formula as in the prospective study. For all studies, replicates were averaged to produce a single viability measurement for all recorded cell line, drug-dose 1, drug-dose 2 triplets. Drug-doses that were not present in combination experiments were dropped.

#### Plate and random holdout construction

From the viability measurements, we constructed synthetic plates to closely match the plates used in the prospective sarcoma study. Drug-doses were randomly divided into groups of 20, and initial plates we constructed by considering each cell line *c* and each pair of groups *g, g*^*′*^ and collecting all measurements satisfying that the cell line is in *c*, one of the drug-doses is in *g*, and the other drug-dose is in *g*^*′*^. Due to the biased sampling of drug combinations in the three experimental designs, this resulted in plates of unequal sizes. We corrected for this by greedily merging the two smallest plates within a cell line until a minimum size threshold was met. We then performed a single pass through the plates for each cell line and merged the largest plate with the smallest plate. To align the plate sizes, we selected a threshold and dropped all plates smaller than the threshold and dropped observations from plates larger than the threshold until they had the same number of observations. The threshold was chosen to minimize the total number of observations removed. In each simulation, a random holdout set was created by subsampling 10% of the observations from every plate.

#### Experimental design

For a given dataset and plate construction, we first created an initial covering plate by randomly selecting observations to greedily cover the cell lines and drug-doses. Given the same dataset, set of plates, and initial covering plate, we ran both the Random and BATCHIE methods. At each round of data collection, the methods picked *K* cell lines and 3 plates per cell line. *K* was selected to be 1/10th the number of cell lines in the dataset, rounded down (*K* = 6 for ALMANAC, *K* = 12 for GDSC^2^, and *K* = 3 for MERCK).

#### Posterior inference

At each round, before selecting plates, 200 MCMC samples were drawn from the current posterior using 5 parallel chains, each with a burn-in period of 2000 steps and a thinning factor of 40. These samples were then used to evaluate accuracy on the holdout set and, for BATCHIE, used to select the next set of plates to observe. For each dataset, we ran the plate construction and simulation using 25 different random seeds.

#### Cell line restrictions

For the cell line restriction simulations, cell lines were subsampled uniformly at random. The number of cell lines chosen per round was adjusted accordingly (1/10th of the number of selected cell lines). The number of plates per cell line in each round remained 3.

#### Metrics

For each dataset construction, we trained a model using the full training set, i.e. everything except for the holdout validation set, by drawing 200 MCMC samples from the posterior using 5 parallel chains, each with a burnin period of 2000 steps and a thinning factor of 40. Given a holdout set of experiment/viability pairs (*x*_1_, *y*_1_), …, (*x*_*m*_, *y*_*m*_) ∈ 𝒳 × [0, 1], a fully trained predictor *f*_full_ : 𝒳 → [0, 1], and a candidate predictor *f* : 𝒳 → [0, 1], the normalized accuracy is given by the ratio of *R*^2^ scores:

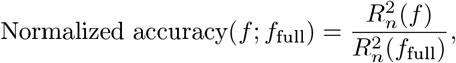

where

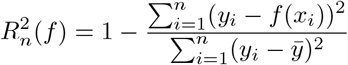

and 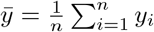 is the empirical mean of observed viabilities.

Efficiency gains/batches saved and experiments saved were computed by calculating the holdout *R*^2^ of the BATCHIE-trained model (at round 15, unless otherwise specified) and then searching for the earliest round at which a RANDOM-trained model had comparable performance. BATCHIE and Random models were only compared within the same random seed, and therefore for the same dataset and plate set construction.

### Pediatric sarcoma combination screen

#### Study design

The prospective sarcoma study consisted of 15 rounds of data collection. At each round, different subsets of cell lines were available based on doubling time and the extent to which they had been used in previous rounds (Supplementary Figure 3c). Adaptive batches were constrained to 3 combination plates each for 3 available cell lines. In the first round and the eleventh round, unseen cell lines were introduced and the plates were selected to greedily cover unseen drug-doses and drug-dose combinations over the new cell lines. In the remaining rounds, plates were selected using the BATCHIE active learning procedure outlined above with the constraint that 3 separate cell lines be chosen, leading to 9 selected plates in total.

In rounds 1-10, each plate was run singly. During the phase I validation, we identified that BATCHIE was sensitive to undetectable random well failures producing corrupted data. As such, in rounds 11-15, plates were run in and quality control checks flagged wells whose duplicates differed by more than 0.5 in viability.

#### Combination plates

Supplementary Fig. 3a shows the general schematic for our combination plate setup. Plates consisted of 384 wells with 16 rows and 24 columns. For row plates, 15 of the rows consist of a single drug applied to each of the corresponding column wells at a particular concentration, excepting two columns that corresponded to high control and low control. The remaining row of the row plate was filled with Dimethylsulfoxide (DMSO), also with two control columns. For column plates, 21 of the columns consist of a single drug applied to each of the corresponding row wells at a particular concentration. The remaining three columns are the high and low control columns (aligned to spatially match the corresponding high/low columns in the row plate) and a column filled only with DMSO.

When a row and column plate are combined, the resulting combination plate contains 15× 21 = 315 wells that correspond to all combinations of the constituent drug-doses, 15 + 21 = 36 wells that correspond to all single drug-doses, 15 high-control wells, and 15 low-control wells.

Our drug library consisted of 206 drugs, with four of the drugs duplicated to allow for the resulting 210 drugs to be evenly divided over 14 row plates and 10 column plates. Each row and column plate was constructed at 2 different doses: 0.1 *μ*M and 1 *μ*M, leading to a total of 28 row plate choices and 20 column plate choices.

Finally, a full plate consists of a cell line, a row plate, and a column plate. Viabilities for a non-control well *w* are calculated as

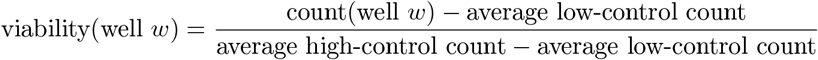

where count(well *w*) is the reading at well *w* and average high/low-control counts are the averages of the readings at the corresponding high/low-control wells.

#### Therapeutic index

We define the *in vitro* therapeutic index (TI) to be a differential score that compares two groups of viabilities: one for the target set of cell lines and one for the control set. A high TI corresponds to low viability for most target cells and high viability for all control cells. To calculate TI, we take the difference between the minimum viability on the control lines and the median viability on the target lines:

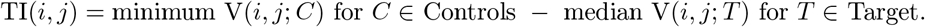

where V(*i, j*; *N* ) is the viability of drug-dose pair (*i, j*) on cell line *N* .

#### Model visualizations

To determine if BATCHIE discerned the relative mechanistic similarities and differences of the drugs tested in an unsupervised manner, we labeled each drug assessed with simplified mechanisms of action (MoAs) and used low-rank approximation and dimensionality reduction to visualize resulting clusters. We used MoAs from the Genomics of Drug Sensitivity in Cancer (GDSC) database^4^ to annotate the 98 drugs assessed in both our study and the GDSC. For the remaining 112 drugs in our library that were not in the GDSC, we manually added MoAs that conformed to the GDSC categories. Mechanisms were adjudicated according to drugs’ International Nonproprietary Name (INN) suffixes, documented mechanisms in the literature, and U.S. Food and Drug Administration (FDA) package inserts for approved molecules. All drugs were categorized as belonging to one of nineteen MoA categories.

Predicted viabilities of the drugs administered as monotherapies were logit-transformed and normalized to a zero-one range. Non-negative matrix factorization (NMF) was performed using the Python package `scikit-learn`’s NMF module at a random initialization ^70^. The decomposed feature matrix represented latent embeddings of the drugs, where each dosage and drug combination was treated as a unique entry, and the coefficient matrix represented latent embeddings of the cell lines tested. By assessing the mean squared error of the resulting low-rank approximation, we determined 15 to be the optimal number of latent components using the elbow method as implemented in the `kneed` Python package ^71^. For plotting purposes, we reduced the dimensionality of the feature matrix using t-distributed stochastic neighbor embedding (t-SNE) as implemented in the scikit-learn t-SNE module ^70^. Results from this analysis are shown for high dose (1.0 *μ*M) drugs in Supplementary Fig. 4e.

To determine if BATCHIE also discerned more fine-grained drug MoAs, we selected 8 more refined molecular mechanisms or pharmaceutical classes that were recurrent among the drugs evaluated. We then identified 2-3 drugs belonging to each such group. We calculated the Pearson correlation coefficients between the BATCHIE-predicted combination therapy viabilities for each drug at both doses assessed. We plotted the resulting hierarchically clustered heatmap using the Python package Seaborn’s clustermap function^72^. These results are show in Supplementary Fig. 4d.

#### Ex vivo analysis

For six of the drug combinations from the top EWS hits, we collected dose-response data on 2 additional patient-derived cell lines. The drugs were tested on a regular grid spanning 0.02nM - 1000nM, and the plates were run in duplicates.

As the concentrations tested in this stage did not align with the concentrations tested in the top hit validation, we performed linear interpolation of the mean observed viabilities (in *log*_10_ concentration space), in order to compute TI scores with respect to the control data observed in the top hit validation analysis.

#### Triplet analysis

For our triplet studies, we collected dose-response data along a regular grid (0.1nM - 400nM for the Ewing study and 0.02nM - 1*μ*M for the osteosarcoma study). Plates were run in duplicate.

For each cell line and triplet of drugs under consideration, we fit a multi-dimensional isotonic regression to the observed viabilities, restricting the regressed variables to monotonically decrease as a function of dose. Mathematically, we solved

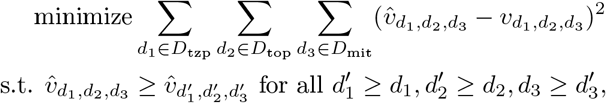

where *D*_*x*_ is the set of concentrations used on drug *x*, and 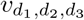 is the mean observed viability applying drug 1 at concentration *d*_1_, drug 2 at concentration *d*_2_, and drug 3 at concentration *d*_3_. For any candidate set of concentrations, we linearly (in log-concentration space) interpolated its viability from the smoothed values 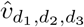 .

#### Gene expression and CRISPR knockout analysis

Both the RNA expression data and CRISPR knockout data were obtained from the DepMap data portal^5^. The RNA expression data is represented as a matrix *X* ∈ ℝ^*N* ×*G*^, where *N* is the number of DepMap cell lines, *G* is the number of protein coding genes in the DepMap library, and *X*_*cg*_ = log_2_(1 + TPM_*c,g*_), where TPM_*c,g*_ is the transcripts per million (TPM) for gene *g* in cell line *c*. Z-scores for the gene expression data were computed by standardizing over cell lines, i.e.

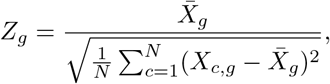

where 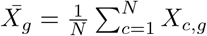.

The CRISPR knockout data is represented as a matrix *Y* ∈ ℝ^*N* ×*G*^, where *Y*_*c,g*_ is a harmonized score representing the effect of knocking out gene *g* in cell line *c*.

### Statistical analyses

All Mann-Whitney U-tests, Spearman’s rho calculations, Pearson’s rho calculations, Pearson’s chi-square calculations were computed using Scipy’s ^73^ `stats` package, using the default settings. All p-values from Mann-Whitney U-tests, Spearman’s rho tests, and Pearson’s rho tests were computed using two-sided alternatives. The permutation test in Fig. 4g was custom coded. Unless otherwise noted, all tests were performed independently of each other.

### Wetlab protocols

#### Cell lines

Established cell lines (SJSA-1, U2OS, SAOS-2, MG-63, A673, SKNEP, MDA-MB-231, RPE and BJ) were obtained from the American Type Culture Collection (ATCC). TC-71 was obtained from the Children’s Oncology Group cell line repository. Wit49 was provided courtesy of Dr. Herman Yeger (Toronto, Canada). Kelly was obtained from the DSMZ-German Collection of Microorganisms and Cell Cultures GmbH. MSKEWS-83311, MSKEWS-83033, MSKEWS-38338, MSKEWS-66647, MSKOST-11890, and MSKRMS-12808 were generated from patient-derived xenograft (PDX) tumor tissue established from patient tumors treated at Memorial Sloan Kettering Cancer Center (MSKCC). Tumor tissue was dissociated utilizing the Human Tumor Dissociation Kit (Miltenyi Biotec) according to manufacturer’s recommendations or using an in-house combination of Collagenase Type II (Gibco), Dispase II (Millipore Sigma), Deoxyribonuclease I (Millipore Sigma) in DMEM (Gibco) containing 10% fetal bovine serum (Corning), followed by mechanical dissociation with Macs dissociator (Miltenyi). MSKCC patients provided informed and signed consent and enrolled onto a tumor profiling research study (Genomic profiling in cancer patients; NCT01775072) approved by the MSKCC Institutional Review Board under protocol IRB#12-245, #06-107, and #17-387 to enable tumor cell line generation. PDX tumor models were generated under MSKCC Institutional Animal Care and Use Committee protocol #16-08-011. Additional cell line information and culture conditions are detailed in Table S1. Media was supplemented with 1% Anti-Anti (Gibco) at the time of drug screening. Authentication of established cell lines by short tandem repeat (STR) was performed. Validation and authentication of PDX-derived cells lines was accomplished by next-generation targeted sequencing using MSK-IMPACT ^74^ and matched with source patient tumor. Optimal seeding densities for drug screens were empirically determined for each cell line.

#### Drugs

The studies were comprised of 206 drugs obtained from multiple sources including Selleckchem, MedChemExpress, Sigma-Aldrich, Tocris, LKT laboratories Inc., Xcess Biosciences, and the National Cancer Institute (NCI) Division of Cancer Treatment and Diagnosis. A list of all screened chemicals and sources are provided in Supplementary Table S2.

#### BATCHIE plates

All assay plates contained baseline viability (high control) and complete cell killing controls (low control) consisting of 1% DMSO (v/v) and 1 *μ*M “killer mix”, a proprietary mixture of cytotoxic drugs at 1% DMSO (v/v), respectively. Phase I and phase II drugs were prepared in 100% DMSO (v/v) and added onto 384-well microplates to generate the “200X source plates” with drugs at a concentration of either 200 μM or 20 μM. To facilitate combination testing, drugs were arrayed in a “row” or “column” format with “row plates” consisting of 15 drugs arrayed in rows per plate and “column plates” consisting of 21 drugs arrayed in columns per plate with the same drug concentration for each drug in both plate configurations.

Drugs were combined using the Apricot Designs Personal Pipettor (SPT Labtech) which mixes a row plate with a column plate and water into a new 384-well plate to generate a “10X intermediate plate” of 10 μM or 1 μM in 10% DMSO (v/v). Subsequently, the 10X intermediate plates are stamped into “assay plates” to be combined with cells generating the final 1X concentration of 1 μM or 0.1 μM in 1% DMSO (v/v) (Supplementary Fig. 11, Supplementary Fig. 12a).

For validation studies, 2-drug combinations were evaluated for random unseen combinations in five osteosarcoma, five Ewing sarcoma, and two non-cancer cell lines; high predicted therapeutic index (TI) in five Ewing sarcoma and two non-cancer cell lines; and Ewing sarcoma ex vivo in two Ewing sarcomas (Supplementary Fig. 11). 7-doses (Plate 1) and 9-doses (Plate 2) 4-fold serial dilutions were prepared into 96-well plates to generate 20X source plates at 8 μM or 20 μM in 10% DMSO (v/v) as the highest concentrations. The plates were further consolidated and combined into 384-well 10X intermediate plates at 4 μM or 10 μM in 10% DMSO (v/v). The 10X intermediate plates were then transferred into assay plates along with cells to produce a 1X concentration of 0.4 μM or 1 μM in 1% DMSO (v/v) (Supplementary Fig. 12b).

Higher order validation plates for Ewing sarcoma and osteosarcoma studies (Supplementary Fig. 11) were performed by first generating 7-doses (Drug 1), 5-doses (Drug 2) and 5-doses (Drug 3) 4-fold serial dilutions, obtaining 30X source plates at 12 μM or 30 μM in 10% DMSO (v/v) as the highest concentrations. The three 30X sources plates were combined to create the 10X intermediate plates with maximum concentrations of 4 μM or 10 μM in 10% DMSO (v/v) and stamped to combine with cells to yield 1X assay plates of 0.4 μM or 1 μM in 1% DMSO (v/v) (Supplementary Fig. 12c). Five Ewing sarcoma and two non-cancer cell lines were used for the Ewing sarcoma validation studies, and five osteosarcoma lines for the osteosarcoma studies.

For osteosarcoma studies validating agents with high predicted therapeutic indices (TI), drugs were plated in 384-well 20X source microplates using the row and column format described above for phase I and phase II (Supplementary Fig. 11) at 20 μM or 2 μM in 10% DMSO (v/v). The sources plates were combined into 10X intermediate plates yielding 10 μM or 1 μM in 10% DMSO (v/v). Assay plates were stamped from the 10X intermediate plate and combined with cells for a final drug concentration of 1 μM (high concentration) or 0.1 μM (low concentration) in 1% DMSO (v/v). All possible two drug combinations were obtained – high/high, high/low, low/high and low/low. Five osteosarcoma and two non-cancer cell lines were used.

#### Cytotoxicity assay

Cells were plated at their optimized seeding densities and in their corresponding media (Supplementary Table S1) into 384-well clear-bottom black assay plates (BATCHIE plates) using the MultiDrop^®^ 384 dispenser (Thermo Fisher Scientific). After incubating cells in drug for 72 hours at 37°C and 5% CO2 in a Heracell™ 240i incubator (Thermo Fisher Scientific), Alamar Blue (Sigma-Aldrich) is added using the MultiDrop™ Combi 384 dispenser (Thermo Fisher Scientific) and incubated for another 24 hours. Fluorescence signal readout was acquired using a Cytation™ 5 multimode reader (Agilent Biotek) using the monochromator with an excitation of 555/20 nm and emission of 596/20 nm.

## A Supplemental results

**Supplementary Figure 1:**
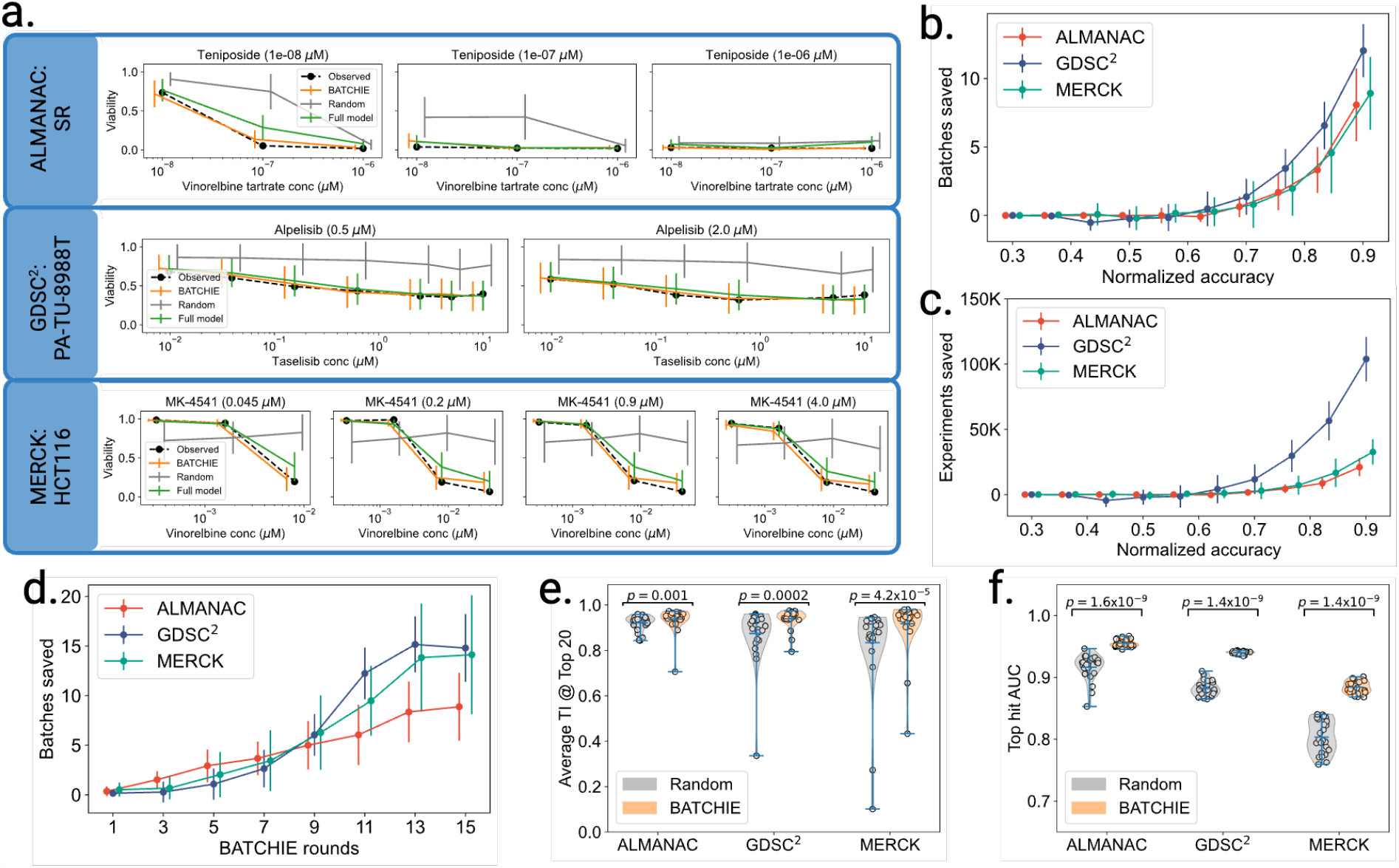
More retrospective results. **(a)** Example dose-response curve fits for completely unseen pairs of drugs combos and cell lines in each of the datasets after 15 rounds of data collection. Model fit on the entire dataset (Full model) depicted for comparison. **(b-c)** Batches and experiments saved as a function of the target normalized accuracy. **(d)** Batches saved as a function of the number of BATCHIE rounds. **(e)** Average therapeutic index (TI) of the top 20 picks after 15 rounds of data collection. **(f)** Area under the receiver operating characteristic (ROC) curve when selecting picks with therapeutic index in the 99th percentile. Error bars in **(b-d)** denote standard deviations. p-values for **(e**,**f)** computed using a Mann-Whitney U Test.

**Supplementary Figure 2:**
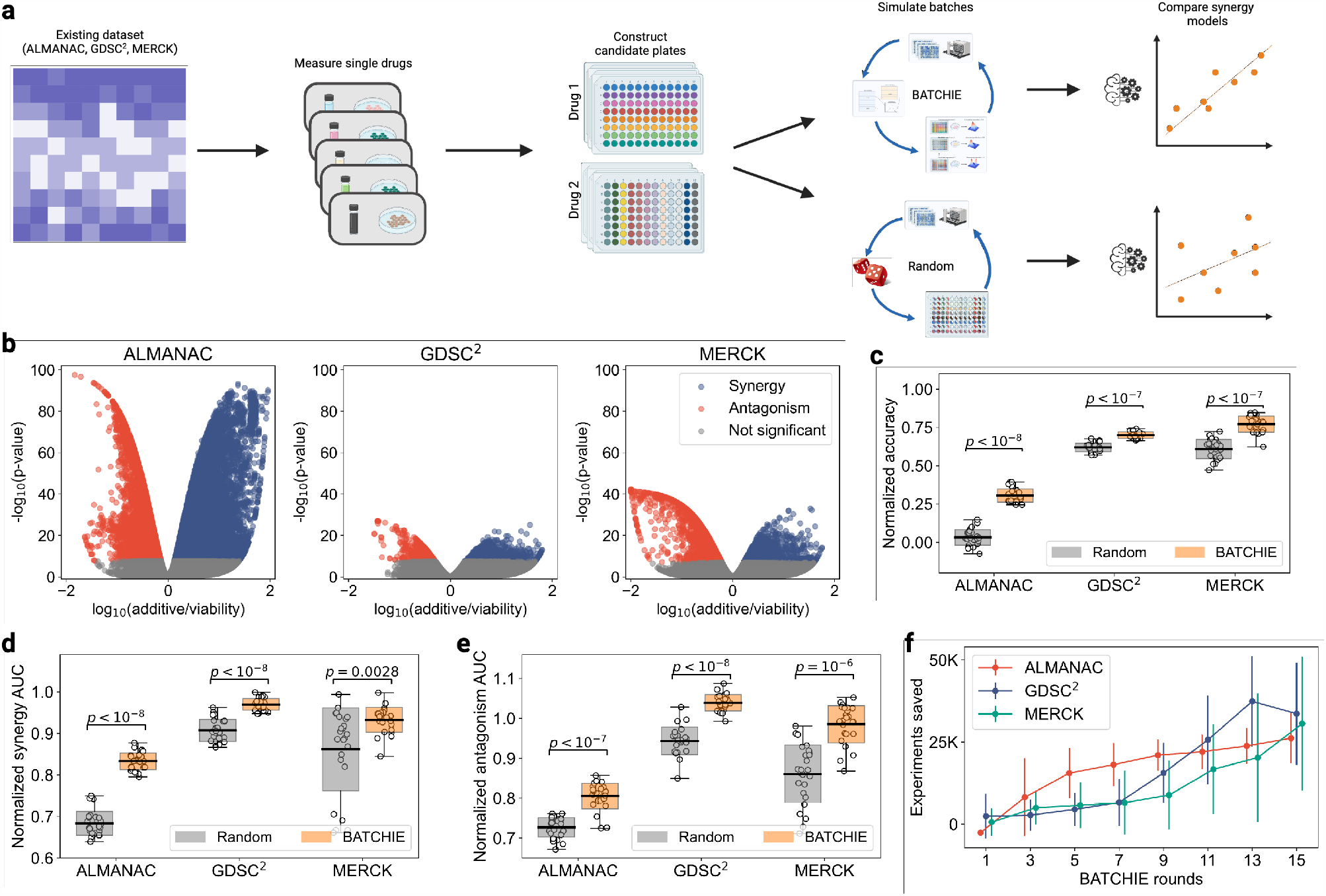
Retrospective Bliss results. **(a)** Retrospective Bliss studies were conducted by first subtracting out the individual drug effects for each combination observation. Datasets were broken into combination plates, and the BATCHIE and Random strategies were simulated over the dataset. **(b)** Volcano plots of p-values compared against the ratio of viability predicted by additive model v.s the observed viability (cutoff at *α* = 0.05 threshold using Benjamini-Hochberg ^75^ correction). **(c)** Normalized accuracy on predicting holdout Bliss scores. **(d)** Normalized AUC of the ROC curve for predicting synergy. **(e)** Normalized AUC of the ROC curve for predicting antagonism. **(f)** Excess experiments needed by Random to achieve accuracy of BATCHIE as a function of the number of BATCHIE rounds. Error bars denote standard deviations. Models in **(c-e)** were trained on data collected up to, and including, round 15. For boxplots in **(c-f)**, center lines denote means, box limits denote standard deviations, and whiskers denote extremal values.

**Supplementary Figure 3:**
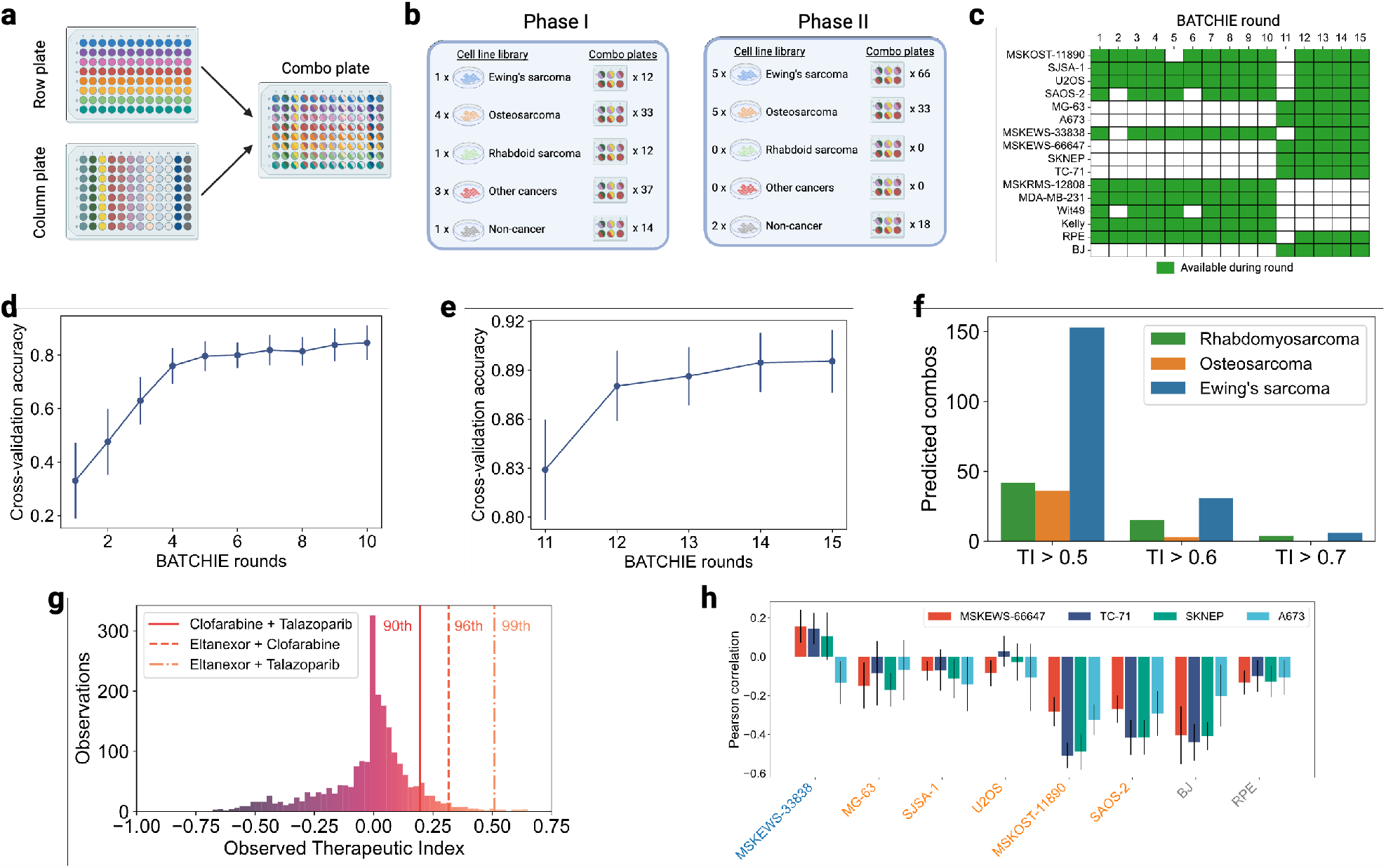
More prospective study design info. **(a)** Illustration of how combination plates are created by overlaying a row plate with a column plate. **(b)** Cell line and combo plate breakdown for phase I/II. **(c)** Availability of cell lines across BATCHIE rounds. **(d-e)** Cross-validation accuracy of the model as a function of the rounds of data collection. **(f)** Cancer-type breakdown of the high TI predictions at round 10. **(g)** TI results for 3 combos tested on a Ewing’s sarcoma cell line at round 10 in comparison with the BATCHIE-collected data observed up to round 10. **(h)** Predicted correlation between the 4 Ewing’s sarcoma lines added at round 11 and the other cell lines used in rounds 11-15. Predictions are with respect to the model trained on data collected up to, and including, round 11. Error bars in **(d**,**e**,**h)** denote standard deviations.

**Supplementary Figure 4:**
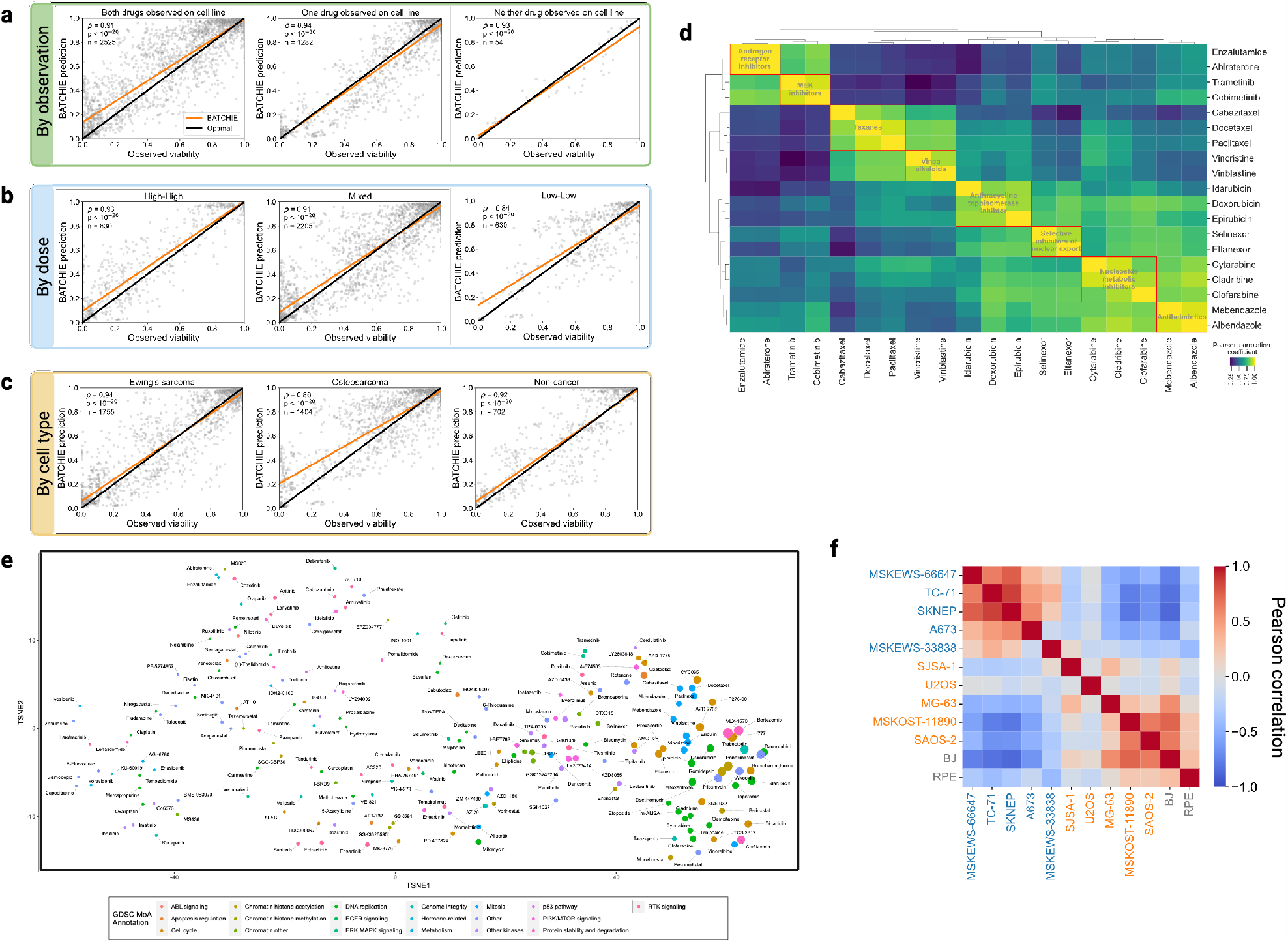
More prospective study results. **(a-c)** Scatter plots of predictions on random validation set broken down by previous observation status, drug concentration, and cell type. *ρ* is Pearson’s *ρ*. **(d)** Heatmap and hierarchical clustering dendrogram of the Pearson correlation coefficients for the BATCHIE-predicted combination therapy viabilities for a selection of drugs at both high and low doses. **(e)** NMF-decomposed feature matrix of the BATCHIE-predicted viabilities projected to two dimensions using t-SNE. Results are shown for high dose experiments. Marker size represents negative mean log_10_-transformed viabilities for the high dose experiments containing a given drug. **(f)** Predicted correlations between all cell lines used in rounds 11-15. For all subfigures, predictions are with respect to the model trained on data collected up to, and including, round 15.

**Supplementary Figure 5:**
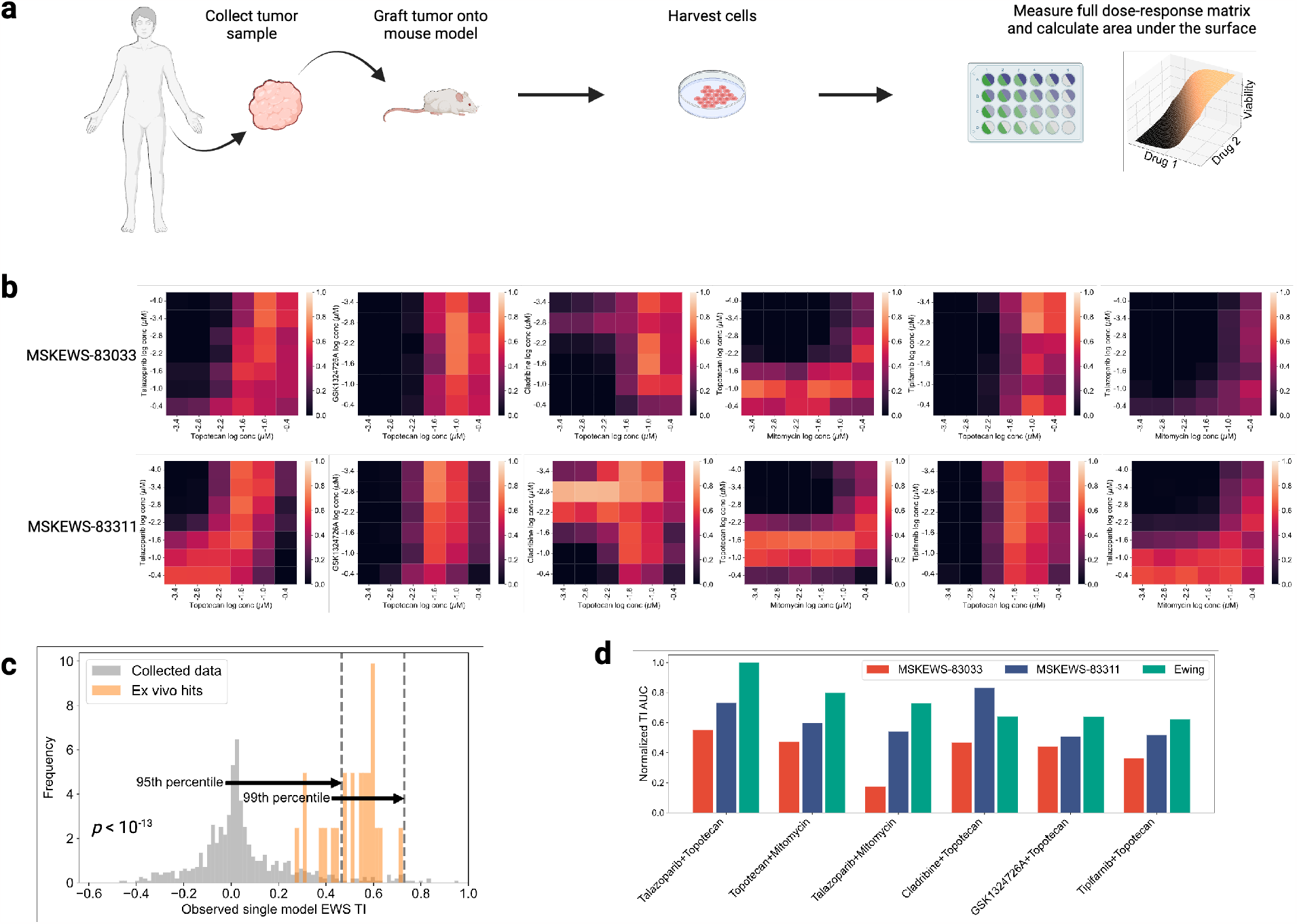
Ewing ex vivo TI results. **(a)** Ex vivo screens were performed by first grafting a patient tumor sample onto a mouse model, harvesting the cells, and then screening the pairwise combinations over a regularly-spaced pairwise combination grid. **(b)** Heatmaps for selected drug combinations on grouped by patient sample. **(c)** Histogram of observed single model EWS TI values for combinations at 0.1*µ*M concentration in BATCHIE-collected data (gray, *n* = 659) and ex vivo data (orange, *n* = 24). Percentiles are drawn with respect to BATCHIE-collected data. *p*-value computed using Mann-Whitney U-test. **(d)** Observed AUCs with respect to the TI dose-response surface. Ewing refers to the TI score with respect to the median EWS cell line response.

**Supplementary Figure 6:**
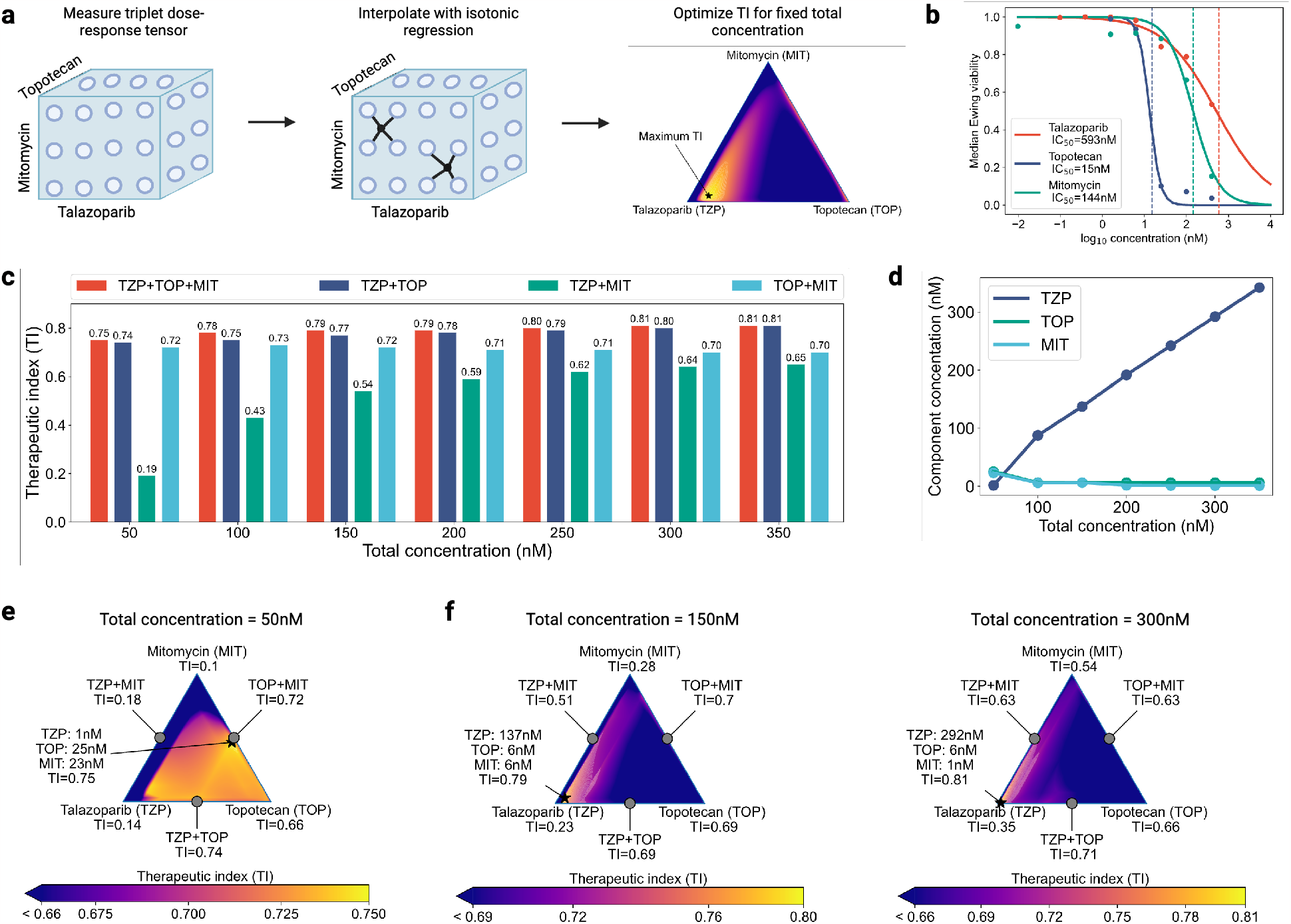
Ewing triplet study. **(a)** Triplet analysis was performed by measuring the 3-way dose response tensor at a grid of concentrations spanning 0.1nM - 400nM. Viabilities were smoothed by first applying isotonic regression, and then linearly interpolating the regressed values. TI scores were computed using the interpolated values over a fine-grid, and the combinations optimizing TI were selected. **(b)** Observed medians of the Ewing viabilities for the single agents in the triplet analysis, along with the corresponding logistic curve fit and the corresponding IC_50_ values. **(c)** TI scores as a function of total concentration for the optimal combination of all three drugs (TZP+TOP+MIT) and the optimal pairwise combinations (TZP+TOP, TZP+MIT, TOP+MIT). **(d)** Per-component concentrations for the optimal combination of the three drugs as a function of total concentration. **(e-g)** Simplex plots of interpolated TI values as the proportion of the three drugs varies. The optimal value is denoted with a star.

**Supplementary Figure 7:**
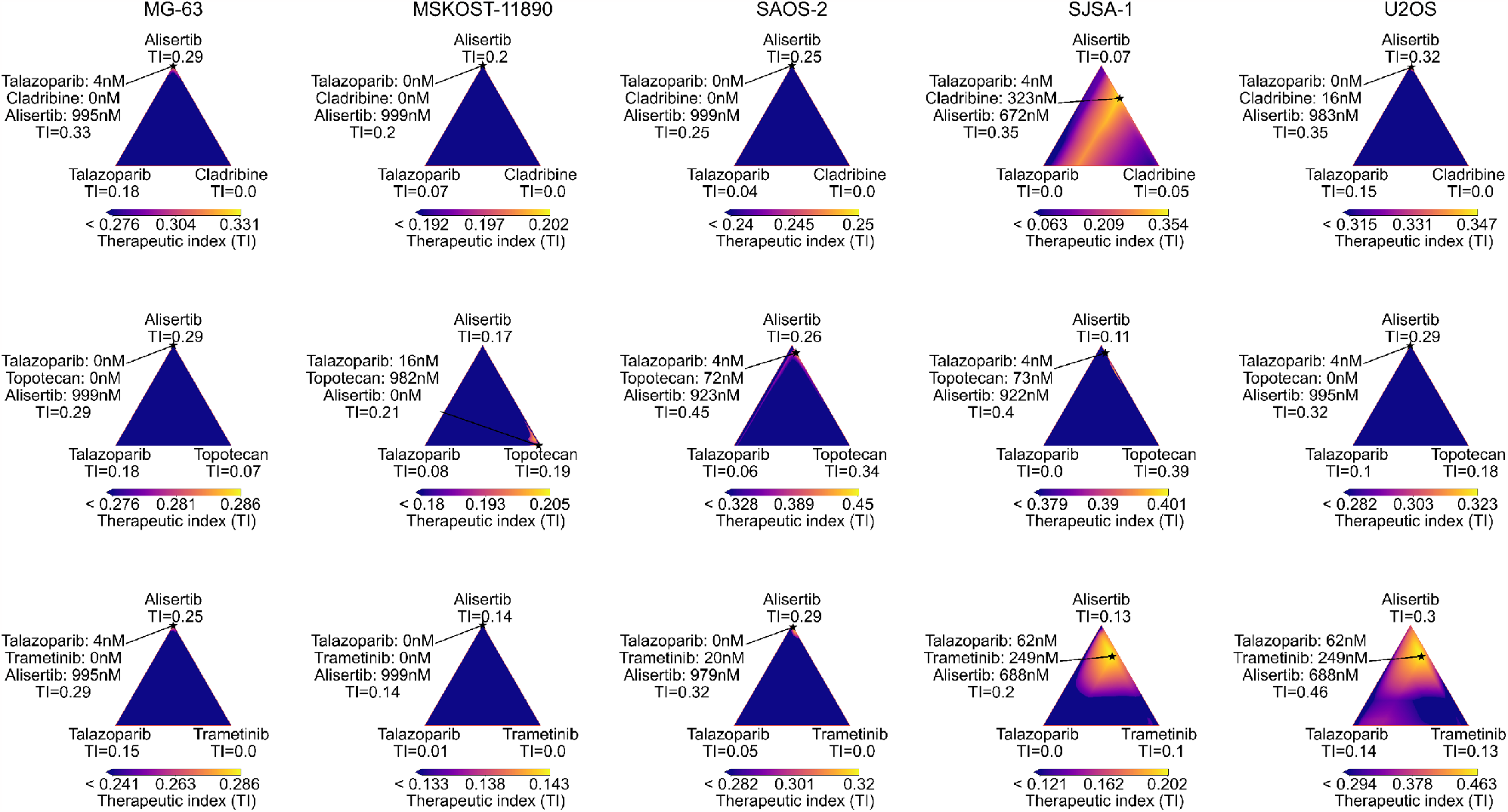
Osteosarcoma/Alisertib triplet simplex analysis. Simplex plots of interpolated TI values as the proportion of each drug varies. The optimal value is denoted with a star.

**Supplementary Figure 8:**
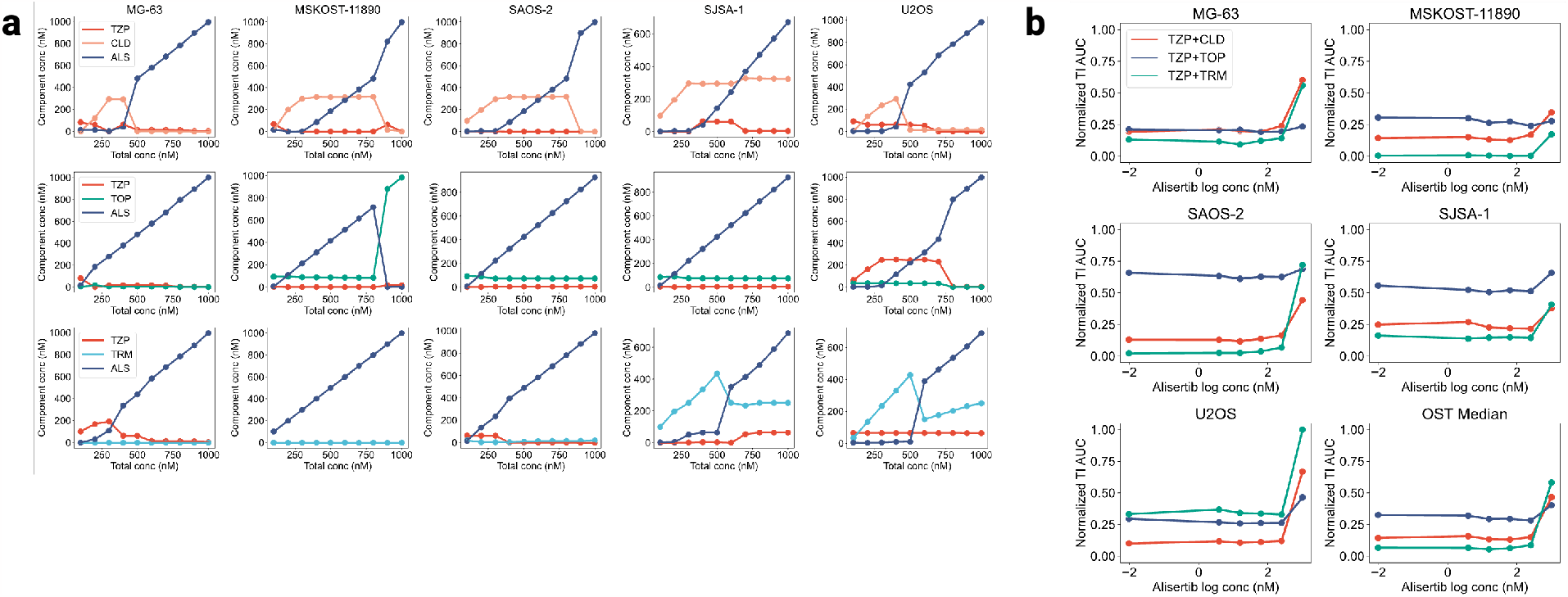
Osteosarcoma/Alisertib triplet concentration analysis. **(a)** Per-component concentrations for the optimal combination of the three drug triplets as a function of total concentration. **(b)** Max-normalized values of the area under the TI surface for pairs of drugs as a function of Alisertib concentration. Spearman’s *ρ* = 0.3, *p* = 0.0036 for correlation between alisertib concentration and TI AUC on osteosarcoma lines.

**Supplementary Figure 9:**
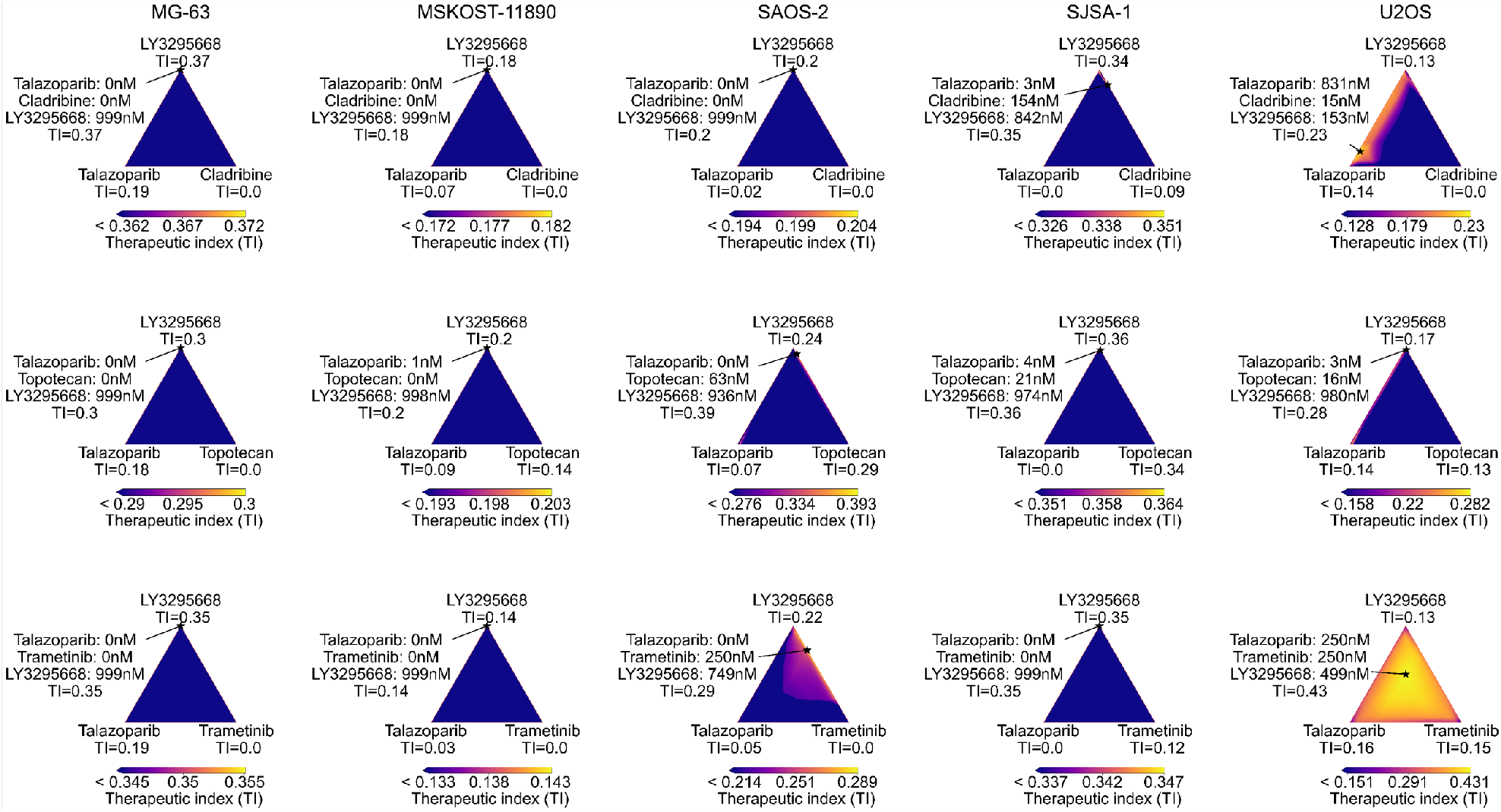
Osteosarcoma/LY3295668 triplet simplex analysis. Simplex plots of interpolated TI values as the proportion of each drug varies. The optimal value is denoted with a star.

**Supplementary Figure 10:**
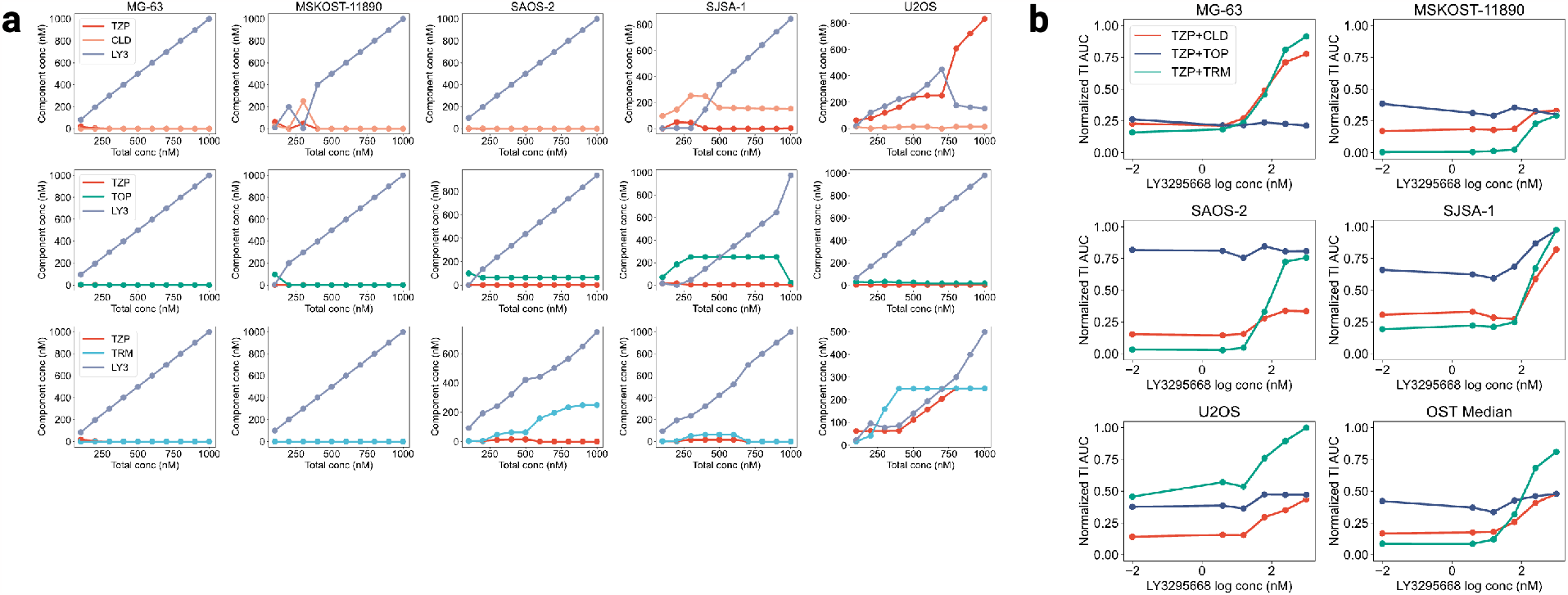
Osteosarcoma/LY3295668 triplet concentration analysis. **(a)** Per-component concentrations for the optimal combination of the three drug triplets as a function of total concentration. **(b)** Max-normalized values of the area under the TI surface for pairs of drugs as a function of LY3295668 concentration.

**Supplementary Figure 11:**
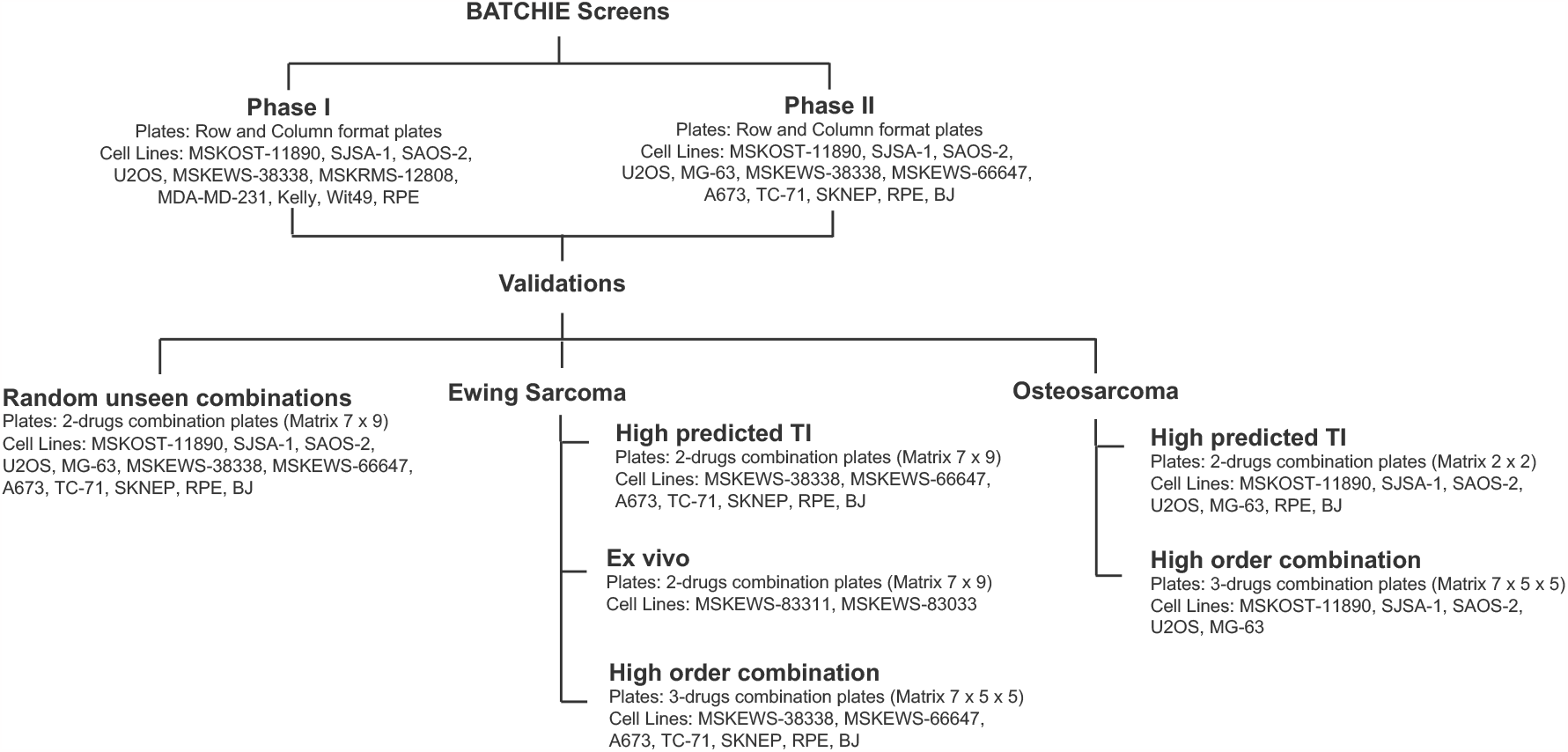
BATCHIE screen details. Diagram of the screens performed in BATCHIE.

**Supplementary Figure 12:**
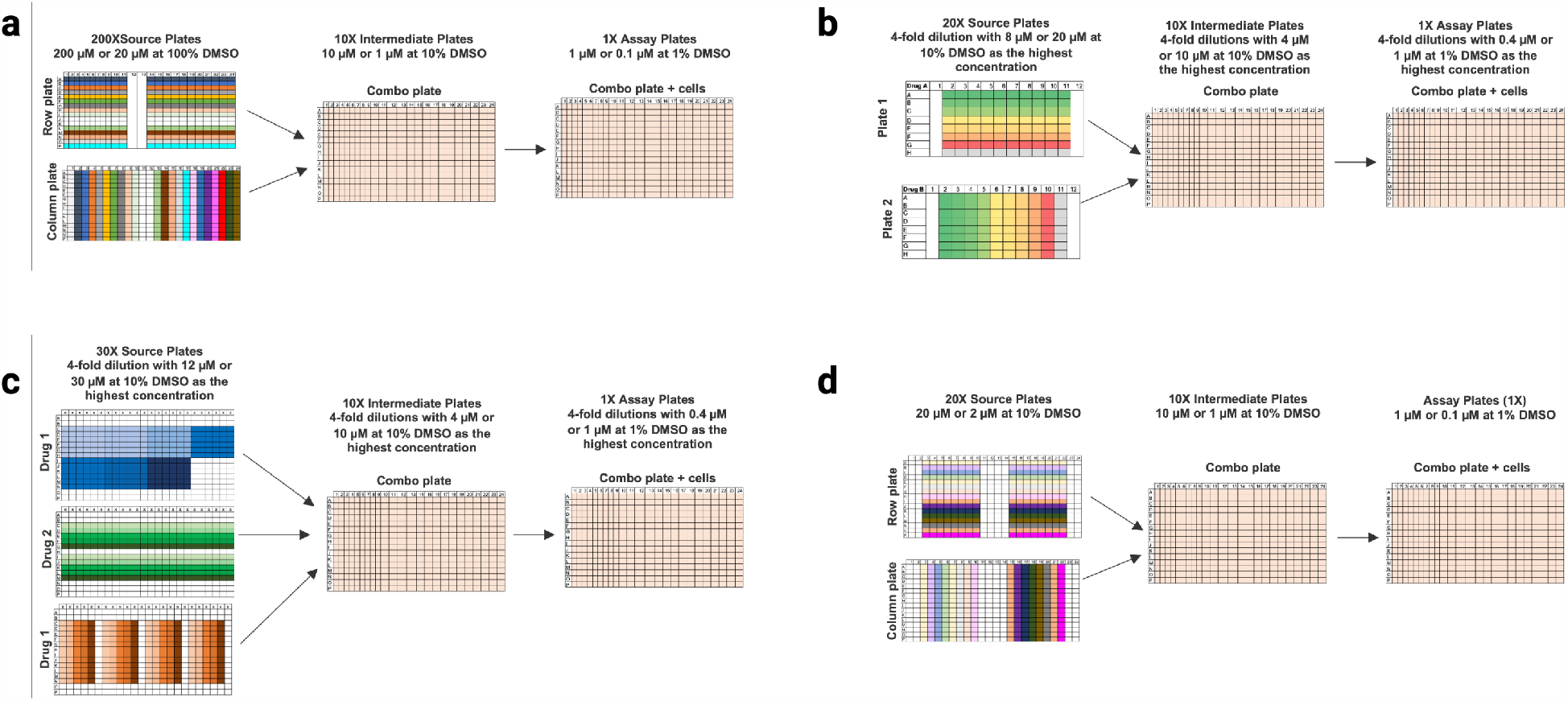
Plate maps for BATCHIE screens. **(a)** Phase I and Phase II plates. **(b)** 2-drugs combination plates (Matrix 7 x 9), **(c)** 3-drugs combination plates (Matrix 7 x 5 x 5), **(d)** 2-drugs combination plates (Matrix 2 x 2).

## B Theory

### B.1 Theoretical setting

Let 𝒳 denote a data space and 𝒴 denote a response space. Let 𝒟 denote a marginal distribution over 𝒳. Our goal is to model our data with some parametric probabilistic model *P*_*θ*_(· ;·), where *θ* lies in a parameter space Θ and *P*_*θ*_(*y*; *x*) denotes the probability (or density) of observing *y* ∈ *𝒴* at data point *x* ∈ *𝒳* . We will use the notation *y*∼ *P*_*θ*_(*x*) to denote drawing *y* from the density *P*_*θ*_(· ; *x*).

We consider models that factorize across data points, that is for *x*_1_, …, *x*_*n*_ ∈ *𝒳*^*n*^ and *y*_1_, …, *y*_*n*_ ∈ *𝒴*^*n*^, we have

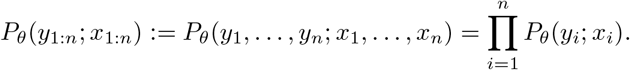

For a data point *x* ∈ *𝒳* and parameter *θ* ∈ Θ, we denote the entropy of the response to *x* under model *θ* as

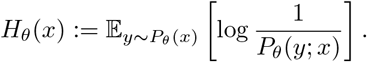

We will take a Bayesian approach to learning. To that end, let *π* denote a prior distribution over Θ. Given observations (*x*_1_, *y*_1_), …, (*x*_*n*_, *y*_*n*_), denote the posterior distribution as

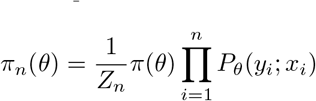

where 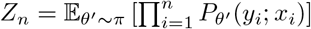 is the normalizing constant to make *π*_*n*_ integrate to one. In this paper, we will assume that we are in the well-specified Bayesian setting, i.e. there is some ground-truth *θ*^⋆^∼*π*, and when we query point *x*_*i*_, the observation *y*_*i*_ is drawn from 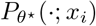. We will also use the notation *π*_*n*_(*y*; *x*) to denote the posterior predictive density

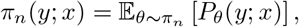

and the notation *y* ∼ *π*_*n*_(*x*) to denote drawing *y* from the density *π*_*n*_(*y*; *x*).

A *risk-aligned distance* is a function *d* : Θ × Θ *→* [0, 1] satisfying two properties for all *θ, θ*^*′*^ ∈ Θ:

- **Identity**. i.e., *d*(*θ, θ*) = 0.
- **Symmetry**. i.e. *d*(*θ, θ*^*′*^) = *d*(*θ*^*′*^, *θ*).

The requirement that *d*(*θ, θ*^*′*^)≤ 1 is not onerous – any smooth distance over a bounded space can be transformed into a distance that satisfies this requirement by rescaling. In our setup, a risk-aligned distance encodes our objective: if we committed to the model *θ* when the true model was *θ*^⋆^, then we expect to suffer a loss of *d*(*θ, θ*^⋆^).

The goal in our setting is to find a posterior distribution *π*_*n*_ with small *average diameter* :

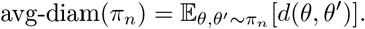

To see that this is a reasonable objective, observe that if *θ*^⋆^∼ *π* and (*x*_1_, *y*_1_), …, (*x*_*n*_, *y*_*n*_) is generated according to 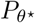, then after observing this data, *θ*^⋆^ is distributed according to *π*_*n*_. If we make predictions by sampling a model from *π*_*n*_, the expected risk of this strategy is exactly the average diameter. Moreover, even without this Bayesian assumption, the risk of this strategy can still be bounded above as a function of the average diameter ^24^.

### B.2 Probabilistic DBAL (pdbal)

We term our active learning selection criterion Probabilistic DBAL (pdbal), as it can been as a generalization of the DBAL active learning criterion ^24^ to the probabilistic setting. Formally, the criterion boils down to a score function: given *x* ∈ *𝒳*,

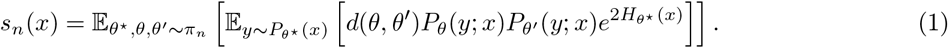

#### Constant entropy models

When Θ parameterizes location models with fixed scale parameters, the entropy term is constant. We rewrite the objective for these models as

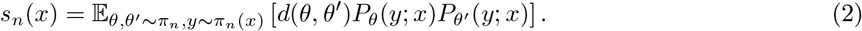

For readability, we will discuss approximations of Eq. (2). Extending these ideas to Eq. (1) can easily be done when we can compute *H*_*θ*_(*x*) in closed form (as is the case for Gaussians, Laplacians, *t*-distributions, and other common likelihoods) or approximate it sufficiently well.

A general approach to approximating Eq. (2) is to draw *θ*_1_, …, *θ*_*m*_∼ *π*_*n*_ and 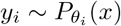 for *i* = 1, …, *m*, and use the estimate

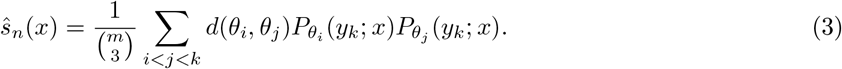

Although Eq. (3) is an unbiased estimator, it may have large variance due to the sampling 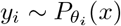. In some cases, we can reduce this variance by avoiding sampling the *y*_*i*_’s whenever we can compute the function

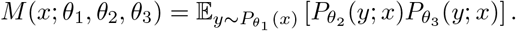

This allows us to compute the alternate approximation

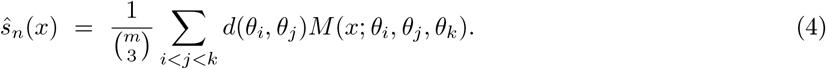

Computing the sums in Eq. (3) or Eq. (4) would take *O*(*m*^3^) time. In practice, we approximate these sums via Monte Carlo by subsampling *N*_mc_ triples (*i, j, k*). Thus, to compute our approximation of Eq. (1) for a set of *B* potential queries takes time *O*(*BN*_mc_). Algorithm 1 presents the full pdbal selection procedure.

The following proposition shows that for the important case of Gaussian likelihoods, we can indeed compute the function *M* in closed form.

##### Proposition 1.

*Fix d ≥* 1, *and let µ*_*i*_ ∈ ℝ^*d*^ *and* 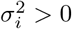 *for i* = 1, 2, 3.

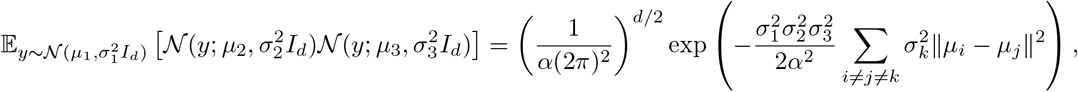

*where* 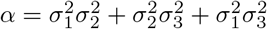.

All proofs are deferred to Appendix B.4.

### B.3 Theoretical results on pdbal

For the purposes of this section, we will assume that all models induce the same entropy for a given *x*.

#### Assumption 1.

*Given any x* ∈ *𝒳, there is a value H*(*x*) *such that H*_*θ*_(*x*) = *H*(*x*) *for all θ* ∈ Θ.

#### Algorithm 1: pdbal selection

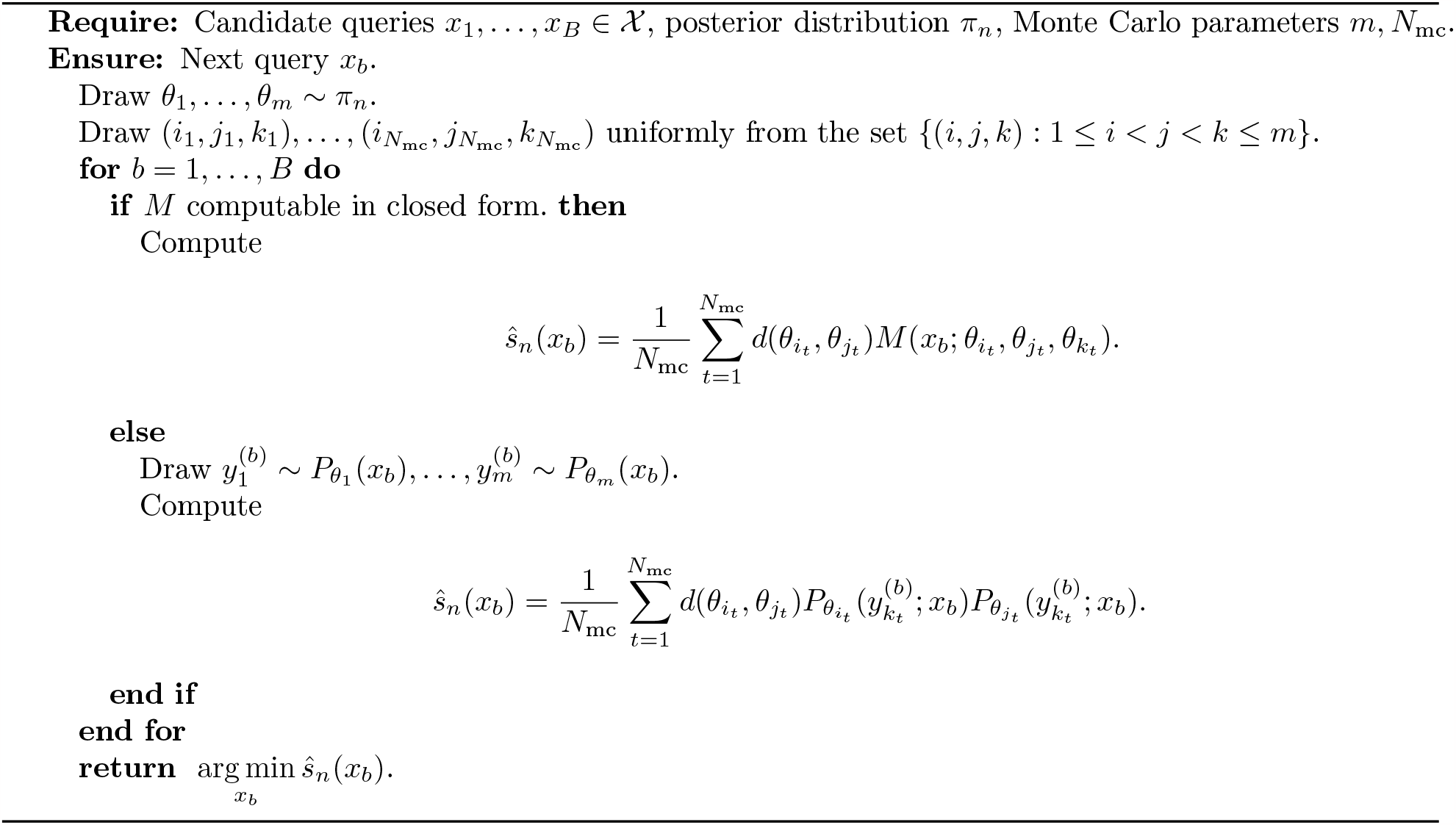

Assumption 1 is satisfied, for example, whenever *P*_*θ*_ is a location model whose scale component is fixed or otherwise assumed to be independent of *θ*. Assumption 1 is not required to implement pdbal, but rather only factors into the analysis in this section.

For a prior *π*, observed data (*x*_1:*n*_, *y*_1:*n*_), and value *ρ* ∈ [0, 1], we say that a data point *x* ∈ *𝒳 ρ-splits* the posterior *π*_*n*_ if

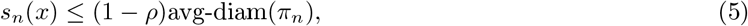

where *s*_*n*_ is the objective function defined in Eq. (1). Intuitively, Eq. (5) captures the notion that there exists a query *x* that shrinks the average diameter by at least (1−*ρ*) in expectation.

We say that *π*_*n*_ is (*ρ, τ* )*-splittable* if

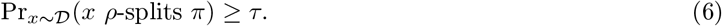

Then for parameters *ρ, ϵ, τ* ∈ (0, 1), we say that Θ (along with corresponding marginal distribution 𝒟 ) has *splitting index* (*ρ, ϵ, τ* ) if for every posterior *π*_*n*_ over Θ satisfying avg-diam(*π*_*n*_) *> ϵ, π*_*n*_ is (*ρ, τ* )-splittable. The definition of splitting in Eq. (5) is similar to those provided in previous diameter-based active learning works ^24,25^. It corresponds to a requirement that posteriors which are not too concentrated (avg-diam *> ϵ*) should have a reasonable number of good queries (at least *τ* % if sampled from 𝒟 ).

One notable difference in this definition is the entropy term inside *s*_*n*_(*x*) in Eq. (5). Without this term, a query with a noisy likelihood will be given higher saliency simply because it produces a wide range of possible outcomes. The entropy term balances out this bias by penalizing queries with a low signal-to-noise ratio.

A key observation in our analysis is that if we query a point that *ρ*-splits the current posterior *π*_*t*_, then in expectation a certain potential function will decrease.

#### Lemma 2.

*If we query a point x*_*t*+1_ *that ρ-splits π*_*t*_, *then*

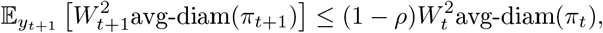

*where* 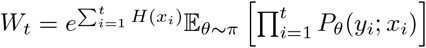.

#### B.3.1 Terminology

In order to prove bounds on the performance of pdbal, we will need to make some assumptions about the complexity of the class Θ and the rates at which empirical entropies within this class concentrate. For a sequence of data pairs *ω*_*n*_ = ((*x*_1_, *y*_1_), …, (*x*_*n*_, *y*_*n*_)), let 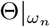 denote the projection of Θ onto *ω*_*n*_. That is:

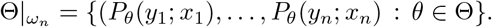

For a sequence *ω*_*n*_ and parameter *ϵ >* 0, define 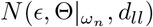 as the size of the minimum cover of 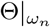 with respect to the distance

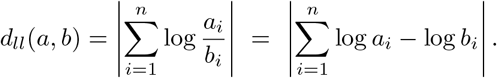

Here, we consider 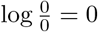. The uniform covering number *N*_*ll*_(*ϵ*, Θ, *n*) is given by

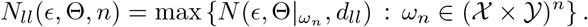

We say that a class Θ is has log-likelihood dimension 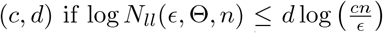for *n ≥ d*. The definition of *N* _*ll*_ (*ϵ*, Θ, *n*) is exactly the same as other uniform covering numbers^76^, modulo the non-standard distance. In the language of statistical learning theory, the log-likelihood dimension is a bound on the *metric entropy* log *N*_*ll*_(*ϵ*, Θ, *n*). We will assume that the log-likelihood dimension is bounded.

##### Assumption 2.

Θ *has log-likelihood dimension* (*c, d*),

As an example of a class with bounded log-likehood dimension, the following result shows how the complexity of a function class translates to the log-likehood dimension of the corresponding Gaussian location model.

##### Proposition 3.

*Fix σ*^2^, *d, B >* 0. *Let* Θ *denote the class of Gaussian location models parameterized by a set of mean functions ℱ*⊂{*θ* : *𝒳* [−*B, B*] *such that P*_*θ*_(*y*; *x*) =𝒩 (*y*|*θ*(*x*), *σ*^2^). *If the responses y lie in* [−*B, B*] *and the pseudo-dimension of ℱ is bounded by d, then* Θ *has log-likelihood dimension* (*cB*^2^*/σ*^2^, *d*) *for some universal constant c >* 0.

Recall a mean-zero random variable *X* is *sub-Gamma* with variance factor *v >* 0 and scale parameter *c >* 0 if

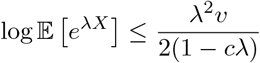

for all *λ* ∈ (0, 1*/c*). We say that the class Θ is *entropy sub-Gamma* with variance factor *v >* 0 and scale parameter *c >* 0 if the random variable 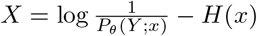 is sub-Gaussian with variance factor *v >* 0 for all *θ* ∈ Θ and *x* ∈ *𝒳*, where *Y* ∼ *P*_*θ*_(*x*). For our bounds we will need Θ to be entropy sub-Gamma.

##### Assumption 3.

Θ *is entropy sub-Gamma with variance v and scale c*^*′*^,

Many likelihoods satisfy Assumption 3 including Gaussians, as illustrated by the following result.

##### Proposition 4.

*Fix σ*^2^ *>* 0 *and let* Θ *denote the class of Gaussian location models from Proposition 3. Then* Θ *is entropy sub-Gamma with variance factor 1 and scale parameter 1*.

Finally, we will require boundedness of both the entropy and the densities of models in Θ.

##### Assumption 4.

*There are constants c*_1_, *c*_2_ *≥* 0 *such that P*_*θ*_(*y*; *x*) *≤ c*_1_ *and* exp(*H*(*x*)) *≤ c*_2_ *for all θ* ∈ Θ, *x* ∈ *𝒳, and y* ∈ *𝒴*.

#### B.3.2 Upper bounds

Given the terminology above, we have the following guarantee on pdbal.

##### Theorem 5.

*Pick d* ≥ 4 *and suppose Assumptions 1 to 4 hold. If at every round t we make a query that ρ-splits π*_*t*_ *and terminate when* avg-diam(*π*_*t*_) ≤ *ϵ, then with probability* 1−*δ*, pdbal *terminates after fewer than*

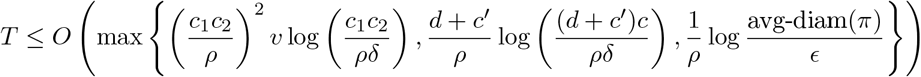

*queries with a posterior satisfying* avg-diam(*π*_*T*_ ) *≤ ϵ*.

As discussed above, Assumptions 1 to 4 are conditions on the complexity and form of Θ. The requirement on the behavior of pdbal can be guaranteed with high probability across all rounds *t* with enough unlabeled data and a fine enough Monte Carlo approximation of Eq. (1).

Given Lemma 2, the proof of Theorem 5 takes two steps

1. Showing that 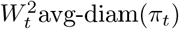 must decrease exponentially quickly.
2. Showing that *W*_*t*_ cannot decrease too quickly.

The only way that both of these can hold is if avg-diam(*π*_*t*_) must also decrease quickly, proving Theorem 5.The first step, showing that 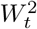avg-diam(*π*_*t*_) decreases quickly, is formalized by the following lemma.

##### Lemma 6.

*Suppose Assumption 4 holds. For any t ≥* 1 *and δ >* 0, *if x*_*i*_ *ρ-splits π*_*i*−1_ *for i* = 1, …, *t, then with probability at least* 1 − *δ*,

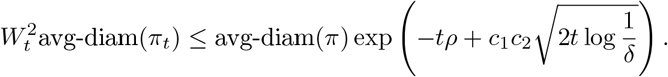

The second step, showing that *W*_*t*_ does not decrease too quickly, is formalized by the following lemma.

##### Lemma 7.

*Suppose Assumption 2 and Assumption 3 hold. If θ*^⋆^ ∼ *π and* 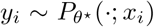*for i* = 1, …, *t, then with probability at least* 1 − *δ*

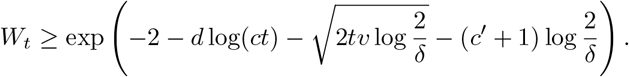

#### B.3.3 Lower bounds

We now turn to showing that in some cases, any optimal active learning strategy must have some dependence on the splitting index of the class. Our first result along these lines is in the deterministic setting.

##### Theorem 8.

*Let* Θ *denote a class of deterministic models that is not* (*ρ, ϵ, τ* )*-splittable for some ρ, ϵ* ∈ (0, 1/4) *and τ* ∈ (0, 1/2). *Let π be any prior distribution satisfying* avg-diam(*π*) *≥* 4*ϵ which is not* (*ρ, τ* )*-splittable. Then any active learning strategy that, with probability at least* 5/6 *(over the random samples from 𝒟 and the observed responses), finds a posterior distribution satisfying* avg-diam(*π*_*t*_) *≤ ϵ must either observe at least* 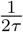 *unlabeled data points or make at least* 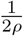*queries*.

We can relax the constraint that our models are deterministic to the case where they have bounded entropy at the expense of a slightly weaker lower bound.

##### Theorem 9.

*Let* Θ *denote a class of models that is not* (*ρ, ϵ, τ* )*-splittable for some ρ, ϵ* ∈ (0, 1/4) *and τ* ∈ (0, 1/2) *such that H*(*x*) = *h < ρ*^3/2^/6 *and P*_*θ*_(*y*; *x*) *≤* 1 *for all x* ∈ *𝒳*., *θ* ∈ Θ *and y* ∈ *𝒴. Let π be any prior distribution satisfying* avg-diam(*π*) *≥* 4*ϵ which is not* (*ρ, τ* )*-splittable. Then any active learning strategy that, with probability at least* 5/6 *(over the random samples from 𝒟 and the observed responses), finds a posterior distribution satisfying* avg-diam(*π*_*t*_) *≤ ϵ must either observe at least* 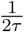 *unlabeled data points or make at least* 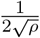 *queries*.

The key ingredient to proving these lower bounds is demonstrating that splitting values are (approximately) sub-additive.

##### Lemma 10.

*Let ρ*_1_, *ρ*_2_ *satisfy ρ*_1_ + *ρ*_2_ *<* 1. *Suppose x*_1_ *ρ*_1_*-splits π, x*_2_ *ρ*_2_*-splits π, and H*(*x*_1_) = *H*(*x*_2_) = *h. Then the following holds:*

- *If h* = 0, *then the combined query* (*x*_1_, *x*_2_) *has splitting value at most ρ*_1_ + *ρ*_2_.
- *If* 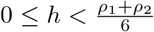, *then the combined query* (*x*_1_, *x*_2_) *has splitting value at most* 2(*ρ*_1_ + *ρ*_2_).

### B.4 Proofs

#### Proof of Proposition 1

From the form of the Gaussian likelihood, we have

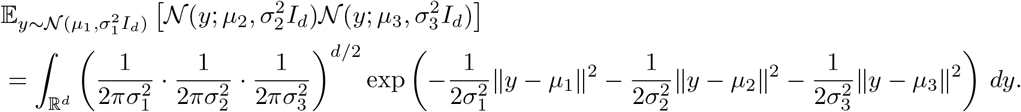

The bias-variance decomposition of squared error implies that for any *c*_1_, …, *c*_*n*_ ≥0 and *b, b*_1_, …, *b*_*n*_ ∈ ℝ ^*d*^, we have

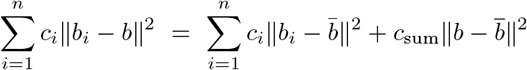

where *c*_sum_ = ∑_*i*_*c*_*i*_ and 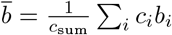. Applying this identity to the exponential above with the notation 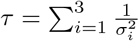 and 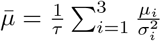, we have

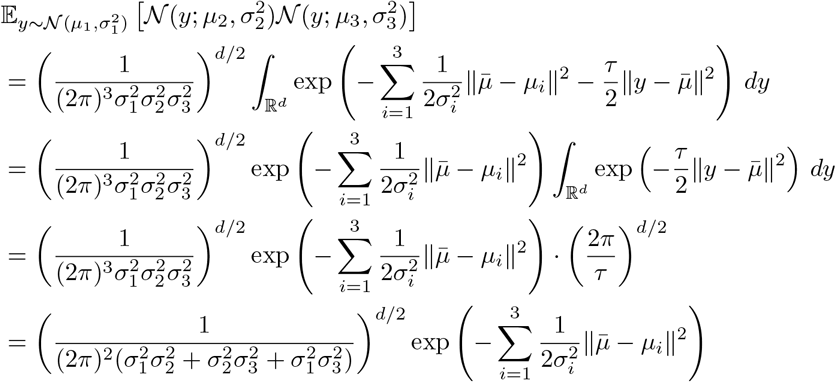

where the second-to-last line follows from the fact that the integral is exactly the normalizing constant of a spherical Gaussian with mean 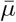 and variance 1*/τ* and the last line follows from expanding the definition of *τ* . Any discrete random variable *X* taking values *x*_*i*_ with probability *p*_*i*_ for *i* = 1, …, *m* satisfies the following:

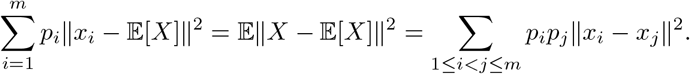

Applying this to the discrete random variable that takes value *µ*_*i*_ with probability 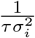,

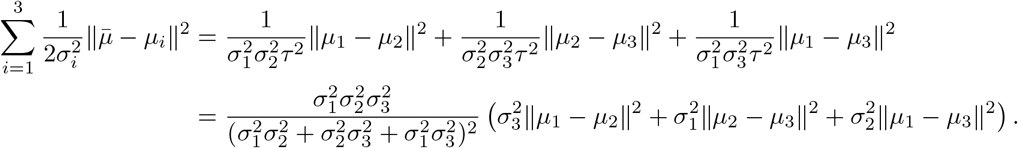

Putting it all together gives us the desired identity. □

#### Proof of Lemma 2

Let *Z*_*t*_ = 𝔼_*θ*∼*π*_[*P* _*θ*_ (*y*_1:t_ ; *x*_1:t_ )], so that we have 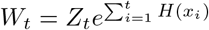. Observe that *Z*_*t*_ is exactly the normalizing constant arising in Bayes’ rule:

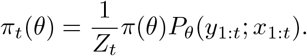

Thus, for any *x*_t+1_, *y*_t+1_, we have

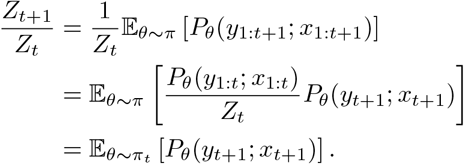

Moreover, by Bayes’s rule, we also have

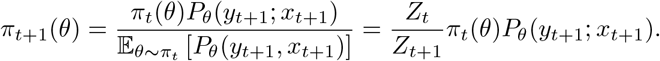

Putting it all together,

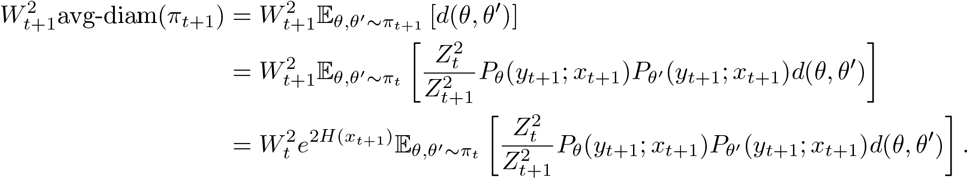

Taking expectations over *y*_t+1_ and applying the definition of splitting finishes the argument.□

#### Proof of Proposition 3

Let (*x*_1_, *y*_1_), …, (*x*_*n*_, *y*_*n*_) be given. For *θ, θ*^*′*^ ∈ Θ,

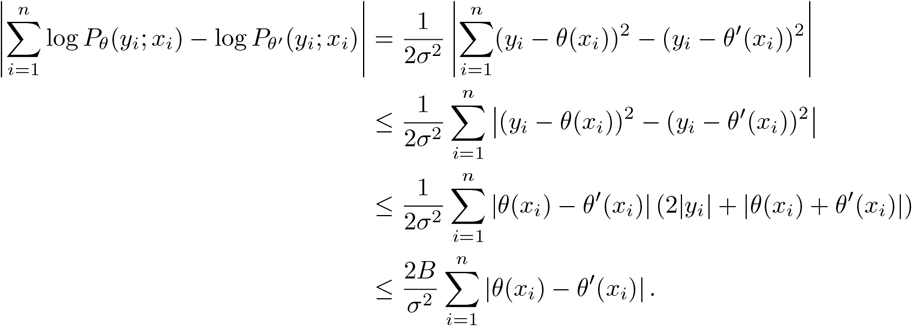

Thus, *N*_*ll*_(*ϵ*, Θ, *n*) ≤*N*_1_(*σ*^2^*ϵ/*2*B*, Θ, *n*), where *N*_1_ denotes uniform covering with respect to *ℓ*_1_ distance. By known bounds on the covering number in terms of the pseudo-dimension, e.g. ^76^ Theorem 18.4, we have

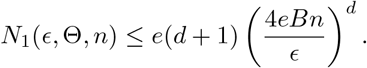

Thus,

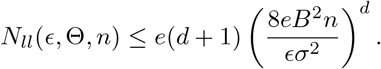

The definition of log-likelihood dimension finishes the argument.□

#### Proof of Proposition 4

For simplicity, let *µ* ∈ ∝ and *σ*^2^ *>* 0. Let *𝒩*(*·*; *µ, σ*^2^) denote the density of a Gaussian with mean *µ* and variance *σ*^2^ . Let 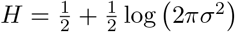denote the entropy of *𝒩*. (· ; *µ, σ*^2^ ).

Suppose *Y* ∼ *𝒩* (*µ, σ* ^2^ ) and 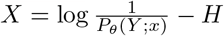, then

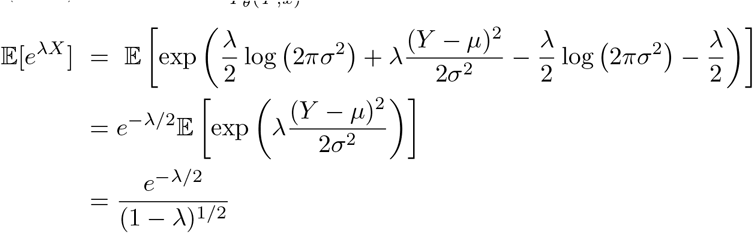

for *λ <* 1. Here, the last line follows from the fact that 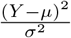 is chi-squared with one degree of freedom, and the known form of the chi-squared moment generating function. Thus, for *λ* ∈ (0, 1), we have

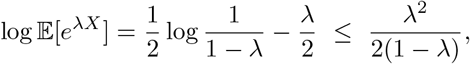

where the inequality follows from the bound log(*x*) ≤ *x*−1 for *x >* 0. Thus, Gaussian location models are entropy sub-Gamma with variance factor 1 and scale parameter 1. □

#### Proof of Lemma 6

For *t ≥* 1, define 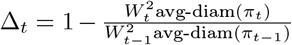 . Let *ℱ*_*t*_ denote the sigma-field of all outcomes up to an including time *t* . Then if we query point *x*_*t*_ which *ρ*-splits *π* _*t*+1_, the definition of splitting implies that

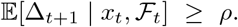

Let 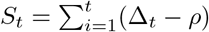 . The above implies that *S*_*t*_ is a submartingale. Moreover, if *P*_*θ*_(*y*; *x*) *≤ c*_1_ uniformly for all *θ, x, y* and *e*^*H*(*x*)^ *≤ c*_2_ for all *x*, then

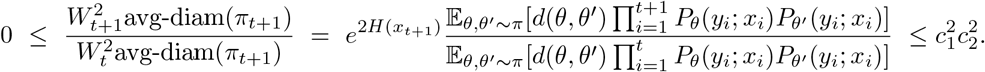

This implies 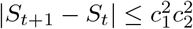, and thus the Azuma-Hoeffding inequality ^77^ tells us that with probability at least 1 − *δ*,

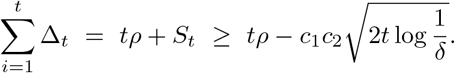

□

#### Proof of Lemma 7

To prove Lemma 7, we will need the following lower bound.

##### Lemma 11.

*Fix ω*_*n*_ = ((*x*_1_, *y*_1_), …, (*x*_*n*_, *y*_*n*_)) ∈ (𝒳 × 𝒴)^*n*^ *and let M be an ϵ-covering of* 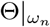 *with respect to d*_*ll*_(·, ·). *Let* Θ_1_, …, Θ_|*M*|_ *be the induced Voronoi partition of* Θ *(breaking ties arbitrarily). Then for any* Θ_*i*_ *and any θ*^⋆^ ∈ Θ_*i*_

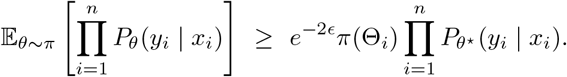

*Proof*. Fix Θ_*i*_ and let *m*_*i*_ ∈ *M* denote the Voronoi ‘center.’ Then for any *θ, θ*^*′*^ ∈ Θ_*i*_, we have

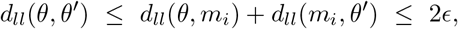

where we have used the fact that *M* is an *ϵ*-cover. Thus, we have

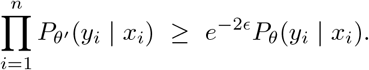

Finally, because Θ_1_, …, Θ_|*M*|_ are disjoint, we have

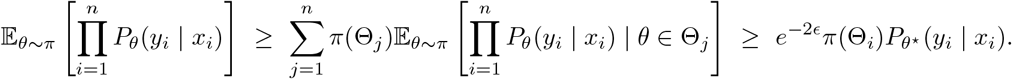

□

We will also require the following result which follows directly from well-known tail bounds for sub-Gamma random variables.

##### Lemma 12.

*Suppose* Θ *is entropy sub-Gamma with variance factor v >* 0 *and scale parameter c >* 0. *If y*_*i*_ ∼ *P*_*θ*_(·; *x*_*i*_) *for i* = 1, …, *t, then with probability at least* 1 − *δ*,

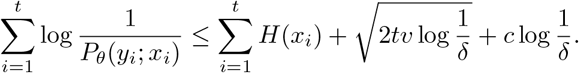

*Proof*. Observe that for independent mean-zero sub-Gamma random variables *X*_1_, …, *X*_*n*_ with variance factor *v >* 0 and scale parameter *c >* 0, the random variable *Z* =∑_*i*_ *X*_*i*_ satisfies

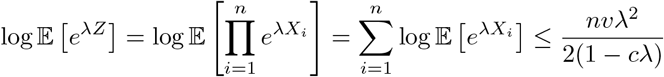

for *λ* ∈ (0, 1*/c*). Thus, *Z* is sub-Gamma with variance factor *nv* and scale parameter *c*. The argument is finished via standard concentration results on sub-Gamma random variables, e.g. ^78^ Chapter 2.4. □

With Lemmas 11 and 12 in hand, we can turn to proving Lemma 7.

*Proof of Lemma 7*. Recall that we can write

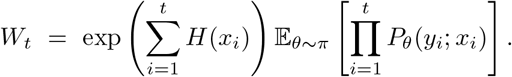

Let *ω*_*t*_ = ((*x*_1_, *y*_1_), …, (*x*_*t*_, *y*_*t*_)) denote our data. By Lemma 11, we have

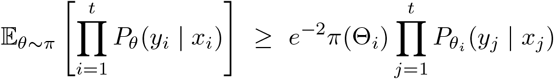

where Θ_1_, …, Θ_*M*_ is the Voronoi partition induced by a minimal 1-covering of 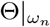, Θ_*i*_ is any of these partition elements, and *θ*_*i*_ is any element in Θ_*i*_. Let *S* denote the set of indices *i* such that *π*(Θ_*i*_) *< δ/*(2*M* ). Then we have

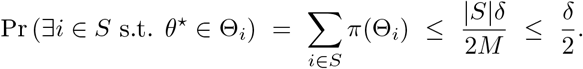

Thus, if Θ^⋆^ is the element of the partition that *θ*^⋆^ falls into, we have with probability at least 1 − *δ/*2

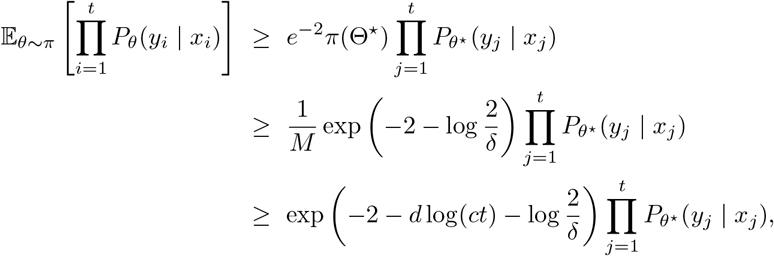

where the last line follows from the fact that *M ≤* (*ct*)^*d*^.

Finally, observe that with probability at least 1 − *δ/*2, Lemma 12 implies

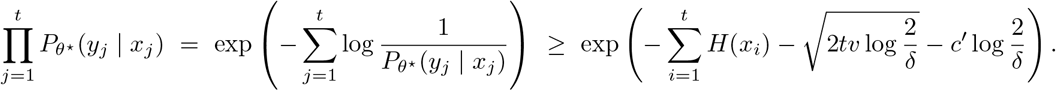

□

A union bound finishes the argument.

#### Proof of Theorem 5

Combining Lemma 6 with a union bound, we have that with probability 1 − *δ/*2

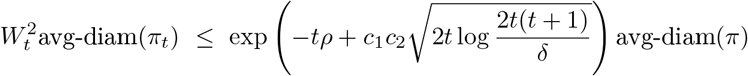

for all *t ≥* 1, simultaneously. Similarly, combining Lemma 7 with a union bound gives us with probability at least 1 − *δ*,

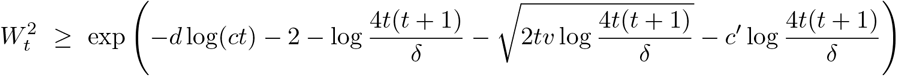

for all *t*≥ 1. Thus, with probability 1−*δ*, both of these occur simultaneously. Plugging in the value of *T* from the theorem statement,

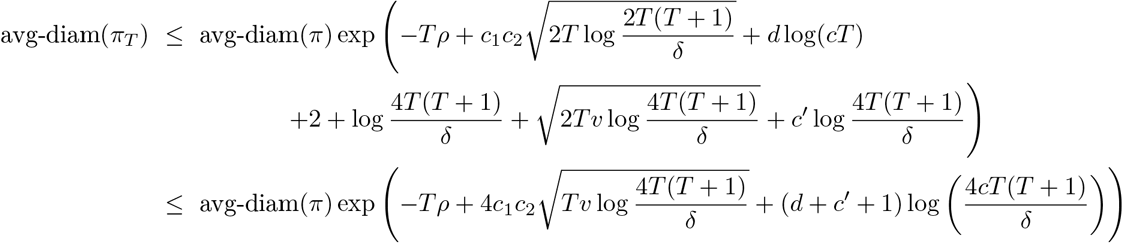

The above is less than *ϵ* when we have

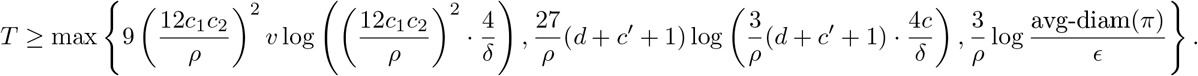

Here, we have made use of the fact that if *a ≥* 1, *b ≥ e* and *x ≥* 9*a* log(*ab*), then *x ≥ a* log(*bx*(*x* + 1)).□

#### Proof of Lemma 10

Observe by the product measure assumption of *P*_*θ*_(·; *x*_1_, *x*_2_), we have

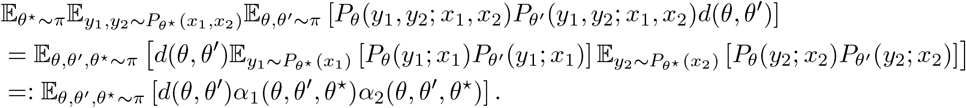

Let *U* be the random variable that takes on value *α*_1_(*θ, θ*^*′*^, *θ*^⋆^) and let *V* denote the random variable that takes on value *α* (*θ, θ*^*′*^, *θ*^⋆^). Here, *θ, θ*^*′*^, *θ*^⋆^ occur with probability 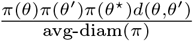. Then it is not hard to see that 𝔼 [*U* ] = (1− *ρ*_1_)*e*^−2*h*^ and 𝔼 [*V* ] = (1− *ρ*_2_)*e*^−2*h*^. Note that *U* and *V* lie in the interval [0, 1] almost surely. Let *A* = 1− *U* and *B* = 1 −*V* .

Let us first consider the case where *h* = 0. Then we have

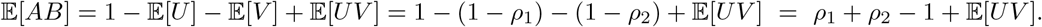

Observe that *AB ≥* 0 almost surely, and so we have

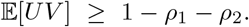

Substituting in our definitions of *U* and *V* gives us the result.

Now consider the case where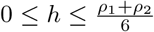 . The same argument as before shows that

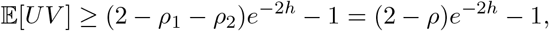

where we have made the substitution *ρ* = *ρ*_1_ + *ρ*_2_. To prove the lemma, we will show that the above is greater than (1 − 2*ρ*)*e*^−4*h*^. This is equivalent to showing

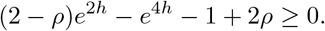

The left-hand side is decreasing for *h ≥* 0. Moreover, we also have the inequality 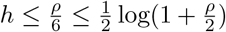. Thus,

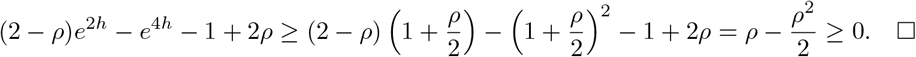

□

#### Proof of Theorem 8

Let *π* be a prior distribution as in the theorem statement. Suppose we draw less than 1/2*τ* unlabeled examples, then with probability at least (1−*τ* )^1/2*τ*^ ≥1/2 none of these *ρ*-split *π*. Let us condition on this event. By induction on Lemma 10, we have that any collection of *n* ≤1*/ρ* of these points does not *nρ*-split *π*.

Suppose that we query *n* of these points (say *x*_1_, …, *x*_*n*_), and receive responses *y*_1_, …, *y*_*n*_. Let *π*_*n*_ denote this posterior. By Lemma 2, we have

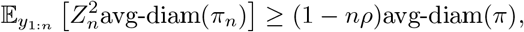

where *Z*_*n*_ = 𝔼_*θ*∼*π*_ [*P*_*θ*_(*y*_1:*n*_; *x*_1:*n*_)] *≤* 1. Thus,

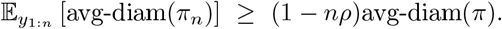

For a random variable *U* satisfying *U ≤ c* almost surely, the reverse Markov inequality gives us

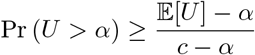

for any *α ≤* 𝔼 [*X*]. Applying this to the random variable 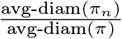 and assuming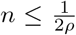, we have that avg-diam(*π*_*n*_) *≥ ϵ* with probability at least 1/3. Putting it all together gives us the theorem statement.□

#### Proof of Theorem 9

We will need the following lemma.

##### Lemma 13.

*Let* 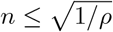, *and let h* ∈ ℝ *satisfy* 0 *≤ h ≤. Suppose x*_1_, …, *x*_*n*_ *all satisfy H*(*x*_*i*_) = *h ≤ ρ/*6*n and have splitting index ≤ ρ. Then the combined query x*_1:*n*_ *has splitting index less than n*^2^*ρ*.

*Proof*. We will show the claim for *n* a power of 2. Extending to other integers is straightforward.

The proof is by induction. Where we first observe that any subsequence *i*_1_, *i*_2_, …, *i*_*k*_ ∈ {1, …, *n}* satisfies that

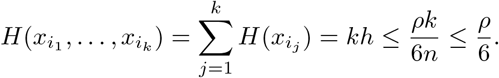

Now for *n* = 1, then the claim trivially holds. For *n ≥* 2, observe that by our inductive hypothesis, we have *x* _1:*n/*2_ and *xn/*2 + 1 : *n* each have splitting index less than 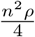 . Applying Lemma 10, completes the argument.□

Turning to the proof of Theorem 9, let *π* be a prior distribution as in the theorem statement. Suppose we draw less than 1/2*τ* unlabeled examples, then with probability at least (1 − *τ* )^1/2*τ*^ *≥*1/2 none of these *ρ*-split *π*. Let us condition on this event.

Now let 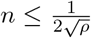 . Using the fact that 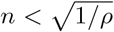 and *h < ρ*^3/2^/6, we can apply Lemma 13, to see that any collection of *n* of these points does not *n*^2^*ρ*-split *π*. Lemma 2 then implies that

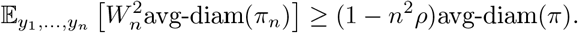

Recall 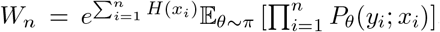. By our assumptions that 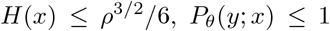, 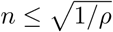 and *ρ ≤* 1/4, we have *W*_*n*_ *≤* 3/2 almost surely. Thus,

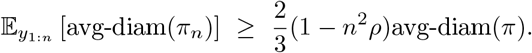

For a random variable *U* satisfying *U ≤ c* almost surely, the reverse Markov inequality gives us

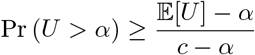

for any *α ≤* 𝔼 [*X*]. Applying this to the random variable 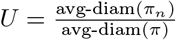 and threshold 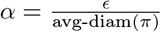, we have

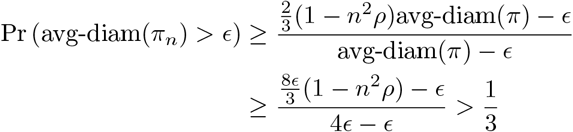

where we have used the fact that 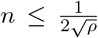 and 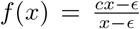 is increasing in *x* when *c* ∈ (0, 1). Thus, (*π*_*n*_) *≥ ϵ* with probability at least 1/3, finishing the argument. □

## Footnotes

1 ALMANAC: https://dtp.cancer.gov/ncialmanac

2 GDSC^2^: https://gdsc-combinations.depmap.sanger.ac.uk/

3 MERCK: https://aacrjournals.org/mct/article/15/6/1155/92159/

4 https://cog.sanger.ac.uk/cancerrxgene/GDSC_release8.4/screened_compounds_rel_8.4.csv

5 https://depmap.org/portal

## Notes

### Competing Interest Statement

SpringWorks Therapeutics [JW]
Emendo Biotherapeutic, Karyopharm Therapeutic, Imago BioSciences, DarwinHealth, Isabl, Labcorp [ALK]
Eisai, Y-mAbs [FSDC]

### Summary of Updates

Fixed some typos in figure captions.

## References

[1] François Clavel and Allan J Hance. HIV drug resistance. New England Journal of Medicine, 350(10):1023–1035, 2004.

[2] Mike Tyers and Gerard D Wright. Drug combinations: A strategy to extend the life of antibiotics in the 21st century. Nature Reviews Microbiology, 17(3):141–155, 2019.

[3] Marilyne Labrie, Joan S Brugge, Gordon B Mills, and Ioannis K Zervantonakis. Therapy resistance: Opportunities created by adaptive responses to targeted therapies in cancer. Nature Reviews Cancer, 22(6):323–339, 2022.

[4] Haojie Jin, Liqin Wang, and René Bernards. Rational combinations of targeted cancer therapies: Background, advances and challenges. Nature Reviews Drug Discovery, 22(3):213–234, 2023.

[5] Susan L Holbeck, Richard Camalier, James A Crowell, Jeevan Prasaad Govindharajulu, Melinda Hollingshead, Lawrence W Anderson, Eric Polley, Larry Rubinstein, Apurva Srivastava, Deborah Wilsker, et al. The National Cancer Institute ALMANAC: A comprehensive screening resource for the detection of anticancer drug pairs with enhanced therapeutic activity. Cancer Research, 77(13):3564–3576, 2017.

[6] Patricia Jaaks, Elizabeth A Coker, Daniel J Vis, Olivia Edwards, Emma F Carpenter, Simonetta M Leto, Lisa Dwane, Francesco Sassi, Howard Lightfoot, Syd Barthorpe, et al. Effective drug combinations in breast, colon and pancreatic cancer cells. Nature, 603(7899):166–173, 2022.

[7] Jennifer O’Neil, Yair Benita, Igor Feldman, Melissa Chenard, Brian Roberts, Yaping Liu, Jing Li, Astrid Kral, Serguei Lejnine, Andrey Loboda, et al. An unbiased oncology compound screen to identify novel combination strategies. Molecular cancer therapeutics, 15(6):1155–1162, 2016.

[8] Jan Wildenhain, Michaela Spitzer, Sonam Dolma, Nick Jarvik, Rachel White, Marcia Roy, Emma Griffiths, David S Bellows, Gerard D Wright, and Mike Tyers. Prediction of synergism from chemical-genetic interactions by machine learning. Cell Systems, 1(6):383–395, 2015.

[9] Peng Li, Chao Huang, Yingxue Fu, Jinan Wang, Ziyin Wu, Jinlong Ru, Chunli Zheng, Zihu Guo, Xuetong Chen, Wei Zhou, et al. Large-scale exploration and analysis of drug combinations. Bioinformatics, 31(12):2007–2016, 2015.

[10] Kristina Preuer, Richard P I Lewis, Sepp Hochreiter, Andreas Bender, Krishna C Bulusu, and Günter Klambauer. DeepSynergy: Predicting anti-cancer drug synergy with Deep Learning. Bioinformatics, 34(9):1538–1546, 12 2017. doi: 10.1093/bioinformatics/btx806.

[11] Qiao Liu and Lei Xie. Transynergy: Mechanism-driven interpretable deep neural network for the synergistic prediction and pathway deconvolution of drug combinations. PLoS Computational Biology, 17(2):e1008653, 2021.

[12] Lianlian Wu, Jie Gao, Yixin Zhang, Binsheng Sui, Yuqi Wen, Qingqiang Wu, Kunhong Liu, Song He, and Xiaochen Bo. A hybrid deep forest-based method for predicting synergistic drug combinations. Cell Reports Methods, 3(2), 2023.

[13] Elizabeth G Ryan, Christopher C Drovandi, James M McGree, and Anthony N Pettitt. A review of modern computational algorithms for Bayesian optimal design. International Statistical Review, 84(1):128–154, 2016.

[14] Burr Settles. Active learning literature survey. Technical Report TR1648, University of Wisconsin-Madison Department of Computer Sciences, 2009.

[15] José Jiménez-Luna, Francesca Grisoni, and Gisbert Schneider. Drug discovery with explainable artificial intelligence. Nature Machine Intelligence, 2(10):573–584, 2020.

[16] David E Graff, Eugene I Shakhnovich, and Connor W Coley. Accelerating high-throughput virtual screening through molecular pool-based active learning. Chemical Science, 12(22):7866–7881, 2021.

[17] Ying Yang, Kun Yao, Matthew P Repasky, Karl Leswing, Robert Abel, Brian K Shoichet, and Steven V Jerome. Efficient exploration of chemical space with docking and deep learning. Journal of Chemical Theory and Computation, 17(11):7106–7119, 2021.

[18] David E Graff, Matteo Aldeghi, Joseph A Morrone, Kirk E Jordan, Edward O Pyzer-Knapp, and Connor W Coley. Self-focusing virtual screening with active design space pruning. Journal of Chemical Information and Modeling, 62(16):3854–3862, 2022.

[19] Paul Bertin, Jarrid Rector-Brooks, Deepak Sharma, Thomas Gaudelet, Andrew Anighoro, Torsten Gross, Francisco Martínez-Peña, Eileen L Tang, MS Suraj, Cristian Regep, et al. RECOVER identifies synergistic drug combinations in vitro through sequential model optimization. Cell Reports Methods, 3(10), 2023.

[20] Adam C Palmer and Peter K Sorger. Combination cancer therapy can confer benefit via patient-to-patient variability without drug additivity or synergy. Cell, 171(7):1678–1691, 2017.

[21] Adam C Palmer, Christopher Chidley, and Peter K Sorger. A curative combination cancer therapy achieves high fractional cell killing through low cross-resistance and drug additivity. eLife, 8:e50036, 2019.

[22] Liang Chang, Paloma Ruiz, Takahiro Ito, and William R Sellers. Targeting pan-essential genes in cancer: challenges and opportunities. Cancer Cell, 39(4):466–479, 2021.

[23] Sarah C. Patterson, Amy E. Pomeroy, and Adam C. Palmer. Ultrasensitive response explains the benefit of combination chemotherapy despite drug antagonism. bioRxiv, 2023. doi: 10.1101/2023.02.27.530263. URL https://www.biorxiv.org/content/early/2023/02/27/2023.02.27.530263.

[24] Christopher Tosh and Sanjoy Dasgupta. Diameter-based active learning. In International Conference on Machine Learning, pages 3444–3452, 2017.

[25] Christopher Tosh and Daniel Hsu. Diameter-based interactive structure discovery. In International Conference on Artificial Intelligence and Statistics, pages 580–590, 2020.

[26] Ronald Aylmer Fisher. Design of experiments. British Medical Journal, 1(3923):554, 1936.

[27] Lei Huang, Fuhai Li, Jianting Sheng, Xiaofeng Xia, Jinwen Ma, Ming Zhan, and Stephen TC Wong. Drug-ComboRanker: Drug combination discovery based on target network analysis. Bioinformatics, 30(12):i228–i236, 2014.

[28] Heli Julkunen, Anna Cichonska, Prson Gautam, Sandor Szedmak, Jane Douat, Tapio Pahikkala, Tero Aittokallio, and Juho Rousu. Leveraging multi-way interactions for systematic prediction of pre-clinical drug combination effects. Nature Communications, 11(1):6136, 2020. doi: 10.1038/s41467-020-19950-z.

[29] Mohammad Lotfollahi, Anna Klimovskaia Susmelj, Carlo De Donno, Leon Hetzel, Yuge Ji, Ignacio L Ibarra, Sanjay R Srivatsan, Mohsen Naghipourfar, Riza M Daza, Beth Martin, Jay Shendure, Jose L McFaline-Figueroa, Pierre Boyeau, F Alexander Wolf, Nafissa Yakubova, Stephan Günnemann, Cole Trapnell, David Lopez-Paz, and Fabian J Theis. Predicting cellular responses to complex perturbations in high-throughput screens. Molecular Systems Biology, page e11517, 2023. doi: 10.15252/msb.202211517.

[30] Anirban Bhattacharya and David B Dunson. Sparse Bayesian infinite factor models. Biometrika, 98(2):291–306, 2011.

[31] Robert H Shoemaker. The nci60 human tumour cell line anticancer drug screen. Nature Reviews Cancer, 6(10): 813–823, 2006.

[32] Liang Chang, Paloma Ruiz, Takahiro Ito, and William R Sellers. Targeting pan-essential genes in cancer: challenges and opportunities. Cancer Cell, 39(4):466–479, 2021.

[33] Aaron Weiss, Jonathan Gill, John Goldberg, Joanne Lagmay, Holly Spraker-Perlman, Rajkumar Venkatramani, and Damon Reed. Advances in therapy for pediatric sarcomas. Current Oncology Reports, 16(8):395, 2014.

[34] David S. Shulman, Sarah B. Whittle, Didier Surdez, Kelly M. Bailey, Enrique de Álava, Jason T. Yustein, Adam Shlien, Masanori Hayashi, Alexander J. R. Bishop, Brian D. Crompton, Steven G. DuBois, Neerav Shukla, Patrick J. Leavey, Stephen L. Lessnick, Heinrich Kovar, Olivier Delattre, Thomas G. P. Grünewald, Cristina R. Antonescu, Ryan D. Roberts, Jeffrey A. Toretsky, Franck Tirode, Richard Gorlick, Katherine A. Janeway, Damon Reed, Elizabeth R. Lawlor, and Patrick J. Grohar. An international working group consensus report for the prioritization of molecular biomarkers for Ewing sarcoma. npj Precision Oncology, 6(1):65, 2022. doi: 10.1038/s41698-022-00307-2.

[35] Leo Kager, Gevorg Tamamyan, and Stefan Bielack. Novel insights and therapeutic interventions for pediatric osteosarcoma. Future Oncology, 13(4):357–368, 2017.

[36] Kelly Bailey, Carrye Cost, Ian Davis, Julia Glade-Bender, Patrick Grohar, Peter Houghton, Michael Isakoff, Elizabeth Stewart, Nadia Laack, Jason Yustein, Damon Reed, Katherine Janeway, Richard Gorlick, Stephen Lessnick, Steven DuBois, and Pooja Hingorani. Emerging novel agents for patients with advanced Ewing sarcoma: A report from the children’s oncology group (COG) new agents for Ewing sarcoma task force. F1000Res, 8, 2019. doi: 10.12688/f1000research.18139.1.

[37] Dorian Yarih Garcia-Ortega, Sara Aileen Cabrera-Nieto, Haydee Sarai Caro-Sánchez, and Marlid Cruz-Ramos. An overview of resistance to chemotherapy in osteosarcoma and future perspectives. Cancer Drug Resistance, 5 (3):762, 2022.

[38] Maya Kansara, Michele W Teng, Mark J Smyth, and David M Thomas. Translational biology of osteosarcoma. Nature Reviews Cancer, 14(11):722–735, 2014.

[39] J Chad Brenner, Felix Y Feng, Sumin Han, Sonam Patel, Siddharth V Goyal, Laura M Bou-Maroun, Meilan Liu, Robert Lonigro, John R Prensner, Scott A Tomlins, and Arul M Chinnaiyan. PARP-1 inhibition as a targeted strategy to treat Ewing’s sarcoma. Cancer Res, 72(7):1608–1613, Apr 2012. doi: 10.1158/0008-5472.CAN-11-3648.

[40] Elizabeth Stewart, Ross Goshorn, Cori Bradley, Lyra M Griffiths, Claudia Benavente, Nathaniel R Twarog, Gregory M Miller, William Caufield, Burgess B 3rd Freeman, Armita Bahrami, Alberto Pappo, Jianrong Wu, Amos Loh, Åsa Karlström, Chris Calabrese, Brittney Gordon, Lyudmila Tsurkan, M Jason Hatfield, Philip M Potter, Scott E Snyder, Suresh Thiagarajan, Abbas Shirinifard, Andras Sablauer, Anang A Shelat, and Michael A Dyer. Targeting the DNA repair pathway in Ewing sarcoma. Cell Rep, 9(3):829–841, Nov 2014. doi: 10.1016/j.celrep.2014.09.028.

[41] Sara M. Federico, Alberto S. Pappo, Natasha Sahr, April Sykes, Olivia Campagne, Clinton F. Stewart, Michael R. Clay, Armita Bahrami, Mary B. McCarville, Sue C. Kaste, Victor M. Santana, Sara Helmig, Jessica Gartrell, Anang Shelat, Rachel C. Brennan, Dana Hawkins, Kimberly Godwin, Michael W. Bishop, Wayne L. Furman, and Elizabeth Stewart. A phase I trial of talazoparib and irinotecan with and without temozolomide in children and young adults with recurrent or refractory solid malignancies. European Journal of Cancer, 137:204–213, 2020.

[42] Rashmi Chugh, Karla V Ballman, Lee J Helman, Shreyaskumar Patel, Jeremy S Whelan, Brigitte Widemann, Yao Lu, Douglas S Hawkins, Leo Mascarenhas, John W Glod, Jiuping Ji, Yiping Zhang, Denise Reinke, and Sandra J Strauss. SARC025 arms 1 and 2: A phase 1 study of the poly(ADP-ribose) polymerase inhibitor niraparib with temozolomide or irinotecan in patients with advanced Ewing sarcoma. Cancer, 127(8):1301–1310, Apr 2021.

[43] Stefan K. Zöllner, James F. Amatruda, Sebastian Bauer, Stéphane Collaud, Enrique de Álava, Steven G. DuBois, Jendrik Hardes, Wolfgang Hartmann, Heinrich Kovar, Markus Metzler, David S. Shulman, Arne Streitbürger, Beate Timmermann, Jeffrey A. Toretsky, Yasmin Uhlenbruch, Volker Vieth, Thomas G. P. Grünewald, and Uta Dirksen. Ewing sarcoma—Diagnosis, treatment, clinical challenges and future perspectives. Journal of Clinical Medicine, 10(8), 2021. doi: 10.3390/jcm10081685.

[44] Kimberly Stegmaier, Jenny S Wong, Kenneth N Ross, Kwan T Chow, David Peck, Renee D Wright, Stephen L Lessnick, Andrew L Kung, and Todd R Golub. Signature-based small molecule screening identifies cytosine arabinoside as an EWS/FLI modulator in Ewing sarcoma. PLoS Med, 4(4):e122, Apr 2007.

[45] Sudan N Loganathan, Nan Tang, Jonathan T Fleming, Yufang Ma, Yan Guo, Scott C Borinstein, Chin Chiang, and Jialiang Wang. BET bromodomain inhibitors suppress EWS-FLI1-dependent transcription and the IGF1 autocrine mechanism in Ewing sarcoma. Oncotarget, 7(28):43504–43517, Jul 2016. doi: 10.18632/oncotarget.9762.

[46] Paradesi Naidu Gollavilli, Aishwarya Pawar, Kari Wilder-Romans, Ramakrishnan Natesan, Carl G. Engelke, Vijaya L. Dommeti, Pranathi M. Krishnamurthy, Archana Nallasivam, Ingrid J. Apel, Tianlei Xu, Zhaohui S. Qin, Felix Y. Feng, and Irfan A. Asangani. EWS/ETS-driven Ewing sarcoma requires BET bromodomain proteins. Cancer Research, 78(16):4760–4773, 08 2018. doi: 10.1158/0008-5472.CAN-18-0484.

[47] Peter L. Bonate, Larry Arthaud, William R. Cantrell, Katherine Stephenson, John A. Secrist, and Steve Weitman. Discovery and development of clofarabine: A nucleoside analogue for treating cancer. Nature Reviews Drug Discovery, 5(10):855–863, 2006.

[48] Thomas P Leist and Robert Weissert. Cladribine: Mode of action and implications for treatment of multiple sclerosis. Clinical neuropharmacology, 34(1):28–35, 2011.

[49] Haydar Çelik, Marika Sciandra, Bess Flashner, Elif Gelmez, Neslihan Kayraklioglu, David V. Allegakoen, Jeff R. Petro, Erin J. Conn, Sarah Hour, Jenny Han, Lalehan Oktay, Purushottam B. Tiwari, Mutlu Hayran, Brent T. Harris, Maria Cristina Manara, Jeffrey A. Toretsky, Katia Scotlandi, and Aykut Üren. Clofarabine inhibits Ewing sarcoma growth through a novel molecular mechanism involving direct binding to CD99. Oncogene, 37(16): 2181–2196, 2018.

[50] F Cidre-Aranaz, T G P Grünewald, D Surdez, L García-García, J Carlos Lázaro, T Kirchner, L González-González, A Sastre, P García-Miguel, S E López-Pérez, S Monzón, O Delattre, and J Alonso. EWS-FLI1-mediated suppression of the RAS-antagonist Sprouty 1 (SPRY1) confers aggressiveness to Ewing sarcoma. Oncogene, 36 (6):766–776, Feb 2017. doi: 10.1038/onc.2016.244.

[51] Peter Norman. Tipifarnib (janssen pharmaceutica). Current Opinion in Investigational Drugs (London, England: 2000), 3(2):313–319, 2002.

[52] José Luis Ordóñez, Ana Teresa Amaral, Angel M Carcaboso, David Herrero-Martín, María del Carmen García-Macías, Vicky Sevillano, Diego Alonso, Guillem Pascual-Pasto, Laura San-Segundo, Monica Vila-Ubach, et al. The PARP inhibitor olaparib enhances the sensitivity of Ewing sarcoma to trabectedin. Oncotarget, 6(22):18875, 2015.

[53] Florian Engert, Cornelius Schneider, Lilly Magdalena Weiß, Marie Probst, and Simone Fulda. PARP inhibitors sensitize Ewing sarcoma cells to temozolomide-induced apoptosis via the mitochondrial pathway. Molecular Cancer Therapeutics, 14(12):2818–2830, 2015.

[54] Sam Behjati, Patrick S. Tarpey, Kerstin Haase, Hongtao Ye, Matthew D. Young, Ludmil B. Alexandrov, Sarah J. Farndon, Grace Collord, David C. Wedge, Inigo Martincorena, Susanna L. Cooke, Helen Davies, William Mifsud, Mathias Lidgren, Sancha Martin, Calli Latimer, Mark Maddison, Adam P. Butler, Jon W. Teague, Nischalan Pillay, Adam Shlien, Ultan McDermott, P. Andrew Futreal, Daniel Baumhoer, Olga Zaikova, Bodil Bjerkehagen, Ola Myklebost, M. Fernanda Amary, Roberto Tirabosco, Peter Van Loo, Michael R. Stratton, Adrienne M. Flanagan, and Peter J. Campbell. Recurrent mutation of igf signalling genes and distinct patterns of genomic rearrangement in osteosarcoma. Nature Communications, 8(1):15936, 2017.

[55] Isidro Cortés-Ciriano, Jake June-Koo Lee, Ruibin Xi, Dhawal Jain, Youngsook L. Jung, Lixing Yang, Dmitry Gordenin, Leszek J. Klimczak, Cheng-Zhong Zhang, David S. Pellman, Kadir C. Akdemir, Eva G. Alvarez, Adrian Baez-Ortega, Rameen Beroukhim, Paul C. Boutros, David D. L. Bowtell, Benedikt Brors, Kathleen H. Burns, Peter J. Campbell, Kin Chan, Ken Chen, Ana Dueso-Barroso, Andrew J. Dunford, Paul A. Edwards, Xavier Estivill, Dariush Etemadmoghadam, Lars Feuerbach, J. Lynn Fink, Milana Frenkel-Morgenstern, Dale W. Garsed, Mark Gerstein, Dmitry A. Gordenin, David Haan, James E. Haber, Julian M. Hess, Barbara Hutter, Marcin Imielinski, David T. W. Jones, Young Seok Ju, Marat D. Kazanov, Leszek J. Klimczak, Youngil Koh, Jan O. Korbel, Kiran Kumar, Eunjung Alice Lee, Yilong Li, Andy G. Lynch, Geoff Macintyre, Florian Markowetz, Iñigo Martincorena, Alexander Martinez-Fundichely, Satoru Miyano, Hidewaki Nakagawa, Fabio C. P. Navarro, Stephan Ossowski, Peter J. Park, John V. Pearson, Montserrat Puiggròs, Karsten Rippe, Nicola D. Roberts, Steven A. Roberts, Bernardo Rodriguez-Martin, Steven E. Schumacher, Ralph Scully, Mark Shackleton, Nikos Sidiropoulos, Lina Sieverling, Chip Stewart, David Torrents, Jose M. C. Tubio, Izar Villasante, Nicola Waddell, Jeremiah A. Wala, Joachim Weischenfeldt, Xiaotong Yao, Sung-Soo Yoon, Jorge Zamora, Peter J. Park, Lauri A. Aaltonen, Federico Abascal, Adam Abeshouse, Hiroyuki Aburatani, David J. Adams, Nishant Agrawal, Keun Soo Ahn, Sung-Min Ahn, Hiroshi Aikata, Rehan Akbani, Kadir C. Akdemir, Hikmat Al-Ahmadie, Sultan T. Al-Sedairy, Fatima Al-Shahrour, Malik Alawi, Monique Albert, Kenneth Aldape, Ludmil B. Alexandrov, Adrian Ally, Kathryn Alsop, Eva G. Alvarez, Fernanda Amary, Samirkumar B. Amin, Brice Aminou, Ole Ammerpohl, Matthew J. Anderson, Yeng Ang, Davide Antonello, Pavana Anur, Samuel Aparicio, Elizabeth L. Appelbaum, Yasuhito Arai, Axel Aretz, Koji Arihiro, Shun-ichi Ariizumi, Joshua Armenia, Laurent Arnould, Sylvia Asa, Yassen Assenov, Gurnit Atwal, Sietse Aukema, J. Todd Auman, Miriam R. R. Aure, Philip Awadalla, Marta Aymerich, Gary D. Bader, Matthew H. Bailey, Peter J. Bailey, Miruna Balasundaram, Saianand Balu, Pratiti Bandopadhayay, Rosamonde E. Banks, Stefano Barbi, Andrew P. Barbour, Jonathan Barenboim, Jill Barnholtz-Sloan, Hugh Barr, Elisabet Barrera, John Bartlett, Javier Bartolome, Claudio Bassi, Oliver F. Bathe, Daniel Baumhoer, Prashant Bavi, Stephen B. Baylin, Wojciech Bazant, Duncan Beardsmore, Timothy A. Beck, Sam Behjati, Andreas Behren, Beifang Niu, Cindy Bell, Sergi Beltran, Christopher Benz, Andrew Berchuck, Anke K. Bergmann, Erik N. Bergstrom, Benjamin P. Berman, Daniel M. Berney, Stephan H. Bernhart, Mario Berrios, Samantha Bersani, Johanna Bertl, Miguel Betancourt, Vinayak Bhandari, Shriram G. Bhosle, Andrew V. Biankin, Matthias Bieg, Darell Bigner, Hans Binder, Ewan Birney, Michael Birrer, Nidhan K. Biswas, Bodil Bjerkehagen, Tom Bodenheimer, Lori Boice, Giada Bonizzato, Johann S. De Bono, Arnoud Boot, Moiz S. Bootwalla, Ake Borg, Arndt Borkhardt, Keith A. Boroevich, Ivan Borozan, Christoph Borst, Marcus Bosenberg, Mattia Bosio, Jacqueline Boultwood, Guillaume Bourque, Paul C. Boutros, G. Steven Bova, David T. Bowen, Reanne Bowlby, David D. L. Bowtell, Sandrine Boyault, Rich Boyce, Jeffrey Boyd, Alvis Brazma, Paul Brennan, Daniel S. Brewer, Arie B. Brinkman, Robert G. Bristow, Russell R. Broaddus, Jane E. Brock, Malcolm Brock, Annegien Broeks, Angela N. Brooks, Denise Brooks, Søren Brunak, Timothy J. C. Bruxner, Alicia L. Bruzos, Alex Buchanan, Ivo Buchhalter, Christiane Buchholz, Susan Bullman, Hazel Burke, Birgit Burkhardt, Kathleen H. Burns, John Busanovich, Carlos D. Bustamante, Adam P. Butler, Atul J. Butte, Niall J. Byrne, Anne-Lise Børresen-Dale, Samantha J. Caesar-Johnson, Andy Cafferkey, Declan Cahill, Claudia Calabrese, Carlos Caldas, Fabien Calvo, Niedzica Camacho, Peter J. Campbell, Elias Campo, Cinzia Cantù, Shaolong Cao, Thomas E. Carey, Joana Carlevaro-Fita, Rebecca Carlsen, Ivana Cataldo, Mario Cazzola, Jonathan Cebon, Robert Cerfolio, Dianne E. Chadwick, Dimple Chakravarty, Don Chalmers, Calvin Wing Yiu Chan, Michelle Chan-Seng-Yue, Vishal S. Chandan, David K. Chang, Stephen J. Chanock, Lorraine A. Chantrill, Aurélien Chateigner, Nilanjan Chatterjee, Kazuaki Chayama, Hsiao-Wei Chen, Jieming Chen, Yiwen Chen, Zhaohong Chen, Andrew D. Cherniack, Jeremy Chien, Yoke-Eng Chiew, Suet-Feung Chin, Juok Cho, Sunghoon Cho, Jung Kyoon Choi, Wan Choi, Christine Chomienne, Zechen Chong, Su Pin Choo, Angela Chou, Angelika N. Christ, Elizabeth L. Christie, Eric Chuah, Carrie Cibulskis, Kristian Cibulskis, Sara Cingarlini, Peter Clapham, Alexander Claviez, Sean Cleary, Nicole Cloonan, Marek Cmero, Colin C. Collins, Ashton A. Connor, Susanna L. Cooke, Colin S. Cooper, Leslie Cope, Vincenzo Corbo, Matthew G. Cordes, Stephen M. Cordner, Kyle Covington, PCAWG Structural Variation Working Group, and PCAWG Consortium. Comprehensive analysis of chromothripsis in 2,658 human cancers using whole-genome sequencing. Nature Genetics, 52(3):331–341, 2020.

[56] Mahmoud Ghandi, Franklin W. Huang, Judit Jané-Valbuena, Gregory V. Kryukov, Christopher C. Lo, E. Robert McDonald, Jordi Barretina, Ellen T. Gelfand, Craig M. Bielski, Haoxin Li, Kevin Hu, Alexander Y. Andreev-Drakhlin, Jaegil Kim, Julian M. Hess, Brian J. Haas, François Aguet, Barbara A. Weir, Michael V. Rothberg, Brenton R. Paolella, Michael S. Lawrence, Rehan Akbani, Yiling Lu, Hong L. Tiv, Prafulla C. Gokhale, Antoine de Weck, Ali Amin Mansour, Coyin Oh, Juliann Shih, Kevin Hadi, Yanay Rosen, Jonathan Bistline, Kavitha Venkatesan, Anupama Reddy, Dmitriy Sonkin, Manway Liu, Joseph Lehar, Joshua M. Korn, Dale A. Porter, Michael D. Jones, Javad Golji, Giordano Caponigro, Jordan E. Taylor, Caitlin M. Dunning, Amanda L. Creech, Allison C. Warren, James M. McFarland, Mahdi Zamanighomi, Audrey Kauffmann, Nicolas Stransky, Marcin Imielinski, Yosef E. Maruvka, Andrew D. Cherniack, Aviad Tsherniak, Francisca Vazquez, Jacob D. Jaffe, Andrew A. Lane, David M. Weinstock, Cory M. Johannessen, Michael P. Morrissey, Frank Stegmeier, Robert Schlegel, William C. Hahn, Gad Getz, Gordon B. Mills, Jesse S. Boehm, Todd R. Golub, Levi A. Garraway, and William R. Sellers. Next-generation characterization of the cancer cell line encyclopedia. Nature, 569(7757): 503–508, 2019.

[57] Joshua M. Dempster, Jordan Rossen, Mariya Kazachkova, Joshua Pan, Guillaume Kugener, David E. Root, and Aviad Tsherniak. Extracting biological insights from the project achilles genome-scale crispr screens in cancer cell lines. bioRxiv, page 720243, 01 2019.

[58] Jeff Hirst and Andrew K Godwin. Aurka inhibition mimics brcaness. Aging (Albany NY), 9(9):1945–1946, Sep 2017.

[59] Andrea K Byrum, Alessandro Vindigni, and Nima Mosammaparast. Defining and modulating ‘brcaness’. Trends Cell Biol, 29(9):740–751, Sep 2019.

[60] Florian Engert, Michal Kovac, Daniel Baumhoer, Michaela Nathrath, and Simone Fulda. Osteosarcoma cells with genetic signatures of brcaness are susceptible to the parp inhibitor talazoparib alone or in combination with chemotherapeutics. Oncotarget, 8(30):48794–48806, Jul 2017.

[61] Harriett Holme, Aditi Gulati, Rachel Brough, Emmy D. G. Fleuren, Ilirjana Bajrami, James Campbell, Irene Y. Chong, Sara Costa-Cabral, Richard Elliott, Tim Fenton, Jessica Frankum, Samuel E. Jones, Malini Menon, Rowan Miller, Helen N. Pemberton, Sophie Postel-Vinay, Rumana Rafiq, Joanna L. Selfe, Alex von Kriegsheim, Amaya Garcia Munoz, Javier Rodriguez, Janet Shipley, Winette T. A. van der Graaf, Chris T. Williamson, Colm J. Ryan, Stephen Pettitt, Alan Ashworth, Sandra J. Strauss, and Christopher J. Lord. Chemosensitivity profiling of osteosarcoma tumour cell lines identifies a model of brcaness. Scientific Reports, 8(10614), 2018.

[62] Chaoyang Sun, Yong Fang, Jun Yin, Jian Chen, Zhenlin Ju, Dong Zhang, Xiaohua Chen, Christopher P Vellano, Kang Jin Jeong, Patrick Kwok-Shing Ng, Agda Karina B Eterovic, Neil H Bhola, Yiling Lu, Shannon N Westin, Jennifer R Grandis, Shiaw-Yih Lin, Kenneth L Scott, Guang Peng, Joan Brugge, and Gordon B Mills. Rational combination therapy with parp and mek inhibitors capitalizes on therapeutic liabilities in ras mutant cancers. Sci Transl Med, 9(392), May 2017.

[63] Francesca Vena, Ruochen Jia, Arman Esfandiari, Juan J Garcia-Gomez, Manuel Rodriguez-Justo, Jianguo Ma, Sakeena Syed, Lindsey Crowley, Brian Elenbaas, Samantha Goodstal, John A Hartley, and Daniel Hochhauser. Mek inhibition leads to brca2 downregulation and sensitization to dna damaging agents in pancreas and ovarian cancer models. Oncotarget, 9(14):11592–11603, Feb 2018.

[64] Sang-Yun Lee, In-Seong Koo, Hyun Ju Hwang, and Dong Woo Lee. In vitro three-dimensional (3D) cell culture tools for spheroid and organoid models. SLAS Discovery, 2023.

[65] Dylan C Mitchell, Miljan Kuljanin, Jiaming Li, Jonathan G Van Vranken, Nathan Bulloch, Devin K Schweppe, Edward L Huttlin, and Steven P Gygi. A proteome-wide atlas of drug mechanism of action. Nature Biotechnology, pages 1–13, 2023.

[66] Hans-Hermann Wessels, Alejandro Méndez-Mancilla, Yuhan Hao, Efthymia Papalexi, William M Mauck III, Lu Lu, John A Morris, Eleni P Mimitou, Peter Smibert, Neville E Sanjana, and Rahul Satija. Efficient combinatorial targeting of rna transcripts in single cells with cas13 rna perturb-seq. Nature Methods, 20(1):86–94, 2023.

[67] Carlos M Carvalho, Nicholas G Polson, and James G Scott. The horseshoe estimator for sparse signals. Biometrika, 97(2):465–480, 2010.

[68] Daniel Golovin and Andreas Krause. Adaptive submodularity: Theory and applications in active learning and stochastic optimization. Journal of Artificial Intelligence Research, 42:427–486, 2011.

[69] Yuxin Chen and Andreas Krause. Near-optimal batch mode active learning and adaptive submodular optimization. In Proceedings of the 30th International Conference on Machine Learning, pages 160–168, 2013.

[70] F. Pedregosa, G. Varoquaux, A. Gramfort, V. Michel, B. Thirion, O. Grisel, M. Blondel, P. Prettenhofer, R. Weiss, V. Dubourg, J. Vanderplas, A. Passos, D. Cournapeau, M. Brucher, M. Perrot, and E. Duchesnay. Scikit-learn: Machine learning in Python. Journal of Machine Learning Research, 12:2825–2830, 2011.

[71] Ville Satopaa, Jeannie Albrecht, David Irwin, and Barath Raghavan. Finding a “kneedle” in a haystack: Detecting knee points in system behavior. In 31st International Conference on Distributed Computing Systems Workshops, pages 166–171, 2011.

[72] Michael L. Waskom. Seaborn: Statistical data visualization. Journal of Open Source Software, 6(60):3021, 2021.

[73] Pauli Virtanen, Ralf Gommers, Travis E. Oliphant, Matt Haberland, Tyler Reddy, David Cournapeau, Evgeni Burovski, Pearu Peterson, Warren Weckesser, Jonathan Bright, Stéfan J. van der Walt, Matthew Brett, Joshua Wilson, K. Jarrod Millman, Nikolay Mayorov, Andrew R. J. Nelson, Eric Jones, Robert Kern, Eric Larson, C J Carey, Ilhan Polat, Yu Feng, Eric W. Moore, Jake VanderPlas, Denis Laxalde, Josef Perktold, Robert Cimrman, Ian Henriksen, E. A. Quintero, Charles R. Harris, Anne M. Archibald, Antônio H. Ribeiro, Fabian Pedregosa, Paul van Mulbregt, and SciPy 1.0 Contributors. SciPy 1.0: Fundamental Algorithms for Scientific Computing in Python. Nature Methods, 17:261–272, 2020.

[74] Donavan T. Cheng, Talia N. Mitchell, Ahmet Zehir, Ronak H. Shah, Ryma Benayed, Aijazuddin Syed, Raghu Chandramohan, Zhen Yu Liu, Helen H. Won, Sasinya N. Scott, A. Rose Brannon, Catherine O’Reilly, Justyna Sadowska, Jacklyn Casanova, Angela Yannes, Jaclyn F. Hechtman, Jinjuan Yao, Wei Song, Dara S. Ross, Alifya Oultache, Snjezana Dogan, Laetitia Borsu, Meera Hameed, Khedoudja Nafa, Maria E. Arcila, Marc Ladanyi, and Michael F. Berger. Memorial sloan kettering-integrated mutation profiling of actionable cancer targets (msk-impact): A hybridization capture-based next-generation sequencing clinical assay for solid tumor molecular oncology. The Journal of Molecular Diagnostics, 17(3):251–264, 2015.

[75] Yoav Benjamini and Yosef Hochberg. Controlling the false discovery rate: A practical and powerful approach to multiple testing. Journal of the Royal Statistical Society: Series B (Methodological), 57(1):289–300, 1995.

[76] Martin Anthony and Peter Bartlett. Neural network learning: Theoretical foundations. Cambridge University Press, 1999.

[77] Kazuoki Azuma. Weighted sums of certain dependent random variables. Tohoku Mathematical Journal, Second Series, 19(3):357–367, 1967.

[78] Stéphane Boucheron, Gábor Lugosi, and Pascal Massart. Concentration inequalities: A nonasymptotic theory of independence. Oxford university press, 2013.

